# Cell fate coordinates mechano-osmotic forces in intestinal crypt morphogenesis

**DOI:** 10.1101/2020.05.13.094359

**Authors:** Qiutan Yang, Shi-Lei Xue, Chii Jou Chan, Markus Rempfler, Dario Vischi, Francisca Mauer Gutierrez, Takashi Hiiragi, Edouard Hannezo, Prisca Liberali

## Abstract

Intestinal organoids derived from single cells undergo complex crypt-villus patterning and morphogenesis. However, the nature and coordination of the underlying forces remains poorly characterized. Through light-sheet microscopy and mechanical perturbations, we demonstrate that organoid crypt formation coincides with stark lumen volume reduction, which works synergistically with actomyosin-generated crypt apical and villus basal tension to drive morphogenesis. We analyse these mechanical features in a quantitative 3D biophysical model and detect a critical point in actomyosin tensions, above which crypt becomes robust to volume changes. Finally, via single-cell RNA sequencing and pharmacological perturbations, we show that enterocyte-specific expressed sodium/glucose cotransporter modulates lumen volume reduction via promoting cell swelling. Altogether, our study reveals how cell fate-specific changes in osmotic and actomyosin forces coordinate robust organoid morphogenesis.

**One Sentence Summary:** Emergence of region-specific cell fates drive actomyosin patterns and luminal osmotic changes in organoid development

---

During development of the mammalian intestine, crypt morphogenesis builds the stem-cell niche compartment, composed of stem cells and Paneth cells, as an essential unit for the homeostasis of the intestinal epithelium (*1*). In both mice and humans, crypts form after the emergence of villi, which include other differentiated cells, such as enterocytes and goblet cells (*2-4*). Mouse villus formation is initiated by the mechanical condensation of the mesenchymal cells underneath the epithelium (*5*), which leads to tissue folding and the establishment of Shh and the BMP morphogen gradients along the villus-crypt axis (*6*). The morphogen gradients induce cell differentiation that further promotes the villus morphogenesis (*6, 7*). While villus morphogenesis has been well studied, less is known about the mechanism of crypt morphogenesis (*4, 6, 8, 9*). Apical constriction of the intra-villi tissue has been shown to initiate the early invagination of the crypts (*10, 11*), however, the mechanisms of crypt formation and especially the coordination between cell type emergence and tissue morphogenesis remains to be investigated.

Major challenges of studying intestinal morphogenesis lie in the difficulty of cellular-level live-imaging and force manipulation tools, because of the limited accessibility of internal organs. *In vitro*, mouse intestinal organoids contain functional crypt and villus regions and provide a unique opportunity to overcome these difficulties (*12-14)*. Mouse intestinal organoids develop from single cells via two sequential symmetry breaking events. The first one is at cell fate level, leading to the emergence of the first Paneth cell between the 8 and 32-cell cyst stages under the regulation of Notch and Hippo pathways (*15*). Paneth cells then induce the Lgr-5^+^ stem cells, after which a small bulge forms around the Paneth and stem cells around Day3 of organoid growth (*15, 16)*. Around Day4, a crypt buds from the bulge in the cyst with a narrow neck, and this shape transformation determines the second symmetry breaking event (Fig. 1A). Mouse intestinal organoids share strong similarities with the *in vivo* tissue including cell type composition, tissue organization and functionalities (*14*). Interestingly, mouse intestinal organoids are one of the few organoid systems that can recapitulate the shape of the tissue and crypt morphogenesis without *in vivo* mesenchymal cells (*14, 17*). Here, we combine 4D multiscale imaging, single-cell RNA sequencing, pharmacological and mechanical perturbations with biophysical modelling, to show that cell fate emergence orchestrates the coordination of mechanical forces via switches in myosin patterns and osmosis. The resulting tissue-scale mechanical responses, which are the differential spontaneous curvature in two regions and the net fluid relocation from lumen to villus cells, determine crypt formation.

**Figure 1:**
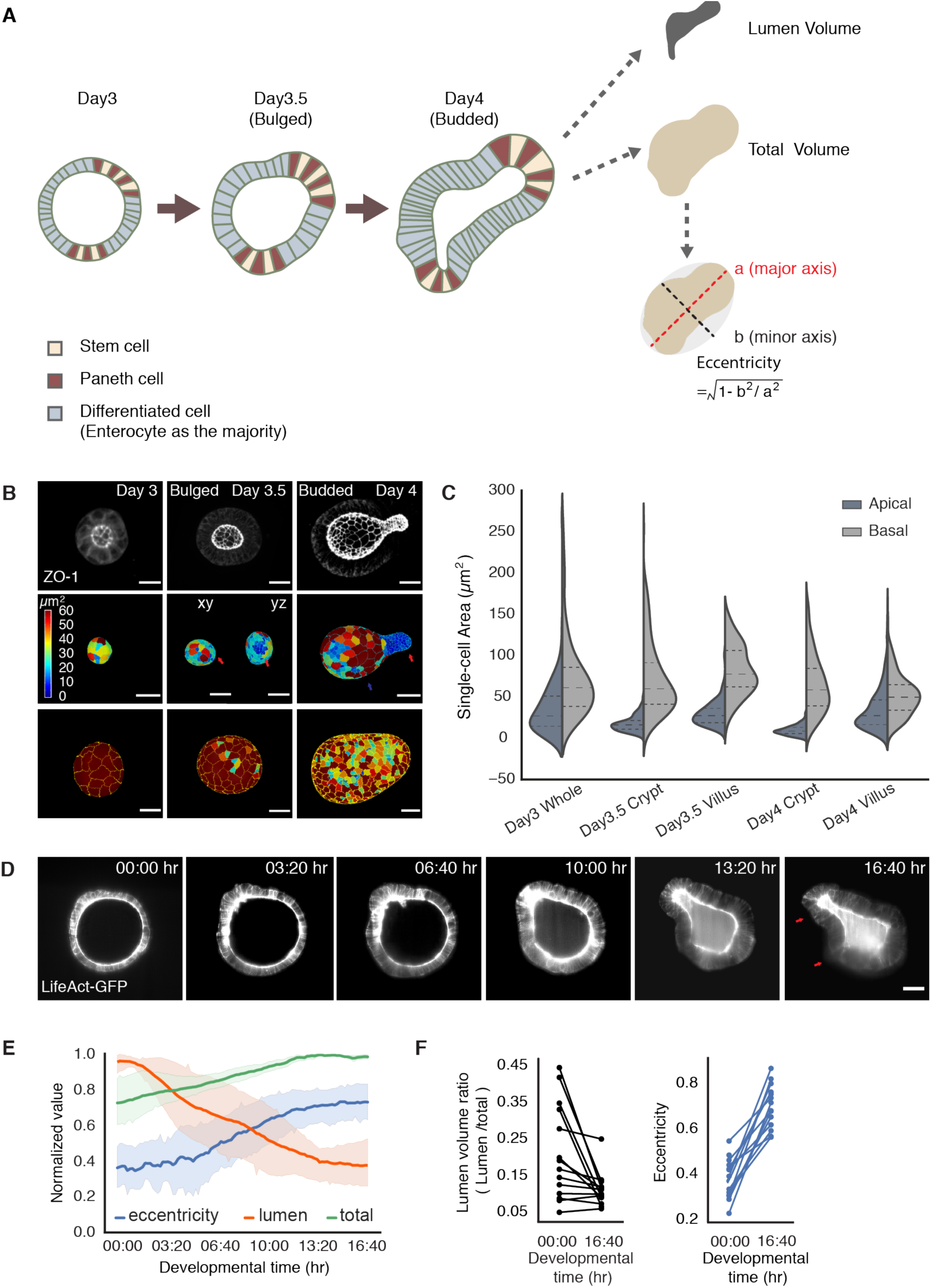
Crypt morphogenesis during intestinal organoids development. **A**. Cartoon representation of crypt morphogenesis and workflow of feature extraction during image analysis. **B**. Segmentation of single-cell apical and basal domains. Top panels, representative time-course imaging of ZO-1 (white) staining in Day3 organoid before bulging (left), Day3.5 bulged organoid (middle) and Day4 budded organoid (right). Middle panels, segmentation of single-cell apical domains corresponding to the top panel, Day3.5 bulged organoid has xy and yz views. Lower panels, segmentation of single-cell basal domains corresponding to the top panels. Red arrows indicate the apical domains of the crypt in Day3.5 bulged and Day4 budded organoids. Colour of heatmap indicates the size of the domains. Scale bars, 20 µm. **C**. Violin plot of the size of single-cell apical and basal domains in B (n = 218 for Day3 whole from 4 organoids, n = 93 for Day3.5 crypt from 3 organoids, n = 187 for Day3.5 villus from 3 organoids, n= 246 for Day4 crypt from 4 organoids, n= 512 for Day4 villus from 4 organoids, two-tailed t-test for apical *vs*. basal domain size in all groups p < 10^−5^, for Day3 whole apical *vs*. Day3.5 crypt apical, p < 10^−11^ ; for Day3 whole apical *vs*. Day4 villus apical, p < 10^−36^ ; for Day3.5 crypt apical *vs*. Day 4 crypt apical, P < 10^−28^ ; for Day3 whole basal *vs*. Day3.5 villus basal, p = 0.14; for Day3 whole basal *vs*. Day4 villus basal, p < 10^−9^; for Day3.5 villus basal *vs*. Day4 villus basal, p < 10^−21^). Violin plot lines denote quartile for each group. **D**. Representative light-sheet time-lapse imaging of crypt bulging and budding in organoid expressing LifeAct-GFP (white), red arrows indicate the crypt regions. Scale bar, 20 µm. **E**. Plot for lumen volume, total volume and eccentricity of organoids in recording in D. Data are collected from 4 individual organoids with 10-mintues time interval, 100 time-points each. Solid lines represent the average values, shadow regions represent the standard deviations. **F**. Quantification of the lumen volume *vs*. total volume ratio (Lumen/total) and eccentricity before bulging at time point 0 (00:00 hr) and after budding at time point 100 (16:40 hr). Paired Student’s *t* test for lumen ratio (n= 15, p < 10^−5^) and eccentricity (n =15, p < 10^−7^).

## Crypt morphogenesis follows crypt apical constriction, villus basal tension and lumen shrinkage

Crypt apical constriction has been reported to initiate crypt morphogenesis *in vivo* (*11*). To validate this and to investigate the global morphological changes occurring during crypt formation in organoids, we initially followed their formation with image based time-course experiments at a high spatial-temporal resolution at Day3, Day3.5 and Day4 (*15*) (Fig. 1A). We quantified the sizes of cell apical and basal membrane domains via segmentation of the staining of the tight junction protein ZO-1 (Fig. 1B). The apical domain of cells in crypts first progressively get smaller (both in absolute terms and compared to villi), showing that organoid crypt cells undergo apical constriction, at a stage we refer to as “bulged” (Fig. 1B and C). In coordination with apical constriction, the crypt monolayer compacts, as shown by a marked reduction of distance between adjacent nuclei (Fig. S1A and B), a feature conserved during *in vivo* crypt morphogenesis (Fig. S1C and D). Interestingly, we noticed that in the villus monolayer, the basal domain of cells reduced its size during development from Day3.5 bulged to Day4 organoid (Fig. 1B and C), indicating a basal constriction in cells from villus, at a later stage of fully-formed crypts which we call “budded”.

To monitor the dynamics of tissue morphological changes during crypt formation, we performed light-sheet time-lapse recording on organoids expressing LifeAct-GFP, the live reporter of actin-filament (F-actin) (*18*). Strikingly, crypt budding was correlated with a dramatic change in lumen volume (Fig. 1D-F and Movie S1). To systematically quantify the phenotypes over 101 time-points in each sample, we applied machine-learning based image segmentation to measure both lumen volume and organoid shape changes (Fig. 1D-F), confirming that a significant lumen shrinkage occurs during crypt budding (higher eccentricity). However, this is compensated by a large volume increase of the epithelium, so that global organoid volume (lumen + cells) increased slightly during morphogenesis (Fig. 1D-F). This suggests that the shrinkage of the lumen volume could have a mechanical contribution to crypt morphogenesis.

## Spontaneous curvature drives crypt morphogenesis

To understand quantitatively the potential contributions of crypt apical contraction, villus basal constriction or lumen shrinkage to organoid crypt morphogenesis, we turned to biophysical modelling and a three-dimensional vertex model description, which is relevant to describe epithelial layers (*19-22*). We modelled organoids as closed monolayers with cells under apical, lateral and basal domain tensions, enclosing a lumen of fixed volume (Fig. S2 and see SI Text for details). Given our results, we allow cell basal, lateral and apical tensions to be different in two spatially distinct regions (crypt and villus, Fig. 2A for a schematic). We then computationally screened parameter-space, to generically understand how each parameter influences the equilibrium shape of an organoid, and derived a simplified model which allowed us to predict simple scaling laws for organoid morphometrics (Fig. 2A). Importantly, we found three classes of mechanisms which could lead to a budded shape (Fig. 2B, see SI Text for details): i) crypt softer (smaller in-plane contraction *α*) than villus, leading to crypt expansion/budding from luminal pressure, ii) differential spontaneous curvature *γ* in crypt *vs*. villus region, leading to crypt bending, iii) contractile boundary of the crypt-villus region, leading to bulging and budding via boundary contraction. However, we found that each class of mechanisms gave rise to qualitatively different predictions on crypt morphometric parameters, *e*.*g*. the ratio of epithelial thickness or of curvature in crypts *vs*. villi (see SI Text for details). For instance, in scenario i), morphogenesis occurs via differential expansion of the budded region, leading to lower epithelial thickness in crypt compared to villus, which is similar to observations of fluid-pressure driven expansion of lung buds/alveoli (*23*), while in scenario ii), bulging and budding occurs via tissue bending, which increases the epithelial thickness in crypts (Fig. 2B).

**Figure 2:**
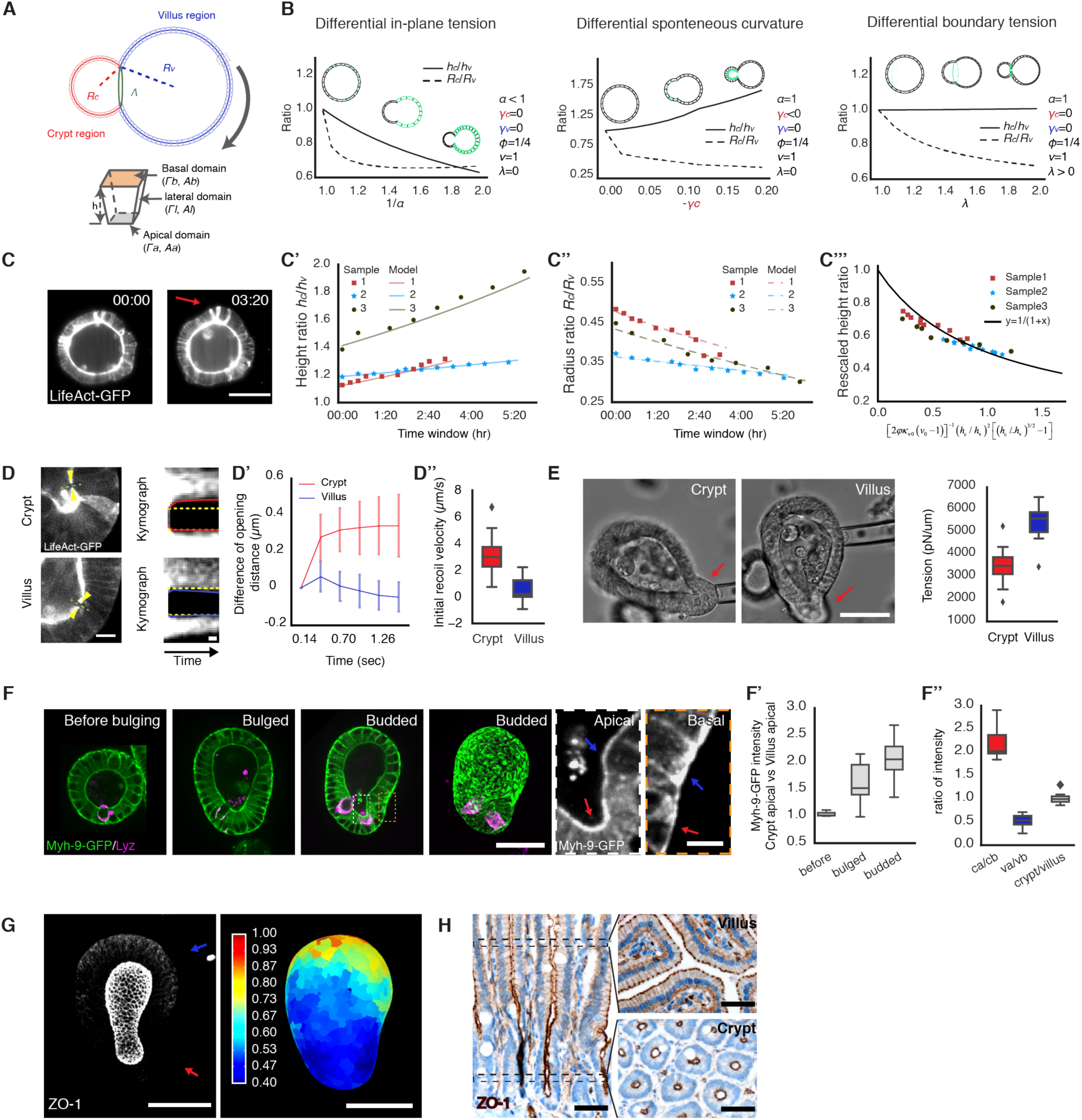
Myosin-patterns determine region-specific spontaneous curvature driving crypt morphogenesis. **A**. Schematic of the theoretical model and morphometric parameters for three-dimensional organoids. Stable organoid configuration is calculated by minimizing the energy *F* (*α, γ_c_, γ_v_, φ, υ, λ*), which depends on a few key parameters (see SI Text for details): α, ratio of in-plane contractions in crypt and villus regions; γ_*c*_, spontaneous curvature crypt region; γ_*v*_, spontaneous curvature of villus region. φ, relative size of the crypt region; υ, normalized lumen volume; λ, potential line tension along the crypt/villus boundary. **B**. Three possible mechanical scenarios that could drive crypt morphogenesis: Left panel, crypt budding driven by smaller crypt in-plane contraction α, leading to thinner crypts (decreased epithelial thickness ratio *h*_*c*_/*h*_*v*_ while radius of curvature *R*_*c*_/*R*_*v*_ decreases). Middle panel, crypt budding driven by spontaneous curvature γ_c_, leading to thicker crypts (increased height ratio *h*_*c*_/*h*_*v*_ while radius of curvature *R*_*c*_/*R*_*v*_ decreases). Right panel, organoid budding driven by line boundary tension, leading to constant thickness (constant height ratio *h*_*c*_/*h*_*v*_ while radius of curvature *R*_*c*_/*R*_*v*_ decreases). The width and location of the green lines indicate the strength and distribution of the driving forces in each model. **C-C’’’**. Experimental data validate the spontaneous curvature model (increased height ratio *h*_*c*_/*h*_*v*_). Images, light-sheet time-lapse recording of representative organoid before and after bulging (C). Plots, thickness ratio (*h*_c_/*h*_*v*_) (C’) and radius ratio (*R*_*c*_/*R*_*v*_) (C’’) from three experimental samples with the corresponding model fit (see SI Text for details), different samples collapse on the same theoretical master curve independent of time (*R*_c_/*R*_*v*_ *vs*. a function of *h*_*c*_/*h*_*v*_) as predicted by the model (black line) (C’’’). **D**. Crypt has higher apical tension than villus. Left panel, imaging of organoid after laser cutting on the apical domain, visualized by LifeAct-GFP (white) in crypt and villus regions. Green dashed lines indicate the position of cutting, yellow arrow heads indicate the end of opening after laser cutting. Middle panel, kymographs across the regions of cutting and opening with surrounded LifeAct-GFP signal. Yellow dashed lines in kymographs outline the size of initial opening, red lines indicate recoil of opening after cutting in crypt and blues lines indicate recoil of opening after cutting in villus regions. D’and D’’, Quantification of the initial recoil velocity in crypt and villus region. Data are collected from time-lapse recording with 0.14-sec time interval, recoil distances from 0.14 to 0.42 sec are used to calculate initial recoil velocities in D’’. Two-tailed t-test for crypt (n = 15) and villus (n = 9), p < 10^−4^. Scale bars, scale bar in organoid image, 10 µm; scale bar in kymograph, 1 µm. Error bars in D’ are SDs. Box plot elements in D’’ show quartiles, and whiskers denote 1.5× the interquartile range. **E**. Villus has higher basal tension than crypt. Left panel, imaging of micropipette aspiration on the basal side of the crypt and villus tissue. Right panel, corresponding quantification of the micropipette aspiration, At steady state, the basal tension γ_basal_ of the organoid is calculated based on Young-Laplace’s law: *γ*_*basal*_ = *P*_*c*_/2(1/*R*_*p*_ – 1/*R*_*o*_), where *R*_*p*_ is the radius of the micropipette (*R*_*p*_ = 15 to 20 µm), *R*_*o*_ is the local radius of curvature of the tissue being aspired, which is measured by experiment. *P*_*c*_ is the pressure used to deform R_o_. Two-tailed t-test for crypt (n = 17) and villus (n = 8), p < 10^−2^. Scale bar, 50 µm. Box plot elements show quartiles, and whiskers denote 1.5× the interquartile range. **F**. Myh-9-GFP is enriched at crypt apical and villus basal domains. Images from left to right, the expression of Myh-9-GFP (green) with Lysozyme (Lyz) staining for Paneth cells (magenta) in organoid before bulging, bulged organoid and budded organoid in the middle z-section, maximum z-projection of Myh-9-GFP in the same budded organoid, zoom-in areas from dashed boxes in the middle section of budded organoid demonstrate the enrichment of Myh-9-GFP (white) at crypt apical domain and villus basal domain. Blue arrows indicate the villus region, red arrows indicate the crypt region. Scale bars, in budded organoid maximum z-projection, 50 µm, in zoom-in regions, 5 µm. **F’**, ratio of Myh-9-GFP intensity in cells at crypt apical domain *vs*. villus apical domain before bulging (n = 30 cells from 3 organoids), bulged (n = 110 cells from 11 organoids) and budded (n = 220 cells from 22 organoids). F’’, ratio of normalized Myh-9-GFP intensity, crypt apical (ca) *vs*. crypt basal (cb), villus apical (va) *vs*. villus basal (vb), crypt (ca + cb) *vs*. villus (va + vb). Numbers, n = 40 cells from 8 organoids for crypt region (ca, cb and crypt), n= 40 cells from 8 organoids for villus regions (va, vb and villus), two-tailed t-test for ca/cb and va/vb (p < 10^−6^), ca/cb and crypt/villus (p < 10^−5^), and va/vb and crypt/villus (p < 10^−4^). Box plot elements show quartiles, and whiskers denote 1.5× the interquartile range. **G**. ZO-1 is enriched in whole apical and villus basolateral domains in budded organoid. Left panel, maximum z-projection of organoids stained with ZO-1 (white). Right panel, heat-map visualization of ZO-1 signal in organoid basal domain. Blue arrow indicates the villus region, red arrow indicates the crypt region. Scale bars, 50 µm. **H**. ZO-1 is localized in epithelial apical domain and villus basolateral domain in adult mouse small intestine. Left image, section along crypt/villus axis. Right images, perpendicular sections of villus and crypt regions along the positions indicated by black dashed rectangles in left image. ZO-1 signal is visualized by oxidized DAB in brown, Cell nuclei are stained with Hematoxylin in blue. Scale bars, 50 µm.

We first modelled the dynamics of bulging, during which we find lumen volume to be almost constant (Fig. S3A and B). Only scenario ii), the differential spontaneous curvature *γ* driven crypt bulging, agreed with the experimental data, as we detect increased thickness ratio (*h*_*c*_/*h*_*v*_) and decreased radius ratio (*R*_*c*_/*R*_*v*_), confirming that crypt apical constriction and spontaneous curvature are a major driving force for crypt morphogenesis (Fig. 2C-C’’, see SI Note for details of the theory and fitting strategy). Importantly, the model predicts that plotting (*R*_*c*_/*R*_*v*_) as a function of (*h*_*c*_/*h*_*v*_) should collapse the data from different organoids on a universal scaling law, independent of the dynamics of tension evolution (Fig. 2C’’’ and Fig. S4, see SI Text for details). Comparing this with the data revealed a good agreement, allowing us to validate the spontaneous curvature mechanism. Furthermore, independently measuring initial organoid and crypt sizes, and assuming a linear increase of crypt apical tension in time allowed for a good fit of the individual temporal evolutions of the ratios (*R*_*c*_/*R*_*v*_) and (*h*_*c*_/*h*_*v*_) (Fig. 2C, Fig. S3C and D and Fig. S4 and SI Text). Next, we sought to further test the model via mechanical measurements, as well as mechanical and pharmacological perturbations.

## Myosin II patterns induce differential spontaneous curvatures

To directly quantify apical tensions in crypt and villus regions, we performed laser nanosurgery on the apical domains of budded organoids (*24*). The analysis showed higher recoil velocities in crypt apical domains after cutting, and thus, apical contraction in crypt is higher compared to villus region (Fig. 2D-D’’). Moreover, through micropipette aspiration experiment (*25*), we found that the basal tension of the villi was higher than that of the crypts (Fig. 2E). The combination of high apical contraction and low basal tension in crypt is thus fully consistent with the prediction of the model in which spontaneous curvature drives crypt morphogenesis (see SI Text for details).

Based on the role of actomyosin in generating active tension (*26*), we decide to investigate if the patterns of F-actin and Myosin correlate with the tensions and spontaneous curvatures that the model predicts. LifeAct-GFP and the phosphorylated Myosin Light chain (pMLC), which binds to Myosin II heavy chain and moves along F-actin to generate contractile force, are enriched in the villus basolateral domain and the apical domain of both crypt and villus region (Fig. S5A and B). However, in the villus apical domain, F-actin and pMLC do not necessarily contribute to apical contraction, but are more likely required for microvilli formation (*27*). To follow the dynamics of Myosin we generated an organoid line from the transgenic mouse of Myh-9-GFP that integrates GFP coding sequence with Myosin IIA encoding gene *myh-9* (*28, 29*). We found that Myh-9-GFP signal gradually increases at the crypt apical side during the developmental process that organoids evolve from near spherical to bulged and budded shapes (Fig. S5C and MovieS2). Moreover, in budded organoids, the expression pattern of Myh-9-GFP is high in crypt apical and villus basolateral domains, low in crypt basolateral and villus apical domains (Fig. 2F, Fig. S5C and Movie S2). These Myh-9-GFP patterns are all in agreement with the region-specific tension changes that are predicted by the spontaneous curvature mechanism (see SI Text).

We next tested whether actomyosin is functionally required for crypt morphogenesis. In mouse intestinal organoids, blocking actomyosin contractility before bulging prevents crypt morphogenesis (Fig. S5B) (10). Moreover, inhibiting myosin activity after budding (via Blebbistatin) disrupted the crypt morphology (Fig. S5D, Movie S3), while blocking cell division (via Aphidicolin) did not (Fig. S5D, Movie S4). Nonetheless, since Myosin IIA exhibits a specific differential pattern in the crypt and villus region, we decided to further validate whether Myosin IIA is necessary for the generation of spontaneous curvature. In order to quantify this, we constructed mosaic organoids composed of Myh-9^+/-^ heterozygous mutant cells and Myh-9-GFP cells (30) (Fig. S5E). The Myh-9^+/-^ cells adjacent to Myh-9-GFP cells have an expanded basal domain in the villus, while the Myh-9-GFP cells contract basally, indicating that Myosin IIA is required for the generation of basal tension which propagates to neighboring cells (Fig. S5E). As a consequence, domain-specific patterns of Myosin IIA modulate cell apical and basal tensions in both crypt and villus, which in turn determine tissue spontaneous curvatures.

To explore the molecular mechanism that contributes to the emergence of the region-specific actomyosin pattern, we analysed the expression patterns of actomyosin regulators, with a recently published single-cell RNA sequencing dataset that distinguishes crypt and enterocyte-rich villus regions (Fig. S6A-B), and by immunostaining in budded organoids (Fig. S6C-E) (*15*). An interesting candidate is the tight junction protein ZO-1 that has recently been reported to orchestrate the apical organization of actomyosin in mouse intestine (*31*). We detected the region-specific expression patterns from both tight junction proteins (ZO-1, Claudin, Occludin) and adherens junction proteins (N-Cadherin) (Fig. 2G, Fig. S6A-E). However, among them, only ZO-1 overlaps with the basal pool of Myh-9-GFP in organoid villus region, and exhibits villus basolateral localization both *in vitro* and *in vivo* during crypt morphogenesis (Fig. 2G-H and Fig. S6F).

## Lumen volume reduction accelerates crypt budding

Next, we explored whether the striking decrease in lumen volume at the onset of crypt budding (Fig. 1D-F) had a functional role for morphogenesis. We hypothesized that, in analogy to bi-layered lipid vesicles with region-specific spontaneous curvature (*32*), lumen volume reduction could contribute to budding by changing the area/volume ratio of the organoid, thus decreasing monolayer tension and promoting out-of-plane deformations. To check this hypothesis, we computed a phase diagram of organoid morphology as a function of normalized lumen volume and relative crypt apical tension (Fig. 3A and Fig.S7, SI Text for details, sensitivity analyses and model extensions). For constant lumen volume, crypts undergo bulging and then budding for increasing values of crypt apical tension (or crypt spontaneous curvature, see Fig. S2D). For low apical tension, budded morphology becomes quasi-spherical again when increasing lumen volume above a critical threshold (Fig. 3A), which ‘unfolds’ the crypt, as expected in lipid vesicles (*32*). Unexpectedly, however, for high values of apical tension, the model predicted that budded shapes cannot be reversed by increasing lumen volume (Fig. 3A). This is because high apical tension increases not only spontaneous curvature in crypts (that leads to budding), but also in-plane rigidity (which resists deformation of crypts and result in preferentially expansion of villi to bear the lumen-induced deformation). This unexpected prediction of a phase transition in the phase diagram (see SI Text for further theoretical exploration) provides a critical test of the model. Moreover, the fits of bulging evolution (Fig. 2C) predict that bulged organoids are well below the transition point (around 40% of the critical apical tension *m*_crit_ at which crypts remain closed upon large inflation, see SI Text for details) in the phase diagram of Fig. 3A, whereas budded crypts should be able to sustain large luminal inflation. We thus proceeded to test this by a series of experiments manipulating lumen volume (Fig. 3A).

**Figure 3:**
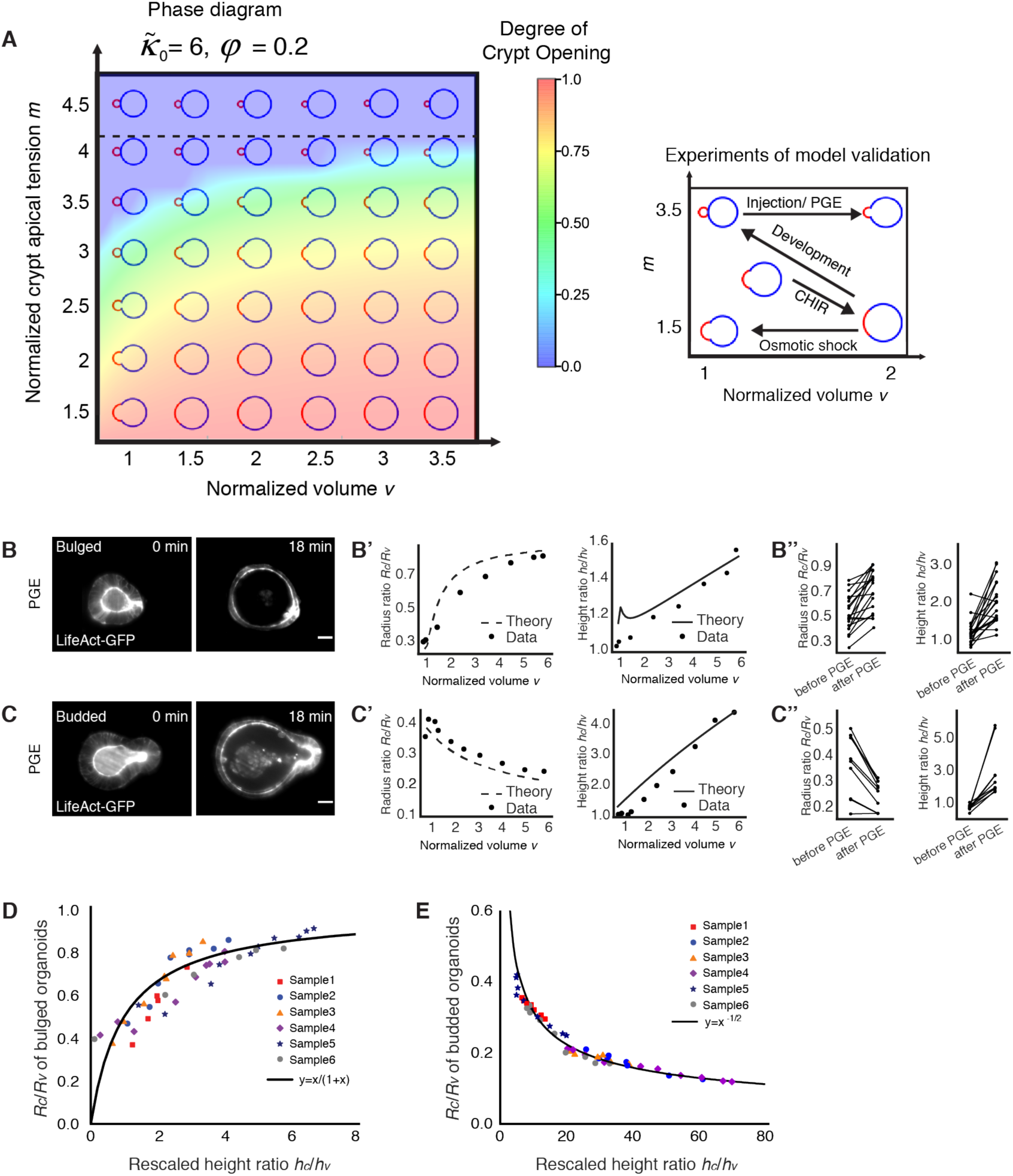
Lumen volume reduction enables crypt budding. **A**. Theoretical prediction for the relative contribution of the crypt apical tension (*m*) versus. lumen volume (*v*) for organoid shape (left panel), and schematic representing the experimental designs for phase diagram validation (right panel). The horizontal dashed line at *m* = 4.2 indicates the critical value, above which the budded shapes cannot be reversed by increasing lumen volume. **B-C’’**. Fitting for the lumen inflation experiments with the theoretical model. Images show organoid morphology before and after lumen inflation by 0.5 µM PGE treatment of bulged (B) and budded (C) organoids with tissue and lumen visualized by LifeAct-GFP (white). Scale bars, 20 µm. Experimental measurements (black dots, data) from the representative samples (cell height/epithelial thickness ratio *h*_*c*_/*h*_*v*_ and radius ratio *R*_*c*_/*R*_*v*,_ B’ for B, bulged organoid, C’ for C, budded organoid) as a function of lumen volume (*v*) change were fitted from the model (black lines, see SI Text for details and best-fit values). Budded crypts systematically remained near-unchanged by inflation (decreased radius ratio *R*_*c*_/*R*_*v*_) (B’), while bulged crypts opened up (increased radius ratio *R*_*c*_/*R*_*v*_) (C’). Plots, measurements of epithelial thickness ratio and radius ratio in bulged (B’’, n= 19) and budded (C’’, n = 10) organoids. Paired Student’s *t* test for bulged *R*_*c*_/*R*_*v*_ (p < 10^−3^), bulged *h*_*c*_/*h*_*v*_ (p < 10^−5^), budded *R*_*c*_/*R*_*v*_ (p < 10^−2^) and budded *h*_*c*_/*h*_*v*_ (p < 10^−3^). **D-E**. All samples from PGE and pipette inflation (bulged: D, budded: E) can be collapsed via fitting of two parameters (see SI Text for details) on the same theoretical master curve (*R*_*c*_/*R*_*v*_ *vs*. a function of *h*_*c*_/*h*_*v*_) independent of volume as predicted by the model (black line), showing opposite trends in bulged (D) *vs*. budded (E).

We first inflated the lumen in a time frame of less than 30 minutes by treating organoids with Prostaglandin E2 (PGE), the lipid autacoid that induces lumen swelling but does not alter cell fate in the first 6 hours of the treatment (*33, 34*), and measured several morphometric parameters as a function of organoid lumen volume (Fig. 3B-C’’ and Movie S5). To explore the phase diagram of Fig. 3A, we performed the experiments both in early (bulged) and late (budded) organoids, which have intermediate and high crypt apical tensions based on Myh-9-GFP intensity (Fig. 2F). Crucially, we found that budded crypts could not be opened by PGE treatment (even increasing volume up to 498%, which stretches nearly exclusively the villus region, leading to decreasing radius ratio *R*_*c*_/*R*_*v*_, Fig. 3C). In contrast, volume expansion in bulged crypts lead to a near-spherical shape (increasing radius ratio *R*_*c*_/*R*_*v*_, Fig. 3B), in excellent agreement with the model. To further confirm this, we performed fluid microinjection via pipette to rapidly and controllably increase lumen volume, and also found the opposite behaviour in bulged *vs*. budded organoids (Fig. S8A-B’’ and Movie S6).

To test this further quantitatively, we analytically derived scaling laws to predict how thickness and radius ratios (crypts/villus, resp. *h*_*c*_/*h*_*v*_ and *R*_*c*_/*R*_*v*_) should vary as a function of lumen volume, for bulged and budded organoids (see SI Text for details). Strikingly, upon fitting of two parameters (crypt apical tension *m* and initial lumen volume *v*_0,_ which gave values consistent with the ones fitted from Fig. 2C, see SI Text for details), all inflation data (PGE and pipette) could be collapsed onto these predicted scaling laws (Fig. S8C and C’), providing excellent agreement between theory and experiments, including opposite trends of the radius ratio *R*_*c*_/*R*_*v*_ as a function of volume in bulged *vs*. budded crypts (Fig. S8C and C’). With the same parameter set, the model also predicted simple scaling relationships between *R*_*c*_/*R*_*v*_ and *h*_*c*_/*h*_*v*,_ independent of lumen volume (see SI Text for details), which again were in excellent agreement with the data (Fig. 3D and E).

Finally, we performed lumen volume reduction, in order to achieve an opposite effect than the lumen inflation experiments (Fig. 3A and Fig. S8D). Three types of organoids were treated with the 250 mM NaCl osmotic shock: i) Day2.5 organoids without crypt/villus regional difference, ii) Day3.5 organoids with established regional difference in crypt and villus, and iii) Day3.5 organoids treated with CHIR (CHIR99021, a highly selective GSK3 inhibitor) that only consist of crypt region (Fig. S8D and Movie S7) (*15*). Seconds after the osmotic deflation, the Day3.5 organoids formed bulges in the crypt regions enriched in Lgr-5^+^ stem cells, while the Day2.5 and Day3.5 CHIR-treated organoids remained spherical and did not display any significant bulging (Fig. S8D and Movie S7). This provides direct evidence that the reduction of the lumen volume can efficiently accelerate the spontaneous-curvature dependent crypt morphogenesis in intestinal organoids.

## Osmotic changes via membrane transporters in enterocytes drives lumen shrinkage

Next, we addressed the cellular and molecular mechanisms of volume reduction. Given the fact that overall organoid volume varies little (Fig. 1D-E), we hypothesized that lumen volume could be redistributed to epithelial cells during morphogenesis. To distinguish whether a specific cell type was responsible for the lumen volume reduction, we analysed organoids that are enriched with different cell types (Fig. 4A). We found that enterocysts, organoids containing only enterocytes (*15*), reduce lumen volume (Fig. 4B and D, Fig. S9A and Movie S8), whereas CHIR-treated organoids, composed mainly of stem cells and Paneth cells (*15*), increase it (Fig. 4C and E, Fig. S9A and Movie S9), and neither of these two kinds of organoids can bud (Fig. 4 A-E, Fig. S9A and Movie S8 and S9) (*15*). These results suggest that the reduction of organoid lumen volume is dependent of enterocytes, which are the major component of the villus region. In agreement with these data, our measurements show that single-cell volume of villus region increases during crypt budding, while single-cell volume of crypt region does not (Fig. S9B). We then refined our biophysical model, to include that lumen volume reduction is compensated by either i) volume increase of all cells, ii) volume increase of crypt cells only, iii) volume increase of villus cells only. Strikingly, we found that hypothesis iii) is the most efficient to produce budded crypts, as it specifically increases the compressive stresses exerted by villus on the crypt region, while increasing the volume of crypt cells would be detrimental for budding as it would increase the bending rigidity of crypts (Fig. 4F and see SI text for details). This strongly suggest that luminal fluid uptake specifically by villus cells is an efficient and robust mechanism to promote crypt budding.

**Figure 4:**
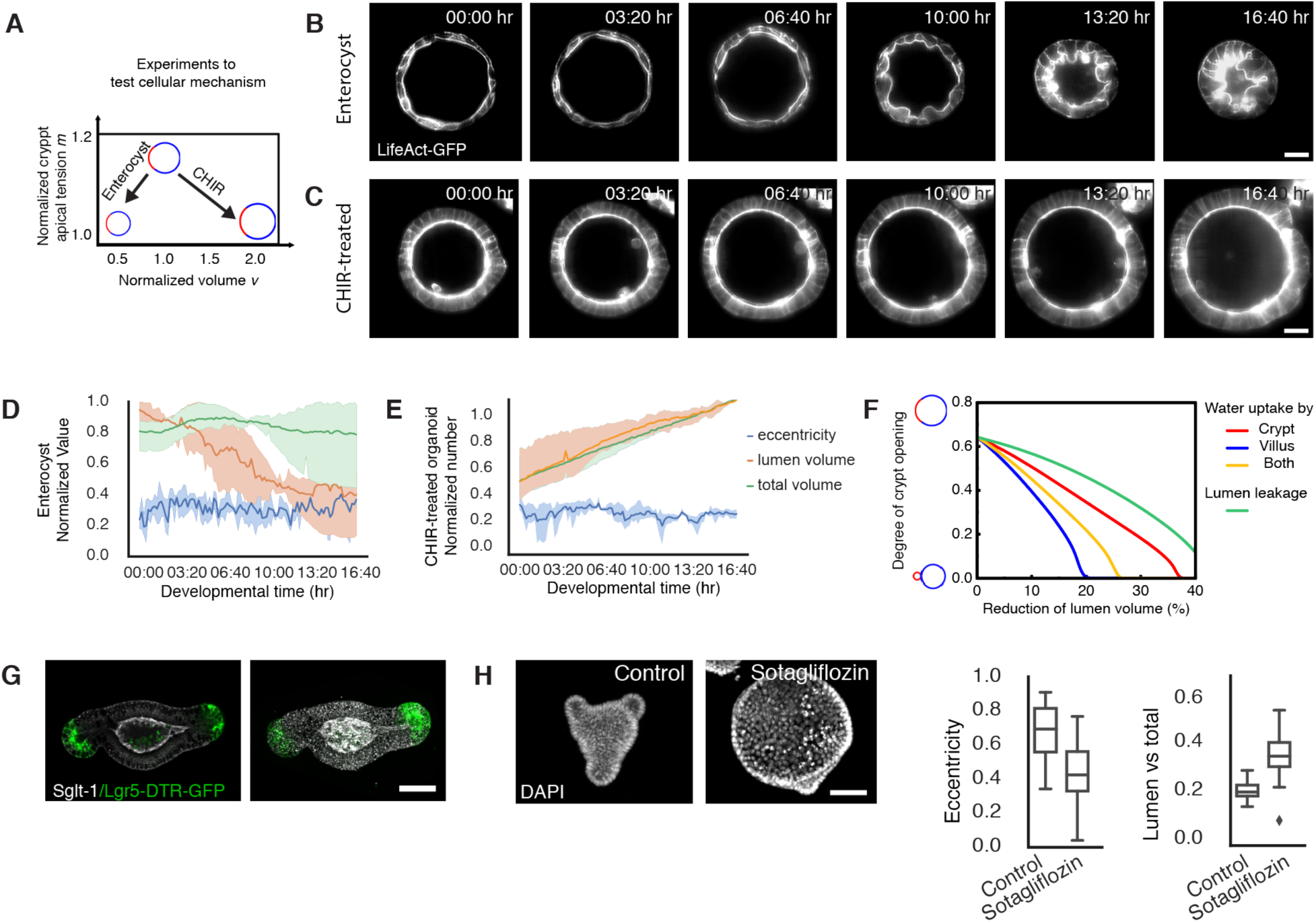
Enterocytes control lumen volume reduction through SGLT-1. **A**. Schematic representation of the experimental design applied. **B**. Light-sheet time-lapse recording of enterocyst, the organoid that is composed of only enterocyte, from Day3 with tissue and lumen visualized by LifeAct-GFP (white). **C**. Light-sheet time-lapse recording of organoid treated with 3 µM CHIR, the organoid that is highly enriched with stem cells and Paneth cells, from Day3 with tissue and lumen visualized by LifeAct-GFP (white). Scale bars, 20 µm. **D**. Corresponding plots of B on eccentricity, lumen volume and total volume (lumen + tissue) (n=4). **E**. Corresponding plots of C on eccentricity, lumen volume and total volume (n =3). **F**. Model prediction of how increased volume of different regions promotes budding, showing that fluid uptake specifically by villus cells is best for budding. **G**. Immunostaining of SGLT-1 (white) in budded organoid with crypt regions visualized by Lgr-5-DTR-GFP (green) indicates the enrichment of SGLT-1 in the villus apical domain, Scale bar: 50 µm. **H**. SGLT-1 is required for lumen volume reduction. Images, Day4 organoid of 5 µM DMSO-treated control and 5 µM Sotagliflozin-treated sample for SGLT-1 inhibition. Scale bar, 20 µm. Plots, quantification of the eccentricity (left plot) and lumen ratio (lumen volume *vs*. total volume, right plot) of control (n = 80 from three replication) and Sotagliflozin treated sample (n= 46 from three replication). Two-tailed t-test for eccentricity (p < 10^−5^), and lumen ratio (p < 10^−8^). Box plot elements show quartiles, and whiskers denote 1.5× the interquartile range.

Finally, to test mechanistically how lumen volume reduction is regulated in enterocytes, we analysed membrane transporters that regulates osmotic gradient and water transfer and selected three groups of candidates based on their expression in scRNAseq data. We found that the family members of the Sodium/glucose cotransporter (SGLT-1), the water channel Aquaporins (AQPs) and the Na^+^/K^+^ ATPases (Atp1a1 and Atp1b1) have enterocyte-specific mRNA expression (Fig. S6A and Fig. S9C). Interestingly, at the protein level, only the SGLT-1 is enriched at villus apical domain (Fig. 4G), while AQPs and Na^+^/K^+^ ATPase are expressed homogenously throughout the epithelium (*35*). Inhibition of each of the membrane transporters lead to the failure of lumen volume reduction and organoid budding (Fig. 4H and Fig. S9D). Remarkably, the inhibition of the SGLT-1 using sotagliflozin leads to the strongest defects (Fig. 4H and Fig. S9D), supporting the role of enterocytes in lumen volume reduction. The SGLT-1 transporters in enterocytes can actively change the osmotic concentrations in the luminal fluid, driving the passive export of fluid from the lumen and supporting the role of enterocytes in lumen volume reduction.

## Discussion

Here we demonstrated that apical contraction in crypts and basal tension in the villus generate the region-specific spontaneous curvatures that lead to crypt bulging and budding, and that enterocytes further promote this process by uptaking fluid and reducing lumen volume. Apical contraction drives tissue folding in morphogenesis of many systems, such as the *Drosophila* mesoderm invagination, the vertebrate neural tube closure, the morphogenesis of mouse lens placode and the gastrulation in various organisms (*36*-*40*). Basal tension has more recently been revealed to be a driving force for tissue morphogenesis (*20, 41*). In our model, two key requirements to reproduce the experimental data are the presence of differential cellular spontaneous curvatures in crypt *vs*. villus during bulging and budding, while exclusively at budding stage a larger in-plane contraction of crypts compared to villus (see SI Text for further discussion on the roles of basal *vs*. apical tension). Together, these two features result in a critical value of crypt apical tension, above which crypt shape becomes fully robust to any value of lumen inflation, which could be relevant to *in vivo* pathological situations such as diarrhea caused by bacteria or virus infection (*42, 43*).

Interestingly, it has been reported recently that basal tension in MDCK cells leads to the increased spontaneous curvature, which is able to bend the 2D-cultured cell layer (*44*). The basal tension in the villus region emerges during enterocytes differentiation and is likely regulated by specific re-localization of ZO-1 to the basolateral domain of the villus. The actomyosin regulator, Rac-1, is required for the basal constriction of hinge cell that guides crypt spacing *in vivo* (*11*), and could play a role in longer-term maintenance of organoid crypt shape. Future study on the region-specific regulation of actomyosin, including the causality between the actin-binding junction protein at the villus basal side, will help to uncover the molecular regulation of tissue mechanical property during crypt morphogenesis. Because differential patterns of actomyosin and ZO-1 are also observed in crypts *in vivo* (Fig. 2H) (*45*), our findings can be extended to *in vivo* gut development and crypt morphogenesis. In mouse, the space between villi is strongly reduced during crypt morphogenesis (Fig. S10), suggesting a possible role of lumen volume reduction and/or villus cell compaction also *in vivo*. Indeed, from the perspective of our model, lumen reduction in organoids and villus cell compaction helps curvature-driven bending by increasing compressive stresses within the epithelium. Thus, *in vivo* changes in villus cell density may play a similar role to the one we decipher for lumen volume changes in organoids.

To conclude, we find that cell differentiation into crypt and villus regions create differential mechanical and epithelial osmotic properties. Osmotic changes in enterocytes drive lumen volume reduction, which can coordinate the forces imposed on different crypts in a given organoid. Such coupling between cell fate, mechanics and osmotic changes thus allows for stable and robust shape transformation. Interestingly, although fully formed budded crypts are able to sustain extensive fluid inflation, preventing lumen volume reduction prior to morphogenesis results in partially bulged, immature crypts, pointing to additional potential feedbacks between luminal volume and crypt apical tension. Multiple cellular processes have been proposed to be curvature-dependent, which could provide such feedback (*46*), as would mechano-sensitive processes such as force-dependent adherens and tight junction maturation (*47, 48*). Fluid can exert significant morphogenetic forces, as shown in early mouse embryo or lung morphogenesis (*23, 49, 50, 51*), which are rapidly transmitted spatio-temporally. The manipulation of lumen volume in combination with biophysical modelling, shown here, is thus a powerful tool and could be broadly applicable to other model systems. Overall, comprehensive understanding of the spatial-temporal coordinated behaviours during development and regeneration requires a link between large-scale tissue mechanics and molecular and cellular functions.

## Acknowledgements

We acknowledge the Lennon-Duménil laboratory for sharing the mouse line of Myh-9-GFP with the Hiiragi laboratory. We are grateful to the members of the Liberali laboratory and the FMI facilities for their support. We thank E. Tagliavini for IT support, L. Gelman for assistance and training, S. Bichet and A. Bogucki for helping with histology of mouse tissues, H. Kohler for FAC sorting, G. Q. G. de Medeiros for maintenance of light-sheet microscopy, M. G. Stadler for scRNA-seq analysis, G. Gay for discussions on 3D vertex model, the members of the Liberali laboratory, C. P. Heisenberg and C. Tsiairis for reading and feedback on the manuscript. Funding: Q.Y. is supported by the Postdoc fellowship from Peter und Taul Engelhorn Stiftung (PTES). This work received funding from the ERC under the European Union’s Horizon 2020 research and innovation programme (grant agreement no. 758617 to P.L.) and Swiss National Foundation (POOP3_157531 to P.L.). The Hannezo laboratory acknowledges funding from the Austrian Science Fund (FWF) (P 31639 to E.H.).

## Authors contributions

P.L. and Q.Y. conceived the project and designed the experiments. Q.Y. performed and analysed the experiments. E.H. proposed the physical theory. S.L.X. designed the physical model, performed the simulations. C.J.C. supported the micropipette aspiration. M.R. and D.V. assisted with the image processing and quantifications. F.M.G. generated organoid lines. T.H. helped design experiments. Q.Y. wrote the first version of the manuscript, S.L.X. wrote the first version of supplementary text, P.L., E.H. and Q.Y. supervised the study. P.L. and E.H. helped design the project and finalize the manuscript.

## Notes

### Competing Interest Statement

The authors have declared no competing interest.

