## Supplementary text and figures for "Cell fate coordinates mechano-osmotic forces in intestinal crypt morphogenesis"

**This PDF file includes:**

Materials and Methods
Supplementary Text
Figs. S1 to S10
Tables S1
Captions for Movies S1 to S9

**Other Supplementary Materials for this manuscript include the following:**

Movies S1 to S9

#### Materials and Methods

##### Animal work

All animal experiments were approved by the Basel Cantonal Veterinary Authorities and conducted in accordance with the Guide for Care and Use of Laboratory Animals. Seven C57BL/6 outbred mice from the same litter and the mother were used to obtain small intestines at the age of P1 (two mice), P2 (two mice), P5, P7, P11 and 6-month for time-course immunohistochemistry. Male and female outbred mice from 7 weeks old onwards were used for all experiments generating organoid lines.

Mouse lines used: C57BL/6 wild type (Charles River Laboratories), Lgr5-DTR-EGFP (Genentech, de Sauvage laboratory), LifeAct-GFP, Myh-9-GFP and Myh-$9^{+/-}$ (T. Hiiragi laboratory, EMBL).

##### Organoid culture

Organoids were generated from isolated crypts of the murine small intestine as previously described (*15*). In brief, the section of jejunum was opened lengthwise, cleaned with cold PBS and, after removal of villi by scraping with a cold glass slide, sliced into small fragments roughly 2 mm in length. The tissue was then incubated in 2.5 mM EDTA/PBS at 4 °C for 30 min with shaking. Supernatant was removed and pieces of intestine were re-suspended in DMEM/F12 with 0.1% BSA. The tissue was then shaken vigorously. To collect the first fraction, the suspension was passed through a 70 µm strainer.

The remaining tissue pieces were collected from the strainer and fresh DMEM/F12 with 0.1% BSA was added, followed by vigorous shaking. The crypt fraction was again collected by passing through a 70 µm strainer. In total, 4 fractions were collected. Each fraction was centrifuged at 300 g for 5 min at 4 °C. Supernatant was removed and the pellet was re-suspended into Matrigel with medium (1:1 ratio) and plated into 24 well plates. Organoids were kept in IntestiCult Organoid Growth Medium (STEMCELL Technologies) with 100 µg/ml Penicillin-Streptomycin for amplification and maintenance.

##### Time course experiments of fixed organoid samples

The method was adapted from descripted before (*15*). Organoids were collected 5-7 days after passaging and digested with Tryple (Thermo Fisher Scientific) for 20 min at 37 °C. Dissociated cells were passed through a cell strainer with a pore size of 30 µm (Sysmex). For all experiments, single alive cells were sorted by FACS (Becton Dickinson FACSAria cell sort or Becton Dickinson Influx cell sorter). Forward scatter and side scatter properties were used to remove cell doublets and dead cells. Sorted cells were collected in ENR medium composed of advanced DMEM/F-12 with 15 mM HEPES (STEM CELL Technologies) supplemented with 100 µg/ml Penicillin-Streptomycin, 1×Glutamax (Thermo Fisher Scientific), 1×B27 (Thermo Fisher Scientific), 1xN2 (Thermo Fisher Scientific), 1mM N-acetylcysteine (Sigma), 500ng/ml R-Spondin (kind gift from Novartis), 100 ng/ml Noggin (PeproTech) and 100 ng/ml murine EGF (R&D Systems). Collected cells were mixed with Matrigel (Corning) in a medium to Matrigel ratio of 1:1. In each well of a 96 well plate, 5µl droplets with 2500 cells were seeded. After 15 min of solidification at 37 °C, 100 µl of medium was overlaid. From day 0 to day 1, ENR was supplemented with 25% Wnt3a-conditioned medium (Wnt3a-CM), 10 µM Y-27632 (ROCK inhibitor, STEMCELL Technologies) and 3 µM of CHIR99021 (GSK3B inhibitor, STEMCELL Technologies, cat # 72054). From day 1 to 3 ENR was supplemented with 25% Wnt3a-CM and 10 µM Y-27632. From day 3 to 5, only

ENR was added to the cells. Wnt3a-CM was produced in-house by Wnt3a L-cells (kind gift from Novartis).

**Compound treatments**

Single cells derived from LifeAct-GFP organoids were plated in a 96-well plate chamber and exposed to 7.5 µM (±)-Blebbistatin (Myosin II inhibitor, Abcam cat # AB120425) or 1 µM Cytochalasin D (inhibitor of actin polymerization, Abcam cat # AB143484) or 0.6 µM Aphidicolin (DNA polymerase inhibitor, SIGMA-ALDRICH, cat # A4487) or 0.5 µM Prostaglandin E2 (PGE, kind gift from Novartis) or 5 µM or 10µM DMSO (SIGMA-ALDRICH, cat # D8418) diluted in ENR medium, from 72 hours until fixation at 96 hours (96-well plate) or from 96 hours until changing medium at 114 hours (Light-sheet chamber) or from 84 or 96 hours for 30 minutes (Light-sheet chamber, PGE inflation).

Single cells derived from organoids C57BL/6 wild type were plated in a 96-well plate and treated with 3 µM CHIR99021 (GSK3B inhibitor, STEMCELL Technologies cat # 72054) or 2 µM IWP-2 (Porcupine Inhibitor, STEMCELL Technologies cat # 72124) or 500 µM Ouabain octahydrate ($Na^+/K^+$ ATPase inhibitor, SIGMA-ALDRICH cat # O3125) or 5 µM Sotagliflozin (SGLT1/2 inhibitor, MedChemExpress cat # HZ-15516) or 12.5 µM CuSO4 (Aquaporin inhibitor, SIGMA-ALDRICH cat # C2284) or 5 µM DMSO in ENR medium, from 72 hours until fixation at 96 hours.

**Organoid immunostaining and imaging**

The method was adapted from descripted before (*15*). Organoids embedded in a Matrigel droplet were fixed in 4% PFA (Electron Microscopy Sciences) in PBS for 45 min at room temperature or 4 degree overnight. Fixed organoids were permeabilized with 0.5% Triton X-100 (Sigma-Aldrich) for 1 h and blocked with 3% Donkey Serum (Sigma-Aldrich) in PBS with 0.1% Triton X-100 for 1 h. Primary and secondary antibodies were diluted in blocking buffer and applied as indicated in **the TableS1**. Cell nuclei were stained with 20 µg/ml DAPI (4',6-Diamidino-2-Phenylindole, Invitrogen) in PBS for 5 min at room temperature. Cells were stained with 1 µg/ml of Alexa Fluor® 647 carboxylic acid succinimidyl ester (CellTrace, Invitrogen) in carbonate buffer (1.95 ml of 0.5 M $NaHCO_3$, 50 µl of 0.5 M $Na_2CO_3$, both from Sigma-Aldrich, in 8 ml of water for 10 ml of buffer).

High-throughput imaging was done with an automated spinning disk microscope from Yokogawa (CellVoyager 7000S), with an enhanced CSU-W1 spinning disk (Microlens-enhanced dual Nipkow disk confocal scanner), a 40x (NA = 0.95) Olympus objective, and a Neo sCMOS camera (Andor, 2,560 × 2,160 pixels). For imaging, an intelligent imaging approach was used in the Yokogawa CV7000 (Search First module of Wako software). For each well, one field was acquired with 2x resolution in order to cover the complete well. This overview fields were then used to segment individual organoids on the fly with a custom written ImageJ macro which outputs coordinates of individual organoid positions. These coordinated were then subsequently imaged with high resolution (40x, NA = 0.95). For each site, z-planes spanning a range up to 90 µm and 2 µm z-steps were acquired.

Confocal imaging of fixed samples was performed using Nikon Ti2-E Eclipse Inverted motorized stand with Yokogawa CSU W1 Dual camera (CAM1 SN: X-11424; CAM2 SN:11736) T2 spinning disk confocal scanning unit, CFI P-Fluor 40x oil/1.4 objective and Visview 4.4.0.9

software. Laser lines used are Toptica iBeam Smart 405/488/639 nm and Cobolt Jive 561 nm. Laser power and digital gain settings were unchanged within a given session to permit direct comparison of expression levels among organoids stained in the same experiment. Image stacks were acquired with slice thickness of 2 μm or less.

**Time-course image analysis**

Organoid segmentation in MIPs was adapted from before (*15*). For each acquired confocal z-stack field, maximum intensity projections (MIP) were generated. All MIP fields of a well were stitched together to obtain MIP well overviews for each channel. The high resolution well overviews were used for organoid segmentation and feature extraction. To this end, we apply the segmentation method from Serra, Mayr et al. (2019), which separates organoid instances observed until 72h post-fixation with a watershed algorithm and with an FCN-augmented watershed for organoids observed at or after 72h post-fixation. For the organoid segmentation based on DAPI, we substitute the segmentation of Serra, Mayr et al. (2019) by a U-Net-like FCN trained to generate instance embeddings using fixed bandwidth and soft jaccard loss (*51*). Artifacts of segmentations were semi-automatically annotated in Fiji and removed by a customized python script. From the segmented MIPs, we calculate the 2D eccentricity of each individual organoid using scikit-image's regionprops method. Using the individual organoid masks from the MIP, we crop Z-stacks from the full 3D images (Yokogawa CV7000) and segment them individually with the same FCN used before (*15*). The 3D volume is approximated by the sum of voxel volume segmented as foreground.

Single-cell apical and basal domains and single-cell volume was quantified in 3D using the MorphoGraphX software. Images were segmented computationally and then manually validated (*52*, *53*).

Distance between cell nuclei were measured by Fiji and a customized python script. The outline of each nuclei was selected one after another along crypt to villus axis in Fiji to obtain the coordinate of its centroid and the size of the nucleus. The distance between neighbor cell nuclei is then calculated based on the location of the centroids and normalized by dividing the sum of the radii of the two nuclei that are approximated from their sizes.

Myh-9-GFP intensity ratios were measured using FIJI. A square of 1x1μm was drawn on the middle of cell apical or lateral or basal membrane in the image of the middle section of a z-stack. After measuring the mean gray value for the intensity of the signal on the membranes, the square was shifted to another cell. In each crypt or villus region, more than five cells that are away from crypt/villus border were randomly selected for measurements. The mean gray value for membrane signal was then normalized by comparing the ratio of signal within same cell or same organoid (see term of ratios in Fig. 2F).

**Light-sheet microscopy.**

Light-sheet microscopy was conducted by using LS1 Live light sheet microscope system (Viventis) or a similar customized microscope system as described before (*15*). Sample mounting was performed as described previously (*15*). For organoids imaging, LifeAct-GFP and Myh-9-GFP organoids were collected and digested with TrypLE (Thermo Fisher Scientific) for 20 min at 37 °C. GFP positive cells were sorted by FACS and collected in medium containing advanced DMEM/F-12 with 15 mM HEPES (STEM CELL Technologies) supplemented with 100 μg/ml

Penicillin-Streptomycin, 1×Glutamax (Thermo Fisher Scientific), 1×B27 (Thermo Fisher Scientific), 1xN2 (Thermo Fisher Scientific), 1mM N-acetylcysteine (Sigma), 500 ng/ml R-Spondin (kind gift from Novartis), 100 ng/ml Noggin (PeproTech) and 100 ng/ml murine EGF (R&D Systems). 2000 cells were then embedded in 5 ul drop of Matrigel/medium in 50/50 ratio. Drops were placed in the imaging chamber and incubated for 20 min before being covered with 1ml of medium. For the first three days, medium was supplemented with 20% Wnt3a-CM and 10 µM Y-27632 (ROCK inhibitor, STEMCELL Technologies). For the first day, in addition, 3 µM of CHIR99021 (STEMCELL Technologies) were supplemented. After more than 2 days culture in a cell culture incubator the imaging chamber was transferred to the microscope kept at 37 °C and 5% $CO_2$. Different organoids were selected as starting positions and imaged every 10 min for up to 4 days, or in PGE treatment, every 3 min for up to 30 min. A volume of 150-200µm was acquired with a Z spacing of 2µm between slices. Medium was exchanged manually under the microscopy every half day.

**Light sheet data analysis.**

For area and volume calculation of light sheet data we proceeded as follow. Initially raw data were cropped around the minimum organoids bounding box in order to reduce storage space. To quantify morphological futures as well as lumen and tissue volume during organoid development, a segmentation of lumen and tissue is generated. To this end, a U-Net-like FCN that was trained to predict probability maps for lumen and tissue based on stacks of three XY-planes is applied to the volume in a sliding window fashion along Z. To improve segmentation quality, we specialize one version of this FCN on enterocysts and one on budding organoids by refining the model on the respective part of the dataset. Segmentations are manually inspected to ensure appropriate quality. Organoid and tissue volume is approximated by the sum of voxel volume segmented as foreground. From the segmented z-projection of each time frame, the 2D eccentricity is extracted from the segmented regions using scikit-image's regionprops method.

Using Fiji, we extract the values of morphometric parameters (epithelial thickness and region radius) from movies of intestinal organoids. The epithelial thickness of each region (crypt/villus) is obtained by averaging the data measured in at least two random positions. The radius $Rc$ (or $Rv$) is defined as the average value of the apical and basal radii. For each region, we first try to find the center of an apical/basal circle (basically by the lateral texture of epithelium), then draw a circle that shares the same center and matches the apical/basal edge. The corresponding radii are extracted from these circles in Fiji measurement function. The length of organoid major axis and neck after budding are measured manually in Fiji and converted to aspect ratio = neck / major axis.

**Immunohistochemistry and data analysis**

Jejunums were dissected for fixation in 10% PFA (Electron Microscopy Sciences) in PBS for 24 hours at 4 degree, washed with 70% EtOH, embedded in paraffin, and 3-µm sections prepared and processed for haematoxylin staining and immunohistochemistry. Immunohistochemical staining was performed on formalin-fixed, paraffin-embedded tissue sections using a Ventana BenchMark fully automated stains system with anti-ZO-1 (Thermo Fisher Scientific #33-9100, 1:50) or anti-CD44v6 (BioRad #MCA1967, 1:100). Distance between cell nuclei were measured in the same method as for organoid (see Time-course imaging analysis). Distance between villi were measured in method as follow. The curves $c_i$ and $c_o$ of two closed edges of neighbour villi were annotated using Fiji's ROI Manager and exported as ROI files. In a second step, the files were imported in

Python and a bivariate polynomial $p$ was fitted to each imported curve. $p$ internally stores two univariate polynomials $p_x$ and $p_y$ of degree 3 which approximate the x and y coordinates for each point on the curve, respectively. We then divided $p$ in 29 partitions, each holding a start point and an end point, which results in a total of 30 points along the polynomial. Given $p_i$ and $p_o$, two bivariate polynomials fitting $c_i$ and $c_o$, we compute the shortest distance between any point on $p_i$ and any point on $p_o$ using the Nelder-Mead algorithm. We define the obtained global shortest distance as an approximation to represent the space between villi. Regions for all measurements were randomly selected for all group.

**Tension Measurements by Laser Nanosurgery**

The method was adapted from previous studies (*24*, *54*). In brief, images are captured by a LSM710 scanning confocal microscopy using ZEN Black software. The microscope is equipped with an incubation chamber to keep the sample at 37°C and to provide 5% of $CO_2$. Organoids were embedded in Matrigel and cultured in ibidi 8-well plates. Random samples were selected for cutting. Cutting of the apical membranes was performed using a wavelength of 850nm with a Chameleon Ultrall Laser. The input power was set at 2.17± 0.05W, the length of the cutting region (green dashed lines in Fig. 2D) was set with 2 um length, and the activation time was calculated by the scan speed of 1.25 ms/pixel. Organoids were imaged at 0.13 sec intervals with a Plan-Neofluar 40x/0.9 Imm Korr Ph3 Objective lens before and after cutting. For each cut, we measured the opening distance in every second time frame after cutting as indicated in red and blue lines in Kymograph images in Fig. 2D and compare it to the initial opening distance (yellow dashed lines in Kymograph), the difference of the first two frames (0.14 to 0.42 sec) is applied to calculate initial average recoil velocity as $v_{initial}$ = difference of opening distances /duration of recoil (μm/sec).

**Organoid basal tension measurement.**

The method was adapted from previous studies (*25*, *49*). In brief, a microforged micropipette coupled to a microfluidic pump (Fluigent, MFCS) was used. To measure the basal tissue tension of organoid, micropipettes with radii 15-20 um (covering 3 to 5 cells) were used to apply step-wise increasing pressures on these basal side until a deformation with the same radius of the micropipette ($R_p$) is reached. At steady state, the basal tension $\gamma$ of the organoid is calculated based on Young–Laplace's law: $\gamma = P_c/2(1/R_p - 1/R_c)$, where $P_c$ is the aspiration pressure used to deform the tissue with a local radius of curvature $R_c$. The measurements were performed on an inverted Zeiss Axio Observer microscope equipped with a dry x20/0.8 PL Apo DICII objective. The inner walls of the micropipettes were made non-adhesive with sigmacote (Sigma) to prevent tissue adherence during aspiration. Random organoid samples were selected for measurements. In every experiment, the same pipette was used to measure the villi and crypts of the same organoids, and data was pooled from 3 independent experiments.

**Single cell RNA-Sequencing analysis.**

The single cell RNA-sequencing data has been previously published, and available at the Gene Expression Omnibus (GEO) under accession codes GSE115956 (*15*). Data analysis and single cell clustering and visualization used methods as described before (*15*). Implementation of the Rtsne R package used for tSNE projections is available at https://cran.r- project.org/web/packages/tsne/.

**Supplementary Text**

In this Supplementary Theory Note, we provide details on our physical model for the morphogenesis of intestinal organoids. The organoid is treated as a closed epithelial monolayer with two distinct regions, encapsulating an incompressible fluid lumen. We develop a three-dimensional biophysical model to study the mechanics of organoids, and use it to derive analytical results of specific morphologies, i.e. bulged and budded shapes, concentrating in particular on influence of crypt apical constriction and lumen volume changes on morphogenesis.

### 1. Two-region vertex model

The macroscopic shape of epithelial tissues and organs can be understood from mechanical interactions at the cellular level, such as cell-cell adhesion, actomyosin-mediated tension along the cell membrane, etc. Vertex models are a class of multiscale mechanical models to understand the interplay between cellular mechanical forces and tissue-scale deformation (*21*, *55*, *56*). In vertex models, a tissue is described as a set of vertices, where each vertex represents a tri-cellular junction where cell edges meet, and on which force balance is written (taking into account forces such as surface tensions, line tensions, internal fluid pressure, and external forces from surrounding environment).

An intestinal organoid is initially a spherical epithelial monolayer with a central luminal fluid cavity. After symmetric breaking which creates segregated stem cell and differentiated cell regions, the organoid will evolve towards pear-shaped configurations composed of two regions, crypt and villus. For simplicity, each region in the model is treated as a spherical cap. In the following, we first discuss the free energy of a single cell in the monolayer, then get the total energy of the whole organoid.

#### 1.1. Free energy of a single cell

Consider a single cell with three surface tensions $\Gamma_a$, $\Gamma_b$, and $\Gamma_l$, and three surface areas $A_a$, $A_b$, and $A_l$, where the subscripts $a$, $b$, and $l$ respectively represent apical, basal, and lateral surfaces/domains. Then, the free energy of a single cell is

$$f = \Gamma_a A_a + \Gamma_b A_b + \frac{1}{2}\Gamma_l A_l, \qquad (1)$$

The apical and basal surfaces are simplified as squares with side lengths $d_a$ and $d_b$ (although more complex shape would give identical results up to pre-factors), the height of a cell is $h$, so that the free energy (1) becomes

$$f=\Gamma_a d_a^2+\Gamma_a d_b^2+\Gamma_l h\left(d_a+d_b\right). \tag{2}$$

Each region is treated as a part of a homogeneous sphere shell, which has total cell number $N'$. In the spherical region, the side lengths are related to the region radii, i.e. $d_a=\sqrt{4\pi/N'}R_a$, $d_b=\sqrt{4\pi/N'}R_b$, where $R_a$ and $R_b$ are the inner (apical) and outer (basal) radii, respectively. Moreover, we have $R_a=R-h/2$, $R_b=R+h/2$, where $R$ is the neutral radius (see Fig. S2A for a schematic). Then, the free energy can be rewritten as

$$f=\frac{4\pi}{N'}\left[\left(\Gamma_a+\Gamma_b\right)R^2+\left(\Gamma_b-\Gamma_a\right)Rh\right]+2\sqrt{\frac{4\pi}{N'}}\cdot\Gamma_l Rh. \tag{3}$$

For simplicity, a thin-film assumption is employed, which means the thickness of the spherical sheet is much smaller than its radius, i.e. $\left(h/R\right)^2<<1$ ($R/h$ is typically 2 –10 in the early stages of bulging), which leads to $N'V_{e0}=4\pi R^2 h\left[1+\frac{1}{12}\left(\frac{h}{R}\right)^2\right]\approx 4\pi R^2 h$, where $V_{e0}$ is the cell volume. This greatly simplifies the analytics, as this yields $h\approx N'V_{e0}/\left(4\pi R^2\right)$. Given that cell volume is under osmotic regulation, involving stresses much larger than the ones produced by actomyosin, it is reasonable to assume that the volume $V_{e0}$ is independent from tension forces. However, cell volume may change during villus cell differentiation, due to active osmotic regulation, which will be discussed in Subsection 1.4. Under these assumptions, the free energy is only related to radius $R$:

$$f\left(R\right)\approx\frac{4\pi}{N'}\left(\Gamma_a+\Gamma_b\right)R^2+\left[\left(\Gamma_b-\Gamma_a\right)+2\Gamma_l\sqrt{\frac{N'}{4\pi}}\right]\frac{V_{e0}}{R}, \tag{4}$$

and the corresponding neutral radius in free state $\tilde{R}$ should satisfy $\left.\frac{\partial f}{\partial R}\right|_{\tilde{R}}=0$, which leads to

$$\tilde{R}=\sqrt{\frac{N'}{4\pi}}\left(\frac{V_{e0}\Gamma_l}{\Gamma_a+\Gamma_b}\right)^{\frac{1}{3}}\left(1+\frac{\Gamma_b-\Gamma_a}{2\Gamma_l}\sqrt{\frac{4\pi}{N'}}\right)^{\frac{1}{3}}. \tag{5}$$

Using Eq. (5), free energy (4) can be recast as

$$f \approx \frac{4\pi}{N'}\left(\Gamma_a + \Gamma_b\right) R^2 \left[1 + 2\left(\frac{\tilde{R}}{R}\right)^3\right]. \qquad (6)$$

Using Eq. (6) and introducing the deformation ratio $\lambda = R/\tilde{R}$, we can further get the free energy density $f/V_{e0} = \frac{4\pi(\Gamma_a+\Gamma_b)\tilde{R}^2}{N'V_{e0}}(\lambda^2+2\lambda^{-1})$, which indicates that $4\pi\left(\Gamma_a + \Gamma_b\right)^2 \tilde{R}^2 / \left(N'V_{e0}\right)$ acts as the stiffness of the spherical epithelium. For a large spherical monolayer ($N'$ is a large number), we can neglect the term of apico-basal difference in Eq. (5), and approximate the stiffness as $\left(\Gamma_a + \Gamma_b\right)^{1/3} \Gamma_l^{2/3} V_{e0}^{-1/3}$, emphasizing the crucial role for the sum of apical and basal tensions in setting in-plane resistance to deformations (which will become crucial to compare the respective responses of villus and crypt regions to lumen inflation, see Fig. 3 of the main text).

##### 1.2. Free energy of a two-region organoid epithelium

The free energy of the whole organoid is the sum of free energies in two regions. For simplicity, every cell in each region is assumed to be the same. Then, the free energy of a two-region epithelium is $F = N_c f_c + N_v f_v$, where $N_i$ and $f_i$ are respectively cell number and cellular free energy in region $i$, with the index $i = \mathrm{c}, \mathrm{v}$ denoting respectively crypt and villus. Using Eq. (6), the free energy of a single cell in region *i* is $f_i \approx \left(4\pi / N_i'\right)\left(\Gamma_a + \Gamma_b\right)_i R_i^2 \left[1 + 2\left(\tilde{R}_i / R_i\right)^3\right]$, and corresponding free energy of the whole epithelium yields

$$F \approx 4\pi\left(\Gamma_a + \Gamma_b\right)_c \frac{N_c}{N_c'} R_c^2 \left[1 + 2\left(\frac{\tilde{R}_c}{R_c}\right)^3\right] + 4\pi\left(\Gamma_a + \Gamma_b\right)_v \frac{N_v}{N_v'} R_v^2 \left[1 + 2\left(\frac{\tilde{R}_v}{R_v}\right)^3\right]. \quad (7)$$

A number of parameters in Eq. (7) can be eliminated as many geometric variables (such as $N_i$, $N_i'$, and $R_i$) are related. Firstly, we have organoid volume $V = V_c + V_v$, where $V_i = \pi R_i^3 \left(2 + 3\cos\theta_i - \cos^3\theta_i\right)/3$ is the volume of region *i*. For simplicity, we introduce an equivalent organoid radius $R_t$ satisfying $V = 4\pi R_t^3 / 3$, and considering the geometric relation $R_c \sin\theta_c = R_v \sin\theta_v$, then the region radius $R_i$ is related to radius $R_t$ and polar angles $\theta_i$ (see Fig. S2B for schematic) by

$$R_i = R_t g_i^{-1/3}, \qquad (8)$$

with

$$
\begin{aligned}
g_\mathrm{c} &= \frac{1}{2}\left[\left(1+\frac{3}{2}\cos\theta_\mathrm{c}-\frac{1}{2}\cos^3\theta_\mathrm{c}\right)+\left(\frac{\sin\theta_\mathrm{c}}{\sin\theta_\mathrm{v}}\right)^3\left(1+\frac{3}{2}\cos\theta_\mathrm{v}-\frac{1}{2}\cos^3\theta_\mathrm{v}\right)\right] \\
g_\mathrm{v} &= \left(\frac{\sin\theta_\mathrm{c}}{\sin\theta_\mathrm{v}}\right)^{-3} g_\mathrm{c}
\end{aligned}
. \qquad (9)
$$

Secondly, considering cells in one region have the same geometric shape, the ratio of cell number in the region (which is a spherical cap) to that in the whole spherical shell is proportional to the ratio of surface areas, that is $N_i / N_i' = A_i / A_i'$ , where the surface area of region $i$ is $A_i = \pi R_i^2 \left(2+2\cos\theta_i\right)$, and the surface are of corresponding spherical shell is $A_i' = 4\pi R_i^2$. Then we can get

$$
\frac{N_i}{N_i'} = \frac{1}{4} s_i, \qquad (10)
$$

where $s_i\left(\theta_i\right) = 2+2\cos\theta_i$.

An intestinal organoid evolves from an initial spherical shape toward a two-region configuration. Crypt apical constriction is found to initiate intestinal morphogenesis *in vivo*, and apical surface areas of crypt cells also evolve during the development of intestinal organoids (Fig. 1B). In view of these, we consider that tensions in crypt cells may be distinct from those of villus cells, and evaluate the role of crypt mechanics in organoid morphogenesis. Given that intestinal organoid initially contains identical cell types, prior to the symmetry breaking of fate (*15*), we take all cells to initially have the same surface tensions. For simplicity, we assume that there is no apical-basal tension difference for an initial spherical organoid, and further assume that lateral tensions are unchanged everywhere during development, i.e. $\Gamma_{l\mathrm{c}} = \Gamma_{l\mathrm{v}} = \Gamma_l$. Then, we can non-dimensionalize Eq. (7) by introducing four dimensionless parameters:

- relative region size of the crypt $\varphi = N_\mathrm{c}/N_\mathrm{t}\left(N_\mathrm{t} = N_\mathrm{c}+N_\mathrm{v}\right)$,
- in-plane contraction ratio $\alpha = \left(\Gamma_a+\Gamma_b\right)_\mathrm{c} / \left(\Gamma_a+\Gamma_b\right)_0$, which quantifies the relative changes in crypt stiffness due to changes of apical/basal tensions.
- normalized organoid radius $\beta = R_\mathrm{t} / \tilde{R}_0$, where $\tilde{R}_0$ is the radius of the initial spherical organoid in free state,
- normalized apico-basal tension difference $\gamma_\mathrm{c} = \frac{1}{2}\left(\frac{\Gamma_b-\Gamma_a}{\Gamma_l}\right)_\mathrm{c}\sqrt{\frac{4\pi}{N_\mathrm{t}}}$, which causes the crypt to

have a spontaneous curvature.

Submitting Eqs. (8) and (10) into Eq. (7), the dimensionless free energy $\hat{F} = F\big/\left[\pi\left(\Gamma_a+\Gamma_b\right)_0 \tilde{R}_0^2\right]$ becomes

$$\hat{F} \approx \alpha\beta^2 \cdot s_{\mathrm{c}} g_{\mathrm{c}}^{-2/3}\left[1+\frac{2}{\beta^3}\left(\frac{\tilde{R}_{\mathrm{c}}}{\tilde{R}_0}\right)^3 g_{\mathrm{c}}\right]+\beta^2 \cdot s_{\mathrm{v}} g_{\mathrm{v}}^{-2/3}\left[1+\frac{2}{\beta^3}\left(\frac{\tilde{R}_{\mathrm{v}}}{\tilde{R}_0}\right)^3 g_{\mathrm{v}}\right], \quad (11)$$

where $\left(\tilde{R}_{\mathrm{c}}/\tilde{R}_0\right)^3 = 8\alpha^{-1}\varphi^{3/2}s_{\mathrm{c}}^{-3/2}\left(1+\varphi^{-1/2}s_{\mathrm{c}}^{1/2}\gamma_{\mathrm{c}}/2\right)$, $\left(\tilde{R}_{\mathrm{v}}/\tilde{R}_0\right)^3 = 8\left(1-\varphi\right)^{3/2}s_{\mathrm{v}}^{-3/2}$.

To simplify these expression, we redefine geometric parameters $G_{\mathrm{c}}\left(\theta_{\mathrm{c}},\theta_{\mathrm{v}}\right)=s_{\mathrm{c}}^{-3/2}g_{\mathrm{c}}$, $G_{\mathrm{v}}\left(\theta_{\mathrm{c}},\theta_{\mathrm{v}}\right)=s_{\mathrm{v}}^{-3/2}g_{\mathrm{v}}$ (which quantify the degree of opening of villus and crypt regions), and introduce the normalized volume $v=\beta^3$. The free energy then reads

$$\hat{F} \approx v^{2/3}\left(\alpha G_{\mathrm{c}}^{-2/3}+G_{\mathrm{v}}^{-2/3}\right)+16v^{-1/3}\left[\varphi^{3/2}G_{\mathrm{c}}^{1/3}+\left(1-\varphi\right)^{3/2}G_{\mathrm{v}}^{1/3}+\frac{1}{2}\varphi g_{\mathrm{c}}^{1/3}\gamma_{\mathrm{c}}\right]. \quad (12)$$

Eq. (12) shows that $\hat{F}$ is a function of only two parameters, i.e. the polar angles $\theta_{\mathrm{c}}$ and $\theta_{\mathrm{v}}$, with the minima of $\hat{F}$ (and corresponding $\theta_{\mathrm{c}}$ and $\theta_{\mathrm{v}}$) determining the shape of organoids at mechanical equilibrium. In principle, in-plane contraction ($\alpha$), spontaneous curvature ($\gamma_{\mathrm{c}}$), lumen volume ($v$), and crypt size ($\varphi$) can all affect organoid morphogenesis, and we first sequentially explored the influence of each of these parameters separately, to gain intuitive insights into their influence on morphology, which can then be verified in experimental data. Finally, to avoid non-physical minima of this energy, we employed a penalty function to guarantee that the inner radii of crypt and villus are always positive, i.e. $R_{ai}=R_i-h_i/2>0$. In the calculation, we use $\exp\left\{\eta\left[\left(v/g_i\right)^{1/3}-\left(2\varphi/\tilde{\kappa}_0\right)\left(v/G_i\right)^{-2/3}\right]\right\}$ as a penalty function, where $\eta$ is chosen as $-10^5$, $\tilde{\kappa}_0=4\pi\tilde{R}_0^3/\left(N_{\mathrm{t}}V_{\mathrm{e0}}\right)$ is a shape factor that characterizes the initial volume ratio between the whole organoid and the epithelial monolayer.

##### 1.2.1. Organoid morphologies

We first study the organoid morphologies with varied volume $v$ and spontaneous curvature $\gamma_{\mathrm{c}}$ (of crypt region), with $\alpha=1$ (equal in-plane contraction in villus and crypt regions) . Setting

$\alpha = 1$ and crypt size $\varphi = 0.2$, the phase diagram in Fig. S2D not only highlights the influence of spontaneous curvature, but also intuitively reveals that the inflation of organoids tends to reopen both the crypt and villus and recover the original spherical shape. In other words, transformation from a budded shape to a bulged one may happen during organoid inflation. This is consistent with classical theoretical result on lipid vesicles with regions of spontaneous curvature, which shows that an increase in vesicle volume will reverse the budding induced by spontaneous curvature (*32*) . Examining organoid morphology with $\gamma_{\mathrm{c}} = -0.25$ in the first graph of Fig. S2D as an example, its crypt is fully closed under moderate volume expansion, but will open up when the lumen volume increases above a critical threshold. We employed the "degree of crypt opening", defined as $\theta_{\mathrm{c}} / \left(\pi - \theta_{\mathrm{v}}\right)$, to quantify the morphogenesis of intestinal organoid. This parameter ranges from 0 to 1, where 0 corresponds to the budded shape with crypt and villus fully closed and 1 to a fully spherical organoid shape.

As shown in Fig. S2D, the in-plane contraction in crypt also affects the organoid morphology. For an organoid with weak in-plane crypt contraction ($\alpha < 1$), the original spherical shape is recovered by lumen volume expansion, while the recovery is harder when the crypt has strong in-plane contraction ($\alpha > 1$). Strikingly, we find that a crypt with a large enough spontaneous curvature may not open up even for arbitrarily large increases in lumen volume. This indicates critical mechanical forces in crypt may exist, beyond which the shape transformation back to spherical shapes never happens.

##### 1.2.2. Morphometric parameters

Upon organoid swelling, the crypt and villus sustain distinct in-plane and out-of-plane deformations, which respectively modulate the thickness and radius of each region. In other words, these geometric quantities can be employed as morphometric parameters to evaluate the mechanical deformations (and corresponding cell tensions) in two regions. For example, profiles of epithelial thickness and radius have been proposed as metrics to infer the nature of forces driving epithelial folds in epithelium-stroma structures (*57*). We thus examine thickness ratio $h_{\mathrm{c}} / h_{\mathrm{v}}$ and radius ratio $R_{\mathrm{c}} / R_{\mathrm{v}}$ to further quantify the morphological evolution during volume expansion. We find in particular that their dependence on two mechanical parameters, i.e. in-plane contraction $\alpha$ and spontaneous curvature $\gamma_{\mathrm{c}}$, is qualitatively different (Fig. S2E-H). The thickness (or radius) ratio shows two distinct trends during organoid inflation. For an organoid with $\alpha = 1$, $\gamma_c = -0.25$, the thickness ratio increases almost linearly with volume expansion at the early stage, bur drops abruptly at $v \approx 2$, while its radius ratio also undergoes both linear and nonlinear variations, but in an opposite way (Fig. S2F). These abrupt transitions of thickness and radius ratios are due to shape transformation of organoids (shown in Fig. S2D), and clearly indicate that, for organoids with different morphologies, the thickness (or radius) ratio is modulated by lumen volume in distinct ways. Furthermore, we find that crypts with strong in-plane contraction (i.e. $\alpha > 1$) are always thicker than villi (Fig. S2E,G), while crypts with $\alpha < 1$ is usually thinner than villi (Fig. S2E, H). This is intuitive as hydrostatic pressure is uniform within the organoid lumen, so that stiffer regions deform less than softer ones (resulting in less thinning). We also find that the inflation of organoids tends to widen the thickness difference between two regions (Fig. S2E, G, H), as the softer region tends to accommodate the bulk of the pressure-induced deformation.

Furthermore, as already shown in Fig. 2B, spontaneous curvature $\gamma_c$ always tends to increase the crypt thickness. This is consistent with classical results in *Drosophila* gastrulation, where ventral cells are lengthened during furrow formation (*58*). However, Fig. S2 F-H further indicate that, for a swelling organoid, the influence of $\gamma_c$ on the thickness ratio $h_c / h_v$ is negligible when the spontaneous curvature is not large enough to close the crypt (which belongs to a budded shape). In other words, the thickness ratio of a swelling organoid with a partially open crypt (e.g., a bulged organoid) is almost independent on $\gamma_c$, although increasing crypt apical tension can influence thickness ratio by increasing $\alpha$.

##### 1.3. Line tension in neck zone

So far, we have only considered changes in the bulk properties of each organoid region, such as in-plane contractions and spontaneous curvatures. However, mechanical forces at the boundary between these two regions may also drive the morphological evolution in biological systems (*59-61*). Here, we assume cells in the neck zone (connection part of crypt and villus) carry distinct surface tensions (and hence the free energy) with cells in two regions. Since the neck zone of organoid is rather narrow, and more like a hollow cylinder rather than a spherical shell, it is reasonable to model the neck zone as a short cylindrical monolayer and neglect its volume contribution to organoid.

Considering neck cells with longitudinal side length $e$, height $h$, and radial side lengths in the apical and basal surfaces $d_a$ and $d_b$ (see Fig. S2C for schematic), then the free energy (1) becomes $f = \Gamma_a ed_a + \Gamma_a ed_b + \Gamma_l eh + \frac{1}{2}\Gamma_l h\left(d_a + d_b\right)$. The geometric relationship of a single cell and a cylindrical epithelium can be described by $d_a = 2\pi R_a / N_{\mathrm{r}}$, $d_b = 2\pi R_b / N_{\mathrm{r}}$, where $N_{\mathrm{r}}$ is the cell number in the radial direction. Letting $R$ be the neutral radius of the cylindrical epithelium, we obtain $h = N_{\mathrm{r}} V_{\mathrm{e0}} / \left(2\pi eR\right)$, which recasts the free energy as

$$f = \frac{2\pi}{N_{\mathrm{r}}}\left(\Gamma_a + \Gamma_b\right)eR + \left[\frac{1}{2}\left(\Gamma_b - \Gamma_a\right) + \Gamma_l \frac{N_{\mathrm{r}}}{2\pi}\right]\frac{V_{\mathrm{e0}}}{R} + \Gamma_l \frac{V_{\mathrm{e0}}}{e}. \quad (13)$$

Eq. (13) indicates that the free energy depends on two geometric variables $R$ and $e$, i.e. $f = f\left(R, e\right)$. Considering the free state of cells, which satisfies $\partial f / \partial R = 0$, $\partial f / \partial e = 0$, we can get radius $\tilde{R}$ and length $\tilde{e}$ in the free state

$$\tilde{R} = \frac{N_{\mathrm{r}}}{2\pi}\left(\frac{V_{\mathrm{e0}}\Gamma_l}{\Gamma_a + \Gamma_b}\right)^{\frac{1}{3}}\left(1 + \frac{\Gamma_b - \Gamma_a}{2\Gamma_l}\frac{2\pi}{N_{\mathrm{r}}}\right)^{\frac{2}{3}}, \quad \tilde{e} = \left(\frac{V_{\mathrm{e0}}\Gamma_l}{\Gamma_a + \Gamma_b}\right)^{\frac{1}{3}}\left(1 + \frac{\Gamma_b - \Gamma_a}{2\Gamma_l}\frac{2\pi}{N_{\mathrm{r}}}\right)^{-\frac{1}{3}}. \quad (14)$$

Using Eq. (14), the free energy of a cell in the neck can finally be expressed as

$$f = \frac{2\pi}{N_{\mathrm{r}}}\left(\Gamma_a + \Gamma_b\right)\left[eR + \tilde{e}\tilde{R}\left(\frac{\tilde{R}}{R} + \frac{\tilde{e}}{e}\right)\right]. \quad (15)$$

For an organoid with two regions (crypt and villus) and a neck zone, the total free energy are contributed by three parts, i.e. $F = N_{\mathrm{c}} f_{\mathrm{c}} + N_{\mathrm{v}} f_{\mathrm{v}} + N_{\mathrm{n}} f_{\mathrm{n}}$, where $N_{\mathrm{n}}$ and $f_{\mathrm{n}}$ are respectively the cell number and cellular free energy in the neck zone ($f_{\mathrm{n}}$ follows the expression in Eq. (15)).

Since the neck zone is mainly constrained by regions in its radial direction, we assume a stress-free state in the longitudinal direction, i.e. $\partial f / \partial e = 0$, which lead to $e = \tilde{e}\sqrt{\tilde{R}_{\mathrm{n}} / R_{\mathrm{n}}}$. Then the free energy of neck zone yields

$$F_{\mathrm{n}} = N_{\mathrm{n}} f_{\mathrm{n}} = 2\pi N_{\mathrm{e}}\left(\Gamma_a + \Gamma_b\right)_{\mathrm{n}} \tilde{e}\sqrt{\tilde{R}_{\mathrm{n}} R_{\mathrm{n}}}\left[2 + \left(\frac{\tilde{R}_{\mathrm{n}}}{R_{\mathrm{n}}}\right)^{3/2}\right], \quad (16)$$

where $N_{\mathrm{e}}$ is the cell number in the longitudinal direction (therefore we have $N_{\mathrm{n}} = N_{\mathrm{r}} N_{\mathrm{e}}$).

The in-plane contraction ratio $\Lambda = \left(\Gamma_a + \Gamma_b\right)_\mathrm{n} / \left(\Gamma_a + \Gamma_b\right)_0$ is introduced to characterize the 'line tension' between two regions. The geometric relationship $R_\mathrm{n} = R_\mathrm{c} \sin\theta_\mathrm{c}$ implies $R_\mathrm{n} = R_\mathrm{t} g_\mathrm{n}^{-1/3}$ with $g_\mathrm{n} = g_\mathrm{c} / \sin^3\theta_\mathrm{c}$. Then, we have $\hat{F}_\mathrm{n} = F_\mathrm{n} \big/ \left[\pi \left(\Gamma_a + \Gamma_b\right)_0 \tilde{R}_0^2\right]$ to be $2N_\mathrm{e}\Lambda\tilde{e}\tilde{R}_\mathrm{n}^{1/2} R_0^{-3/2}\left[2g_\mathrm{n}^{-1/6}\beta^{1/2} + g_\mathrm{n}^{1/3}\beta^{-1}\left(\tilde{R}_\mathrm{n}/\tilde{R}_0\right)^{3/2}\right]$. To further simplify $\hat{F}_\mathrm{n}$, we still need to determine $N_\mathrm{r}$, which affects both $\tilde{e}$ and $\tilde{R}_\mathrm{n}$. The radial cell number of the neck depends on the total cell number of organoid $N_\mathrm{t}$ and the position of neck (dominated by crypt size $\varphi$), that is $N_\mathrm{r}=N_\mathrm{r}\left(\varphi, N_\mathrm{t}\right)$. Specific expression of $N_\mathrm{r}$ can be estimated as follows: A narrow neck in an spherical organoid in free state satisfies $N_\mathrm{r}\tilde{d}=2\pi\tilde{R}_0\sin\theta_\mathrm{n}$, where $\theta_\mathrm{n}$ is the polar angle of neck, $\tilde{d} = \sqrt{4\pi/N_\mathrm{t}}\tilde{R}_0$ is the side length of a single cell. Further considering the geometric relation $\varphi = 2\pi\tilde{R}_0^2\left(1-\cos\theta_\mathrm{n}\right)/\left(4\pi\tilde{R}_0^2\right) = \left(1-\cos\theta_\mathrm{n}\right)/2$, we can get $N_\mathrm{r}=\sqrt{4\pi N_\mathrm{t}}\cdot\Delta$, where $\Delta = \sqrt{\varphi-\varphi^2}$. To focus on the in-plane contraction in the neck, the difference of apical and basal tensions (i.e. spontaneous curvature) is neglected, which finally leads to a simplified free energy of the neck

$$\hat{F}_\mathrm{n}=8N_\mathrm{e}\sqrt{\frac{2\pi}{N_\mathrm{t}}}\left(\Lambda^{1/2}\Delta^{1/2}\beta^{1/2}g_\mathrm{n}^{-1/6} + \sqrt{2}\Delta^2\beta^{-1}g_\mathrm{n}^{1/3}\right). \quad (17)$$

By adding free energy (17) into Eq. (12), we can evaluate the influence of the overall line tension, arising from the in-plane contraction of cells in the neck, on organoid morphogenesis. Fig. 2B shows that, although a contractile neck can promote the bulging and budding of organoids (i.e. decreased radius ratio $R_\mathrm{c}/R_\mathrm{v}$), it has negligible effects on the thickness ratio $h_\mathrm{c}/h_\mathrm{v}$. This is in contrast with our experimental findings (Fig. 2C) where bulging of organoids is robustly accompanied by thickness increases on the crypts compared to villi. This implies that the line tension in neck is not the major driving force for crypt bulging.

##### 1.4. Cell volumes and villus mechanics

The model in Subsection 1.2 considers the influence of crypt mechanics and lumen volume on morphogenesis. However, mechanical contributions from the villus could also impact intestinal organoid development. For example, in the late stage of organoid morphogenesis, the villus shows both cell swelling (Fig. 4B) and increased intensity of basal myosin (Fig. 4G), which might result in elevated basal tensions. To explore this, we extended the previous model, which assumes a constant cell volume in both regions and constant cell tensions in villus during morphogenesis, to incorporate potential variations in cell volumes and villus tensions. We thus introduce normalized cell volumes $v_{\mathrm{ec}} = V_{\mathrm{ec}} / V_{\mathrm{e0}}$, $v_{\mathrm{ev}} = V_{\mathrm{ev}} / V_{\mathrm{e0}}$, where $V_{\mathrm{ec}}$ and $V_{\mathrm{ev}}$ are respectively the volumes of a crypt cell and a villus cell. In analogy to the definitions in crypt mechanics, in-plane contraction ratio $\alpha_{\mathrm{v}} = \left(\Gamma_a + \Gamma_b\right)_{\mathrm{v}} \big/ \left(\Gamma_a + \Gamma_b\right)_0$, and spontaneous curvature $\gamma_{\mathrm{v}} = \frac{1}{2}\left(\frac{\Gamma_b - \Gamma_a}{\Gamma_l}\right)_{\mathrm{v}} \sqrt{\frac{4\pi}{N_{\mathrm{t}}}}$ are introduced to examine the effects of villus tensions. With these extensions of the model, this rescaled organoid energy $\hat{F}$ now reads:

$$\begin{aligned}\hat{F} = v^{2/3}\left(\alpha_{\mathrm{c}} G_{\mathrm{c}}^{-2/3} + \alpha_{\mathrm{v}} G_{\mathrm{v}}^{-2/3}\right) + 16 v^{-1/3}\left[\varphi^{3/2} v_{\mathrm{ec}} G_{\mathrm{c}}^{1/3} + \left(1-\varphi\right)^{3/2} v_{\mathrm{ev}} G_{\mathrm{v}}^{1/3}\right] \\ + 8 v^{-1/3}\left[\varphi v_{\mathrm{ec}} g_{\mathrm{c}}^{1/3} \gamma_{\mathrm{c}} + \left(1-\varphi\right) v_{\mathrm{ev}} g_{\mathrm{v}}^{1/3} \gamma_{\mathrm{v}}\right]\end{aligned}. \quad (18)$$

##### 1.4.1. Influence of cell swelling on morphogenesis

We first evaluate the dependence of organoid morphologies on cell swelling in either crypt or villus. As shown in Fig. S2I, both the swelling of crypt cells and villus cells can promote budding. Furthermore, crypt size $\varphi$ impacts the efficiency of cell swelling on budding. Given the fact that the villus is usually much larger than the crypt, swelling of villus cells is more efficient to promote budding. Furthermore, even when both regions have an equal size, the cell swelling in villus is still more efficient. For a crypt undergoing both cell swelling and tension-modulated deformations, the in-plane contraction will limit the extension of crypt region, while the crypt bending will be hindered by cell swelling. Overall, cell swelling is less efficient on budding when it happens in the tension-enhanced region (i.e. the crypt) than in the normal region (i.e. the villus). Besides, Fig. S2J shows that the effect of cell swelling on budding can be eliminated by lumen expansion. This is different from the influence of crypt mechanics, which leads to maintained closure of the crypt even under infinite lumen expansion.

##### 1.4.2. Influence of villus mechanics on morphogenesis

We then examine the influence of spontaneous curvature of villus $\gamma_{\mathrm{v}}$ on organoid morphology. Unlike spontaneous curvature $\gamma_{\mathrm{c}}$, which is negative due to the enhanced apical tension in crypt, spontaneous curvature $\gamma_{\mathrm{v}}$ is chosen to be positive in Fig. S2K-L, in light of the elevated basal myosin accumulation observed in villus (Fig. 2G) as well as basal constriction observed in wild-type cells next to cells with reduced Myosin levels (Fig. S5E). Interestingly, the spontaneous curvature $\gamma_{\mathrm{v}}$ will promote the opening of two regions only when $\gamma_{\mathrm{c}}$ is quite small ( $|\gamma_{\mathrm{c}}| < 0.05$ or estimated value in initial bulging phase), while the out-of-plane bending of villus will facilitate the closure of two regions when the crypt engenders notable spontaneous curvature and strong in-plane contraction (Fig. S2K). Importantly, the dependence of thickness (or radius) ratio on $\gamma_{\mathrm{v}}$ is negligible for an organoid with either equal or stronger in-plane contraction in crypt than in villus (Fig. S2L), which we show from Fig. 3 is the relevant case for us. This argues that although basal enrichment of Myosin in the villus region is expected to help and contribute to bulging and budding, it cannot be the dominant/sole driving force (otherwise in-plane contraction of villi would be larger than crypts and lumen inflation would cause crypt dilation), so that we neglect $\gamma_{\mathrm{v}}$ in first approximation for the fits discussed in Section 4.

#### 2. Analytic approximations

Experimentally, crypt regions are much smaller than villus regions, in particular during the first phases of bulging/budding which we explore here. Based on this, we can simplify the model by considering $V_{\mathrm{c}} \ll V_{\mathrm{v}}$ and $\theta_{\mathrm{v}} \to 0$. The volumetric relation $V = V_{\mathrm{c}} + V_{\mathrm{v}}$ can be expressed as $R_{\mathrm{t}}^3 \approx p_{\mathrm{c}} R_{\mathrm{c}}^3 + R_{\mathrm{v}}^3$, where $p_{\mathrm{c}} = \left(2 + 3\cos\theta_{\mathrm{c}} - \cos^3\theta_{\mathrm{c}}\right)/4$. Considering $V_{\mathrm{c}} \ll V_{\mathrm{v}}$ (or $p_{\mathrm{c}} R_{\mathrm{c}}^3 \ll R_{\mathrm{v}}^3$ ) leads to $R_{\mathrm{v}} \approx R_{\mathrm{t}}\left[1 - (p_{\mathrm{c}}/3)(R_{\mathrm{c}}/R_{\mathrm{t}})^3\right]$. Combined with Eq. (10), free energy (7) can be rewritten as

$$\begin{aligned} F \approx\ & \pi\left(\Gamma_a + \Gamma_b\right)_{\mathrm{c}} s_{\mathrm{c}} R_{\mathrm{c}}^2 \left[1 + 2\left(\frac{\tilde{R}_{\mathrm{c}}}{R_{\mathrm{c}}}\right)^3\right] \\ & + \pi\left(\Gamma_a + \Gamma_b\right)_{\mathrm{v}} s_{\mathrm{v}} R_{\mathrm{t}}^2 \left\{1 + 2\left(\frac{\tilde{R}_{\mathrm{v}}}{R_{\mathrm{t}}}\right)^3 + \frac{2}{3}\left[\left(\frac{\tilde{R}_{\mathrm{v}}}{R_{\mathrm{t}}}\right)^3 - 1\right] p_{\mathrm{c}} \left(\frac{R_{\mathrm{c}}}{R_{\mathrm{t}}}\right)^3\right\} \end{aligned} . \qquad (19)$$

Letting $\beta_{\mathrm{c}} = R_{\mathrm{c}} / \tilde{R}_0$ be the normalized crypt radius, one obtains

$$\begin{aligned} \hat{F} \approx\ & \alpha s_{\mathrm{c}} \beta_{\mathrm{c}}^2 + \left(16\varphi^{3/2} s_{\mathrm{c}}^{-1/2} + 8\varphi\gamma_{\mathrm{c}}\right)\beta_{\mathrm{c}}^{-1} + s_{\mathrm{v}}\beta^2 + 16(1-\varphi)^{3/2} s_{\mathrm{v}}^{-1/2}\beta^{-1} \\ & + \frac{2}{3}\left[8(1-\varphi)^{3/2} s_{\mathrm{v}}^{-1/2}\beta^{-3} - s_{\mathrm{v}}\right] p_{\mathrm{c}} \beta_{\mathrm{c}}^3 \beta^{-1} \end{aligned} , \qquad (20)$$

For a small $\theta_{\mathrm{v}}$, using $R_{\mathrm{c}}\sin\theta_{\mathrm{c}} = R_{\mathrm{v}}\sin\theta_{\mathrm{v}}$ ,we have $\theta_{\mathrm{v}} \approx \left(R_{\mathrm{c}} / R_{\mathrm{v}}\right)\sin\theta_{\mathrm{c}}$ ,which leads to $s_{\mathrm{v}} \approx 4-\left(\beta_{\mathrm{c}} / \beta\right)^2 \sin\theta_{\mathrm{c}}^2$. With these approximations, the free energy $\hat{F}$ only depends on $\theta_{\mathrm{c}}$ and $\beta_{\mathrm{c}}$:

$$\begin{aligned}\hat{F} \approx\ & 4\beta^2 + 8\left(1-\varphi\right)^{3/2}\beta^{-1} + \alpha s_{\mathrm{c}}\beta_{\mathrm{c}}^2 + \left(16\varphi^{3/2}s_{\mathrm{c}}^{-1/2} + 8\varphi\gamma_{\mathrm{c}}\right)\beta_{\mathrm{c}}^{-1} \\ & + \left[\left(1-\varphi\right)^{3/2}\beta^{-3} - 1\right]\left(\beta_{\mathrm{c}}^2\sin^2\theta_{\mathrm{c}} + \frac{8}{3}p_{\mathrm{c}}\beta_{\mathrm{c}}^3\beta^{-1}\right)\end{aligned} \quad . \qquad (21)$$

In the following, based on this simplified free energy (21), we will analyze specific organoid morphologies and get corresponding analytical expressions of morphometric parameters. As a limiting case, crypt morphologies under infinite organoid expansion will be discussed. Besides, the influence of cell volumes will be explicitly explored with analytic formulation.

##### 2.1. Scaling laws for thickness and radius modulations

As aforementioned, after the initial symmetric breaking event, an intestinal organoid will evolve towards non-spherical configurations. The organoid first undergoes a bulging phase with the crypt gradually bulges out, then enters into a budded phase. Here, we focus on these two typical morphologies, the bulged shape and the budded one, during the development of intestinal organoids, and make use of their shape features to further simplify the free energy shown in Eq. (21) and get analytical expressions of the radius ratio $R_{\mathrm{c}} / R_{\mathrm{v}}$ and the thickness ratio $h_{\mathrm{c}} / h_{\mathrm{v}}$.

###### 2.1.1. Bulging

In the bulging stage, the crypt just begins to form and is rather small (i.e., $\varphi$ is small). These indicate $\theta_* = \pi - \theta_{\mathrm{c}} \sim \varphi^{1/2}$ can be served as a small parameter (i.e., $\theta_* \to 0$), and functions of $\theta_{\mathrm{c}}$ in Eq. (21) can be approximated as $s_{\mathrm{c}} \approx \theta_*^2$, $\sin^2\theta_{\mathrm{c}} \approx \theta_*^2$, and $p_{\mathrm{c}} \approx 0$.Then, the free energy (21) is simplified as

$$\hat{F} \approx 4\beta^2 + 8\beta^{-1} + \left(\alpha - 1 + \beta^{-3}\right)\bar{\theta}_*^2 + 16\varphi^{3/2}\bar{\theta}_*^{-1} + 8\varphi\gamma_{\mathrm{c}}\beta_{\mathrm{c}}^{-1}, \qquad (22)$$

where $\bar{\theta}_* = \beta_{\mathrm{c}}\theta_*$, and the radius ratio and thickness ratio are respectively approximated as $R_{\mathrm{c}} / R_{\mathrm{v}} \approx \beta_{\mathrm{c}}/\beta$ and $h_{\mathrm{c}} / h_{\mathrm{v}} \approx 4\varphi\beta^2\bar{\theta}_*^{-2}$. One obtains $\bar{\theta}_* = 2\varphi^{1/2}\left(\alpha - 1 + \beta^{-3}\right)^{-1/3}$ from $\partial\hat{F} / \partial\bar{\theta}_* = 0$. To get an estimate of the normalized crypt radius $\beta_{\mathrm{c}}$, we need to expand the functions of $\theta_{\mathrm{c}}$ ( or

$\theta_*$ ) in Eq. (21) to a higher order $O\left(\theta_*^4\right)$, i.e. $s_{\mathrm{c}} \approx \theta_*^2 - \theta_*^4/12$, $\sin^2\theta_{\mathrm{c}} \approx \theta_*^2 - \theta_*^4/3$, and $p_{\mathrm{c}} \approx 3\theta_*^4/16$, which yield an additional sequence of terms in free energy (22) as $\left(2/3\right)\varphi^{3/2}\beta_{\mathrm{c}}^{-2}\bar{\theta}_* + \bar{\theta}_*^4\left\{-\alpha\beta_{\mathrm{c}}^{-2}/12 + \left(\beta^{-3}-1\right)\left(-\beta_{\mathrm{c}}^{-2}/3 + \beta_{\mathrm{c}}^{-1}\beta^{-1}/2\right)\right\}$. Using the extended free energy and considering $\partial\hat{F}/\partial\beta_{\mathrm{c}} = 0$ yield $\beta_{\mathrm{c}}^{-1} = \beta^{-1} - 16\varphi\gamma_{\mathrm{c}}\bar{\theta}_*^{-4}\Big/\left(1-\beta^{-3}\right)$. Thus, the radius ratio and thickness ratio of a bulged organoid can be finally estimated as

$$\frac{R_{\mathrm{v}}}{R_{\mathrm{c}}} \approx 1 - \varphi^{-1}\gamma_{\mathrm{c}}\frac{\left[\left(\alpha-1\right)v+1\right]^{4/3}}{v-1}, \frac{h_{\mathrm{c}}}{h_{\mathrm{v}}} \approx \left[\left(\alpha-1\right)v+1\right]^{2/3}. \tag{23}$$

Eq. (23) thus predicts that the thickness ratio depends only, at first order, on the in-plane contraction ratio $\alpha$. We found excellent agreement between numerical solutions of the full model, and the analytical criteria of Eq. (23), and confirmed in particular that the thickness ratio depends crucially on $\alpha$, while it is almost independent on $\gamma_{\mathrm{c}}$ (Fig. S4A). Furthermore, the radius ratio of a bulged organoid is expected to depend on $\varphi^{-1}\gamma_{\mathrm{c}}$ and $\alpha$ from Eq. (23). In a bulging crypt, the apical actomyosin accumulation is initially small (Fig. 2C), so that it is expected to engender weak in-plane contraction and out-of-plane bending, and corresponding mechanical parameters $\alpha - 1$ and $\gamma_{\mathrm{c}}$ can both be considered small. However, Eq. (23) indicates that the radius ratio is less dependent on $\alpha - 1$ than $\gamma_{\mathrm{c}}$, and the only leading parameter of $R_{\mathrm{c}}/R_{\mathrm{v}}$ is $\varphi^{-1}\gamma_{\mathrm{c}}$. As verified in Fig. S4A, the crypt size $\varphi$ and spontaneous curvature $\gamma_{\mathrm{c}}$ are indeed combined to affect the crypt radius, and the resulting parameter $\varphi^{-1}\gamma_{\mathrm{c}}$ can modulate $R_{\mathrm{c}}/R_{\mathrm{v}}$. Besides, Eq. (23) can fit well with the numerical results of a bulged organoid with varying volumes (Fig. S4A).

##### 2.1.2. Budding

For a budded organoid (which is equivalent to a near-closed organoid in our simplified spherical region models), we can take the converse limit of small $\theta_{\mathrm{c}}$ (i.e. $\theta_{\mathrm{c}} \to 0$), which results in $s_{\mathrm{c}} \approx 4$, $\sin^2\theta_{\mathrm{c}} \approx 0$, and $p_{\mathrm{c}} \approx 1$. Then the full expression of free energy (21) reduces to

$$\hat{F} \approx 4\beta^2 + 8\left(1-\varphi\right)^{3/2}\beta^{-1} + 4\alpha\beta_{\mathrm{c}}^2 + \left(8\varphi^{3/2} + 8\varphi\gamma_{\mathrm{c}}\right)\beta_{\mathrm{c}}^{-1} + \frac{8}{3}\left[\left(1-\varphi\right)^{3/2}\beta^{-3} - 1\right]\beta_{\mathrm{c}}^3\beta^{-1}, \tag{24}$$

which only depends on the normalized crypt radius $\beta_c$. Minimizing this energy with respect to the crypt radius (i.e. $\partial\hat{F}/\partial\beta_c = 0$) leads to $\beta_c\beta^{-1} = \left[\alpha - \left(\varphi^{3/2} + \varphi\gamma_c\right)\beta_c^{-3}\right] \Big/ \left[1 - (1-\varphi)^{3/2}\beta^{-3}\right]$, which can be recast as

$$1 - \left(\frac{R_c}{\tilde{R}_c}\right)^{-3} = \alpha^{-1}\frac{R_c}{R_t}\left[1 - (1-\varphi)^{3/2} v^{-1}\right]. \quad (25)$$

Experiments indicate that a budding organoid undergoes sustaining apical actomyosin accumulation in the crypt, which will lead to an enhanced in-plane contraction, i.e. $\alpha > 1$. Besides, considering the crypt volume is usually much smaller than the overall volume of the organoid, i.e. $V_c << V$, where $V_c = 4\pi R_c^3/3$ and $V = 4\pi R_t^3/3$ for a budded organoid, we can find that $R_c/R_t < 1$ always holds. Thus, the value of the right side of Eq. (25) is usually close to 0, which indicates that $R_c/\tilde{R}_c \approx 1$ (i.e. $R_c \approx \tilde{R}_c$). Further considering $R_v \approx R_t$, then the radius/thickness ratio of a budded organoid can be approximated as

$$\frac{R_c}{R_v} \approx w_c v^{-1/3}, \; \frac{h_c}{h_v} \approx \varphi(1-\varphi)^{-1} w_c^{-2} v^{2/3}, \quad (26)$$

where $w_c = \tilde{R}_c/\tilde{R}_0 = \varphi^{1/2}\alpha^{-1/3}\left(1+\varphi^{-1/2}\gamma_c\right)^{1/3}$. Eq. (26) indicates that the thickness (or radius) ratio of a budded organoid depends only on the crypt size $\varphi$ and $u = \alpha\left(1+\varphi^{-1/2}\gamma_c\right)^{-1}$, a parameter coupling the in-plane contraction and spontaneous curvature of crypt. As verified in Fig. S4B, the mechanical modulation of the thickness (or radius) ratio can be depicted by a single parameter $u$. Besides, Eq. (26) indicates a simple scaling law between organoid morphometrics and lumen volume for budded organoids: $R_c/R_v \sim v^{-1/3}$, $h_c/h_v \sim v^{2/3}$, which again shows excellent agreement with numerical solutions to the full model (Fig. S4B).

Strikingly, this predicts a key difference between the inflation of bulged vs budded organoids. In the former, the radius ratio is an increasing function of lumen volume (leading to near-spherical shapes upon inflation), while in the latter, the radius ratio always decreases with lumen volume (as the crypt never opens up, and the bulk of the deformation is born by the villus region). As discussed in the main text, we challenged this prediction via two different types of inflation experiments, and found good qualitative and quantitative agreement (Fig. 3B-C, Fig. S8A-B), see also Section 4 for details on the fitting strategy used.

Although the above derivations are based on Eq. (21), which can only describe organoids with small crypts, Eq. (26) actually holds for budded organoids with varied crypt sizes. In the following, we will directly use Eq. (7), a generic formulation of free energy, to derive Eq. (26). For a budded organoid, both $\theta_{\mathrm{c}}$ and $\theta_{\mathrm{v}}$ are close to 0, which lead to $N_{\mathrm{c}} / N'_{\mathrm{c}} \approx 1$, $N_{\mathrm{v}} / N'_{\mathrm{v}} \approx 1$. Then, Eq. (7) reduces to

$$F \approx 4\pi\left(\Gamma_a+\Gamma_b\right)_{\mathrm{c}} R_{\mathrm{c}}^2\left[1+2\left(\tilde{R}_{\mathrm{c}} / R_{\mathrm{c}}\right)^3\right]+4\pi\left(\Gamma_a+\Gamma_b\right)_{\mathrm{v}} R_{\mathrm{v}}^2\left[1+2\left(\tilde{R}_{\mathrm{v}} / R_{\mathrm{v}}\right)^3\right], \quad (27)$$

which is a function of two radii $R_{\mathrm{c}}$ and $R_{\mathrm{v}}$. These two radii should also satisfy the volumetric constraint, which is simplified as $R_{\mathrm{t}}^3 = R_{\mathrm{c}}^3 + R_{\mathrm{v}}^3$ in the budded case. Hence, the radii can be determined by constructing an auxiliary function that contains both free energy (27) and the volumetric constraint. For the normalized radii $\bar{R}_{\mathrm{c}} = R_{\mathrm{c}} / \tilde{R}_{\mathrm{c}}$ and $\bar{R}_{\mathrm{v}} = R_{\mathrm{v}} / \tilde{R}_{\mathrm{v}}$, the auxiliary function can be written as $y = F \big/ \left[\pi\left(\Gamma_a+\Gamma_b\right)_0 \tilde{R}_0^2\right] + L\left[w^3 \bar{R}_1^3 + \bar{R}_2^3 - \bar{R}_{\mathrm{t}}^3 \big/ \tilde{R}_2^3\right]$, where $L$ is a Lagrange multiplier , $w = \tilde{R}_{\mathrm{c}} / \tilde{R}_{\mathrm{v}}$. That is

$$y = \alpha w^2 \bar{R}_{\mathrm{c}}^2\left(1+2\bar{R}_{\mathrm{c}}^{-3}\right)+\bar{R}_{\mathrm{v}}^2\left(1+2\bar{R}_{2\mathrm{v}}^{-3}\right)+L\left[w^3 \bar{R}_{\mathrm{c}}^3+\bar{R}_{\mathrm{v}}^3-\bar{R}_{\mathrm{t}}^3 \big/ \tilde{R}_{\mathrm{v}}^3\right]. \quad (28)$$

Calculating $\partial y / \partial \bar{R}_{\mathrm{c}} = 0$ and $\partial y / \partial \bar{R}_{\mathrm{v}} = 0$ lead to $1-\bar{R}_{\mathrm{c}}^{-3} = w\alpha^{-1}\left(\bar{R}_{\mathrm{c}} / \bar{R}_{\mathrm{v}}\right)\left(1-\bar{R}_{\mathrm{v}}^{-3}\right)$, which will further result in Eq. (26) by using $\bar{R}_{\mathrm{c}} \approx 1$ and $\bar{R}_{\mathrm{v}} \approx v^{1/3} \tilde{R}_0 / \tilde{R}_{\mathrm{v}}$.

#### 2.2. Infinite volume expansion

As discussed above and in the main text, a key experimental finding is that budded organoids tend to stay closed upon volume expansion, while bulged organoids do not. To further explore the difference between the two morphologies, we examine the limit of infinite organoid inflation (i.e. $\beta \to \infty$), for which the boundary between these two morphologies in phase-space can be derived analytically.

We compare the free energies of organoids in partially open vs fully closed crypts. Since the partially open and fully closed crypt morphologies respectively belong to bulged and budded organoids discussed above, we can approximate their free energies by following the analysis in Subsection 2.1, and considering $\beta \to \infty$. Then, we have $\hat{F}_{\mathrm{po}} \approx 4\beta^2 + 12\varphi\left(\alpha-1\right)^{1/3}$ for a partially open case, and $\hat{F}_{\mathrm{fc}} \approx 4\beta^2 + 12\varphi\left(1+\varphi^{-1/2}\gamma_{\mathrm{c}}\right)^{2/3}\alpha^{1/3}$ for a fully closed shape. The crypt in a budded organoid will stay closed when $\hat{F}_{\mathrm{fc}} < \hat{F}_{\mathrm{po}}$, which holds for $\left(1+\varphi^{-1/2}\gamma_{\mathrm{c}}\right)^2 < 1-\alpha^{-1}$, which specifies a critical value of crypt apical tension distinguishing the two configurations.

A phase diagram of crypt morphologies under infinite lumen expansion are shown in Fig. S4C. The effects of in-plane contraction $\alpha$ and spontaneous curvature $\gamma_{\mathrm{c}}$ are examined in a representative parameter-regime: an initially large lumen (or thin monolayer) ( $\tilde{\kappa}_0 = 10$ ) and a large crypt region ( $\varphi = 0.2$ ). From the phase diagram, there also exists the third crypt shape: fully closed with vanishing apical surface (i.e. $R_{ac} = 0$ ). For a fully closed crypt in a budded organoid to get $R_{ac} = 0$, it needs to satisfy $\varphi^{1/2} + \gamma_{\mathrm{c}} = \left(2\tilde{\kappa}_0\right)^{-1}\alpha$.

##### 2.3. Dependence on cell volumes

In aforementioned derivations, cell volumes were set to be constant and identical in crypt and villus regions, i.e. $v_{\mathrm{ec}} = 1$ and $v_{\mathrm{ev}} = 1$. However, this is typically not the case, as discussed in the main text: Intestinal organoids display increases in cell volume as the lumen volume decreases during morphogenesis (Fig. 4D). To consider effects of cell volumes on specific development stages listed in Subsection 2.1, we incorporate the possibility for varying and different cell volumes to the simplified free energy (21), which is then modified as:

$$\begin{aligned}\hat{F} \approx\ & 4\beta^2 + 8(1-\varphi)^{3/2} v_{\mathrm{ev}}\beta^{-1} + \alpha s_{\mathrm{c}}\beta_{\mathrm{c}}^2 + \left(16\varphi^{3/2}s_{\mathrm{c}}^{-1/2} + 8\varphi\gamma_{\mathrm{c}}\right)v_{\mathrm{ec}}\beta_{\mathrm{c}}^{-1} \\ & + \left[(1-\varphi)^{3/2} v_{\mathrm{ev}}\beta^{-3} - 1\right]\left(\beta_{\mathrm{c}}^2 \sin^2\theta_{\mathrm{c}} + \frac{8}{3}p_{\mathrm{c}}\beta_{\mathrm{c}}^3\beta^{-1}\right)\end{aligned}, \qquad (29)$$

and follow the similar analysis in Subsections 2.1. In particular, we show that the generalized analytic expressions for the radius (or thickness) ratios, including Eqs. (23) and (26), become:

$$\begin{aligned}&\text{Bulged}: && \frac{R_{\mathrm{v}}}{R_{\mathrm{c}}} \approx 1 - \varphi^{-1}\gamma_{\mathrm{c}}v_{\mathrm{ec}}^{-1/3}\frac{\left[(\alpha-1)v + v_{\mathrm{ev}}\right]^{4/3}}{v - v_{\mathrm{ev}}}, \quad \frac{h_{\mathrm{c}}}{h_{\mathrm{v}}} \approx v_{\mathrm{ec}}^{1/3}v_{\mathrm{ev}}^{-1}\left[(\alpha-1)v + v_{\mathrm{ev}}\right]^{2/3} \\ &\text{Budded}: && \frac{R_{\mathrm{c}}}{R_{\mathrm{v}}} \approx \varphi^{1/2}v_{\mathrm{ec}}^{1/3}u^{-1/3}v^{-1/3}, \quad \frac{h_{\mathrm{c}}}{h_{\mathrm{v}}} \approx (1-\varphi)^{-1}v_{\mathrm{ec}}^{1/3}v_{\mathrm{ev}}^{-1}u^{2/3}v^{2/3}\end{aligned} .(30)$$

In view of the fact that cell swelling typically happens during the later development phases (Fig. 1, 2 and Fig. S9B), which correspond to the budding stage, we will discuss the dependence of cell volumes $v_{\mathrm{ec}}$ and $v_{\mathrm{ev}}$ on thickness (or radius) ratio only for budded organoids. For a budded organoid, scaling laws $R_{\mathrm{c}} / R_{\mathrm{v}} \sim v_{\mathrm{ec}}^{1/3}$ and $h_{\mathrm{c}} / h_{\mathrm{v}} \sim v_{\mathrm{ec}}^{1/3}v_{\mathrm{ev}}^{-1}$ are suggested by Eq. (30) and verified by numerical results in Fig. S4D. It can be seen from the scaling laws that, cell swelling in crypt always results in an increased radius (or thickness) ratio, while cell swelling in villus decreases the thickness ratio.

##### 2.4. Summary of analytic results

These analytic results provide insights into the physical mechanisms of crypt morphogenesis. As aforementioned, modulated by cell tensions, an epithelial sheet can engender two types of active deformations: in-plane contraction and spontaneous bending, which are respectively described by in-plane contraction ratio $\alpha$ and spontaneous curvature $\gamma_c$. However, in-plane contraction and bending can both vary at the same time (for instance if only the apical tension in crypt increases, all other parameters being kept constant) which implies the two mechanical variables $\alpha$ and $\gamma_c$ are combined to affect the geometric quantities of organoid epithelium, such as thickness (and radius) ratios. The analytic results in Subsection 2.1 indicates that the initial bulging morphology depends on $\alpha$ for the thickness ratio, $\varphi^{-1}\gamma_c$ for the radius ratio, while the budding configuration is only controlled by $u=\alpha\left(1+\varphi^{-1/2}\gamma_c\right)^{-1}$.

Here, we restricted ourselves to a two-region morphology (one crypt and one villus), although highly similar results are expected when considering more than one crypt region. Although in principle, budded shapes can arise even in the case of one-region organoids as explored by Rozman et al. (2019), who consider all cells of an organoid have equal properties, and budded shape can occur for remarkable apico-basal tension difference) (*22*), we note that this unlikely to occur in intestinal organoids, as i) we experimentally observed strong region differences in both actomyosin patterns and apico-basal tensions (assessed both via laser ablation in Fig. 2D and micropipette aspiration in Fig. 2E), and ii) one-region organoids are predicted to become spherical when inflated above a critical size, which is not what we observed in our inflation experiments (Fig. 3B-C, Fig. S8A-B).

In view of the fact that shape transformation from budded to open seldom happens even though the lumen volume increases dramatically by ~5 times (Fig. 3C'), the diagrams of crypt morphology with infinite volume (Fig. S4C) can be used to determine bounds for the parameters $\gamma_c$ and $\alpha$. We thus use these analyses and analytical criteria to guide the fitting of experimental data (both during normal organoid morphogenesis and upon organoid inflation).

#### 3. Morphogenesis with enhanced apical constriction and water uptake

To evaluate the influence of specific parameters on organoid morphologies, parameters in crypt mechanics (e.g. in-plane contraction $\alpha$ and spontaneous curvature $\gamma_c$) and volumes (e.g. organoid volume $v$ and volume of a villus cell $v_{ev}$) are usually analyzed separately in previous sections. Here, we focus on specific biophysical mechanisms uncovered by experiments, showing that these parameters may be coupled together to modulate organoid morphologies.

Firstly, experiments indicate that enhanced apical constriction of crypt is the leading mechanism in organoid morphogenesis. As assumed in Section 1, the initial spherical organoid has the same tension $\Gamma_0$ on both apical and basal surfaces, and the lateral tensions in both regions are $\Gamma_l$. With the morphological evolution of organoids, the accumulation of actomyosin on crypt apical surface leads to an increase in crypt apical tension, that is $\Gamma_{ac} = m\Gamma_0$ with $m$ the normalized crypt apical tension satisfying $m \geq 1$, while the other tensions are assumed to be constant, i.e. $\Gamma_{bc} = \Gamma_{av} = \Gamma_{bv} = \Gamma_0$. Considering that the size of the initial spherical organoid are regulated by two tensions $\Gamma_0$ and $\Gamma_l$, one can easily find the relation between the shape factor $\tilde{\kappa}_0$ and these two tensions, i.e. $\tilde{\kappa}_0 = \frac{\Gamma_l}{2\Gamma_0}\sqrt{\frac{N_t}{4\pi}}$. Then, one can rewrite in-plane contraction $\alpha$ and spontaneous curvature of crypt $\gamma_c$ as

$$\alpha = \frac{1+m}{2}, \gamma_c = \frac{1-m}{4\tilde{\kappa}_0}. \tag{31}$$

Eq. (31) shows that crypt apical tension can simultaneously modulate in-plane contraction $\alpha$ and spontaneous curvature $\gamma_c$, and that shape factor $\tilde{\kappa}_0$ is also important for the resulting shape. In this study, actomyosin accumulation is considered to be the sole mechanism that modulates cellular tensions, although other regulatory mechanisms, such as stretch-induced cortex dilation (*62*), are reported to be important for epithelia under deformation.

Secondly, experiments verify that villus cells up-regulate apical ion pumps that lead to the swelling of villus cells and shrinkage of the lumen (Fig. 4). This water uptake of villus cells will modulate two parameters in our model: the organoid volume $v$, which is the sum of the lumen volume and half the epithelial volume in villus, and the volume of villus cell $v_{ev}$. For simplicity, we assume the organoid volume is only modulated by water uptake. Then, during the water uptake of villus cells, the organoid volume is related to volume of a single villus cell $v_{\mathrm{ev}}$ as

$$v = 1 - \frac{3(1-\varphi)}{2\tilde{\kappa}_0}\left(v_{\mathrm{ev}} - 1\right). \tag{32}$$

Obviously, the water uptake will lead to a decrease in volume $v$. Before specific discussion on the influence of water uptake by villus cells on orgnaoid morphogenesis, we reassess the efficiency of water uptake by different cell types, although the influence of cell swelling has been evaluated in Subsection 1.4.1. Consider the relative reduction of lumen volume $\Delta v_{\mathrm{lu}}$ is compensated by i) volume increase in all cells, which lead to $v_{\mathrm{ec}} = v_{\mathrm{ev}} = \tilde{\kappa}_0 \Delta v_{\mathrm{lu}}/3 + 1$, ii) volume increase in crypt cells only, which yields $v_{\mathrm{ec}} = \tilde{\kappa}_0 \Delta v_{\mathrm{lu}}/(3\varphi) + 1$, and iii) volume increase in villus cells only, which results in $v_{\mathrm{ev}} = \tilde{\kappa}_0 \Delta v_{\mathrm{lu}}/[3(1-\varphi)] + 1$. For the water uptake by cells, including all the three cases above, the overall volume is $v \approx 1 - \Delta v_{\mathrm{lu}}/2$. Besides, we also consider the case that the reduction of luminal fluid is due to the leakage of epithelium, which corresponds to $v_{\mathrm{ec}} = v_{\mathrm{ev}} = 1$, and $v = 1 - \Delta v_{\mathrm{lu}}$. As shown in Fig. 4F, water uptake by villus cells is the most efficient mechanism for organoid budding.

We examine the influence of crypt apical constriction and water uptake of villus cells on organoid morphogenesis in Fig. S7. The water uptake is evaluated by the normalized volume of a villus cell $v_{\mathrm{ev}}$, and causes variations in the lumen volume, as shown in Eq. (32). As expected, both the enhanced apical constriction in crypt and water uptake of villus cells can lead to budding (Fig. S7A), and the critical apical tension and degree of water uptake are affected by the region size $\varphi$ and shape factor $\tilde{\kappa}_0$ (Fig. S7B). It is hard to close a large crypt by apical constriction alone, since the in-plane contraction of the epithelium will lead to the elevation of luminal fluid pressure, which further hinders the bending of the crypt, and a larger contractile region (i.e. the crypt) will lead to a higher fluid pressure. As shown in Eq. (31), spontaneous curvature is inversely related to the shape factor $\tilde{\kappa}_0$, thus strong apical constriction is needed for the budding of an organoid with a thin epithelium or a big lumen (i.e. large shape factor in Fig. S7B). Besides, as observed in experiments, an organoid usually undergoes enhanced apical constriction in the bulging stage, which is followed by the water uptake of villus cells. Setting the normalized crypt apical tension to $m=2$ ,we also examine the degree of water uptake that resulting in the closure of two regions in Fig. S7B.

The morphometric parameters, i.e. thickness ratio $h_{\mathrm{c}} / h_{\mathrm{v}}$ and radius ratio $R_{\mathrm{c}} / R_{\mathrm{v}}$, also evolve with apical constriction and water uptake. An organoid with enhancing crypt apical constriction may undergo three phases: Bulging, budding, and budding with vanishing crypt apical surface (i.e. $R_{a\mathrm{c}}=0$). In the first two phases, enhanced apical constriction in crypt leads to an increase in the thickness ratio and a decrease in the radius ratio (Fig. S7C-D), and the transformation from the bulged shape to the budded one results in negligible variations in the trends of thickness ratios and notable changes in those of radius ratios. With continued enhancement of apical constriction, the apical surface of a closed crypt will contract towards a point, then the crypt will stop thickening and the thickness (or radius) ratio goes into a plateau, as shown in Fig. S7C. Morphometric parameters in these three phases are also affected by shape factor $\tilde{\kappa}_0$ and region size $\varphi$. As aforementioned, spontaneous curvature $\gamma_{\mathrm{c}}$ results in an increased thickness and decreased radius of the crypt. Considering $\gamma_{\mathrm{c}}$ is inversely proportional to shape factor $\tilde{\kappa}_0$ (Eq. (31)) , one can find that a larger $\tilde{\kappa}_0$ will leads to a smaller thickness ratio $h_{\mathrm{c}} / h_{\mathrm{v}}$ and larger radius ratio $R_{\mathrm{c}} / R_{\mathrm{v}}$ in the bulging and budding phases, as verified in Fig. S7C. While crypt size $\varphi$ has negligible influence on the thickness (or radius) ratio in these two phases (Fig. S7D). However, both shape factor $\tilde{\kappa}_0$ and crypt size $\varphi$ are crucial on the third phase. Since the apical surface of an organoid with a thin epithelium/large lumen is hard to contract into a point, a large shape factor will delay the happening of the third phase (Fig. S7C). And a large crypt in the budding phase will become a closed sphere with large radius $R_{\mathrm{c}}$, which makes it hard to get $R_{a\mathrm{c}}=R_{\mathrm{c}}-h_{\mathrm{c}} / 2=0$ and enter into the third phase (Fig. S7D). After enhanced apical constriction in crypt, water uptake of villus cells will keep promoting the morphogenesis. As expected, water uptake of villus cells will decrease the thickness ratio and promote the closure of two region (Fig. S7E). Morphometric parameters show distinct trends in bulging and budding phases. With the water uptake, the thickness ratio decreases more sharply in a budded organoid than in a bulged one, while the radius ratio only shows notable changes in the bulging phase.

We further explore the influence of crypt apical constriction on the evolution of thickness ratio $h_{\mathrm{c}} / h_{\mathrm{v}}$ and radius ratio $R_{\mathrm{c}} / R_{\mathrm{v}}$ during organoid expansion. As expected and shown in Fig.

S7F, the thickness ratio always increases with volume expansion for an organoid with enhanced crypt apical constriction, which prevents the crypt from inflating with organoid expansion. As already found in Fig. S2 and analyzed in Subsection 2.1, for a bulged organoid with weak crypt apical constriction, the radius ratio increases with volume expansion, while the radius ratio of a budded organoid decreases with volume expansion (Fig. S7F). Besides, the transformation from a budded shape to a bulged one can also happen, and will also affect the thickness (or radius) ratio. We also discuss morphologies of crypts with enhanced apical constriction under infinite volume expansion in Fig. S7G. Setting crypt size $\varphi = 0.2$, the crypt morphologies are modulated by two parameters: normalized crypt apical tension $m$ and shape factor $\tilde{\kappa}_0$. The phase diagram indicates that the three morphologies discussed in Fig. S4C (partially open, fully closed, and fully closed with vanishing apical surface) still exist for crypts with enhanced apical constriction. Inserting Eq. (31) into the critical condition in Subsection 2.2, one obtains that the crypt will never open up if the normalized crypt apical tension $m$ satisfies $\left[1+(1-m)/\left(4\varphi^{1/2}\tilde{\kappa}_0\right)\right]^2 + 2/(1+m) < 1$ (whose lower bound is named as $m_{\text{crit}}$ afterwards), no matter the lumen volume. However, when $m$ is larger than $2\varphi^{1/2}\tilde{\kappa}_0$, the apical surface will contract into a point, resulting in $R_{ac}=0$. The influence of cell swelling on thickness (or radius) ratio have been discussed in Subsection 2.3, and both numerical and analytic results indicate that the crypt morphologies under infinite organoid expansion are irrelevant to cell volume $v_{\text{ev}}$.

#### 4. Organoid morphometric measurements and fitting strategy

To validate the theory and extract mechanical parameters, we measured the thickness (and radius) ratios of crypt and villus during normal organoid morphogenesis (Fig. 2C) and inflation experiments, when the lumen volume is increased by PGE treatment (Fig. 3B-C) or micropipette injection (Fig. S8A-B). In the measurements, a dimensionless volume $\bar{v}$, which is the current volume of a sample normalized by its originating volume, is used to characterize the organoid inflation. Let $v_0$ be the initial volume, $\tilde{v}$ be the volume in free mechanical state, which can be estimated as $\tilde{v} \approx (1-\varphi)^{3/2} v_{c2}$ by using $\partial\hat{F}/\partial\beta = 0$ in Eq. (29), then $\bar{v}$ is related to the volume $v$ employed in the model as $\bar{v} = v/\left(v_0\tilde{v}\right)$. Considering the crypt apical constriction as the main mechanical cue of organoid morphogenesis in bulding phase, then the crypt mechanical parameters can be described by Eq. (31). Moreover, luminal volume decrease and swelling of villus cells occurs in the budding phase (Fig. 4). In view of these, one can find that the evolutions of thickness (and radius) ratios depend on four parameters: $m$, $\tilde{\kappa}_0$, $\varphi$, and $v_0$, for bulged samples, and one more parameter $v_{\mathrm{ev}}$ for budded ones. We can directly measure some of them, and determine the remaining parameters by fitting experimental data with analytic formulation or numerical results.

##### 4.1. Independently-measured geometric parameters

Firstly, to estimate the shape factor of organoids, we can measure the shape factor of villus $\kappa_{\mathrm{v}} = 4\pi R_{\mathrm{v}}^3 \big/ \left(N_{\mathrm{v}} V_{\mathrm{ev}}\right) \approx R_{\mathrm{v}} / h_{\mathrm{v}}$, which is linearly dependent on $\bar{v}$ as $\kappa_{\mathrm{v}} \approx \kappa_{\mathrm{v}0}\bar{v}$, with the initial shape factor of villus $\kappa_{\mathrm{v}0}$ related to $\tilde{\kappa}_0$ as $\kappa_{\mathrm{v}0} \approx \tilde{\kappa}_0 v_0$ for bulged organoids and $\kappa_{\mathrm{v}0} \approx \left(1-\varphi\right)^{1/2} \tilde{\kappa}_0 v_0$ for budded ones. Secondly, we also directly measured the crypt size $\varphi$ of bulged organoids as $\varphi \approx h_{\mathrm{c}} l_{\mathrm{c}}^2 \big/ \left(4 h_{\mathrm{v}} R_{\mathrm{v}}^2\right)$, where the arclengh of crypt section is denoted $l_{\mathrm{c}}$, to further constrain the system.

##### 4.2. Parameters extracted via direct fitting.

Analytic results in Section 2 provide guidance on the fitting of morphometric data. Replacing volume $v$ by the new normalized volume $\bar{v}$ in Eq. (30) leads to

$$\begin{aligned} &\text{Bulging:} \quad \frac{R_{\mathrm{v}}}{R_{\mathrm{c}}} \approx 1 - \varphi^{-1}\gamma_{\mathrm{c}}\bar{v}_{\mathrm{e}}^{-1/3}\frac{\left[\left(\alpha-1\right)v_0\bar{v}+1\right]^{4/3}}{v_0\bar{v}-1}, \quad \frac{h_{\mathrm{c}}}{h_{\mathrm{v}}} \approx \bar{v}_{\mathrm{e}}^{1/3}\left[\left(\alpha-1\right)v_0\bar{v}+1\right]^{2/3} \\ &\text{Budding:} \quad \frac{R_{\mathrm{c}}}{R_{\mathrm{v}}} \approx \varphi^{1/2}\left(1-\varphi\right)^{-1/2}\bar{v}_{\mathrm{e}}^{1/3}\left(uv_0\right)^{-1/3}\bar{v}^{-1/3}, \quad \frac{h_{\mathrm{c}}}{h_{\mathrm{v}}} \approx \bar{v}_{\mathrm{e}}^{1/3}\left(uv_0\right)^{2/3}\bar{v}^{2/3} \end{aligned} \quad ,(33)$$

where $\bar{v}_{\mathrm{e}} = v_{\mathrm{ec}} / v_{\mathrm{ev}}$ is the volume ratio of a crypt cell to a villus cell. In this new formulation, thickness (or radius) ratio is only related to cell volumes by $\bar{v}_{\mathrm{e}}^{1/3}$. As aforementioned, cell swelling is insignificant in the bulging phase, but the swelling of villus cells becomes important in the budding phase. Therefore, we have $\bar{v}_{\mathrm{e}} \approx 1$ for a bulged organoid, and $\bar{v}_{\mathrm{e}} = v_{\mathrm{e}2}^{-1} < 1$ for a budded one. During the bulging of organoids, the crypt mechanical parameters vary with time, while the lumen volume stays almost constant. In contrast, the inflation experiments provide a setting where lumen volumes change drastically while crypt mechanics can be considered constant. In view of these, based on the analytical expressions of Eq. (33), we further discuss specific relations between morphometric parameters, i.e. thickness (and radius) ratios, and bulging time or lumen volume in the following, and also derive the relation between thickness ratio and radius ratio. Using these analytical relations, we can fit and rescale the experimental data.

##### 4.2.1. Dynamics of organoid bulging

First, we consider the bulging dynamics of organoids. Experiments show that the volume stays constant in this process, that is $\bar{v} \approx 1$. According to Eq. (33), the morphometric parameters $R_c / R_v$ and $h_c / h_v$ are then linked to each other via a simple scaling relation:

$$\frac{R_v}{R_c} \approx 1 + \mathrm{pm1} \cdot (h_c/h_v)^2 \left[ (h_c/h_v)^{3/2} - 1 \right], \tag{34}$$

where pm1= $\left[2\varphi\kappa_{v0}(v_0 - 1)\right]^{-1}$ is a single fitting parameter. Importantly, this expression is independent on the dynamics of how crypt apical tension varies in time, providing a simple and robust model prediction. Extracting pm1 from the data, and using the geometric parameters independently measured i.e. shape factor $\kappa_{v0}$: 3.9±2.1 (mean ± SD) (respectively 3.33, 2.05, and 6.21 for three samples), and crypt size $\varphi$: 0.19±0.1 (mean ± SD) (respectivley 0.2, 0.09, and 0.29), then allowed us not only, to estimate $v_0$ (see below), but also rescale the morphometric parameters of every sample to verify that, even for organoids with distinct initial shape factors and crypt sizes, their morphometric parameters were well-fitted on the master curve predicted in Eq. (34), as shown in Fig. 2C'''.

To further reproduce the morphological evolution of each ratio in time, we must then assume a specific dynamics for tension changes in time. For simplicity, we consider a linear increase of the normalized crypt apical tension $m$ with time $t$, that is $m = m_0 + m't$, where $m_0$ and $m'$ are respectively the initial value and the slope. Then, the evolution of thickness ratio $h_c / h_v$ and radius ratio $R_c / R_v$ can be estimated as

$$\frac{h_c}{h_v} \approx (\mathrm{pm2} + \mathrm{pm3} \cdot t + 1)^{2/3}, \quad \frac{R_v}{R_c} \approx 1 + \mathrm{pm1} \cdot (\mathrm{pm2} + \mathrm{pm3} \cdot t)(\mathrm{pm2} + \mathrm{pm3} \cdot t + 1)^{4/3}, \tag{35}$$

where pm2=$(m_0 - 1)v_0 / 2$, pm3=$m'v_0 / 2$. We can get pm2 and pm3 simultaneously by fitting the experimental data of thickness ratios, and obtain pm1 by fitting the data of radius ratios (both of the evolutions were well-fitted by these scaling forms, see Fig. S3C-D). For the three samples we measured, we can get their mean value of initial volume $v_0$: 2.4 ± 0.5 (mean ± SD), and the initial crypt apical tension $m_0$:1.3 ± 0.2 (mean ± SD). We can also get the enhanced crypt apical tension *m* at the end of the bulging phase (prior to water uptake by villus cells): 1.8 ± 0.6 (mean ± SD). This value is interesting, as it remains significantly smaller (60%) of the critical value of *m* that leads to crypt budding (calculated for each individual sample taking into account crypt domain size and shape factor), and also argues that changes in lumen volume will play a key role on crypt morphogenesis. Furthermore, *m* is also much smaller than the critical value $m_{\text{crit}}$ that allows to remain budded upon infinite volume expansion (40%).

##### 4.2.2. Inflation of bulged organoids

Our analysis of the dynamics of bulging organoid suggests that their apical tension *m* is below the critical point of Fig. 3A, so that these organoids would be expected to open up upon inflation, a key prediction we now test. For the inflation of bulged samples, we can assume that tensions remain constant, and eliminate volume from the equation to derive again a relation between $R_c / R_v$ and $h_c / h_v$:

$$\frac{R_v}{R_c} \approx 1 + \frac{\left(h_c/h_v\right)^2}{\text{pg1}\cdot\left[\left(h_c/h_v\right)^{3/2} - \alpha\right]}, \tag{36}$$

where $\text{pg1} = -\varphi\gamma_c^{-1}\left(\alpha - 1\right)^{-1}$. In contrast to the relation of morphometric parameters in Eq. (34) for bulging evolution, the thickness ratio $h_c / h_v$ and the radius ratio $R_c / R_v$ show similar trends during organoid inflation. The dependence of morphometric parameters on volume $\bar{v}$ yields

$$\frac{h_c}{h_v} \approx \left(\text{pg2}\cdot\bar{v} + 1\right)^{2/3},\ \frac{R_v}{R_c} \approx 1 + \frac{\left(\text{pg2}\cdot\bar{v} + 1\right)^{4/3}}{\text{pg3}\cdot\left(v_0\bar{v} - 1\right)}, \tag{37}$$

where $\text{pg2} = \left(m - 1\right)v_0/2$, $\text{pg3} = 2\varphi\kappa_{v0}/\text{pg2}$. The experimental data again was in good agreement with these scaling forms (Fig. S8C-C'), so that by fitting the experimental data of thickness ratios, we can get pg2, which can be further used to estimate pg3. Then, the initial volume $v_0$ is employed as the only fitting parameter to fit the data of radius ratios. Further using the relations between parameters in Eq. (36) and those in Eq. (37): $\text{pg1} = v_0 \cdot \text{pg3}/\text{pg2}$, $\alpha = 1 + \text{pg2}/v_0$, we can find that the functional form of Eq. (36) predicts well the evolution of all six bulged inflation samples (from PGE or pipette), as shown in Fig. 3D. For the six samples we measured, we can get estimates of the initial volume $v_0$: 1.5 ± 0.3 (mean ± SD) (i.e. always larger than 1, consistent with initially swollen organoids as found in the fits of the bulging evolution), and the normalized crypt apical tension $m$: 1.3 ± 0.05 (mean ± SD), which is close to the intial tension $m_0$ estimated for the three bulging samples in Subsection 4.2.1, providing a good consistency check of the fitting approach and model.

##### 4.2.3. Inflation of budded organoids

Finally, as aforementioned, the morphometric parameters of budded samples obey a simple scaling law, and we can easily get the relation between $R_c / R_v$ and $h_c / h_v$:

$$\frac{R_c}{R_v} = \left(\text{pd1} \cdot h_c / h_v\right)^{-1/2}, \tag{38}$$

where $\text{pd1} = \varphi^{-1}(1-\varphi)v_{ev}$, and their relation with $\bar{v}$ can be recast as

$$\frac{h_c}{h_v} \approx \left(\text{pd2} \cdot \bar{v}\right)^{2/3}, \; \frac{R_c}{R_v} = \left(\text{pd3} \cdot \bar{v}\right)^{-1/3}, \tag{39}$$

where $\text{pd2} = v_{ev}^{-1/2} \cdot uv_0$, $\text{pd3} = \text{pd1}^{2/3} \cdot \text{pd2}$. These scaling relationships can in fact be derived from purely geometric considerations, under the assumption that near-spherical villi bear the deformation alone. To estimate the normalized crypt apical tension $m$, we can further introduce the shape factor of crypt $\kappa_c = 4\pi R_c^3 / (N_c V_{ec})$, which can be estimated either by directly using $\kappa_c \approx R_c / h_c$ or by using its relation with other fitting parameters, that is $\kappa_c = \kappa_{v0}\left(\text{pd2}^2 \cdot \text{pd3}\right)^{-1/3}$, then the normalized crypt apical tension is

$$m = \frac{4\varphi^{1/2}\tilde{\kappa}_0 + 2}{2\kappa_c + 1} - 1. \tag{40}$$

Although $\varphi^{1/2}\tilde{\kappa}_0$ is hard to get by direct measurement, we can use the analytic critial conditions of crypt morphologies under infinite volume expansion (discussed in Subsection 2.2 and Section 3) to estimate the lower bound of $m$ (i.e. $m_{crit}$). Now we have $\left(\frac{m+1}{m-1}\right)^{3/2} - \frac{m+1}{m-1} < \frac{1}{2\kappa_c}$ to forbid the crypt to open up with volume expansion and $\kappa_c > 0.5$ to guarantee that $R_{ac} > 0$ always holds. For the six samples we measured, we can get an estimation of $m_{crit}$: 3.6 ± 0.8 (mean ± SD). Alternatively assuming, based on the pattern of apical myosin intensity in bulged vs budded crypts (Fig. 2F'), that tensions *m* are roughly twice larger in budded organoids would also predict values close to the critical threshold for stable crypts upon inflation (see Subsection 4.2.2).

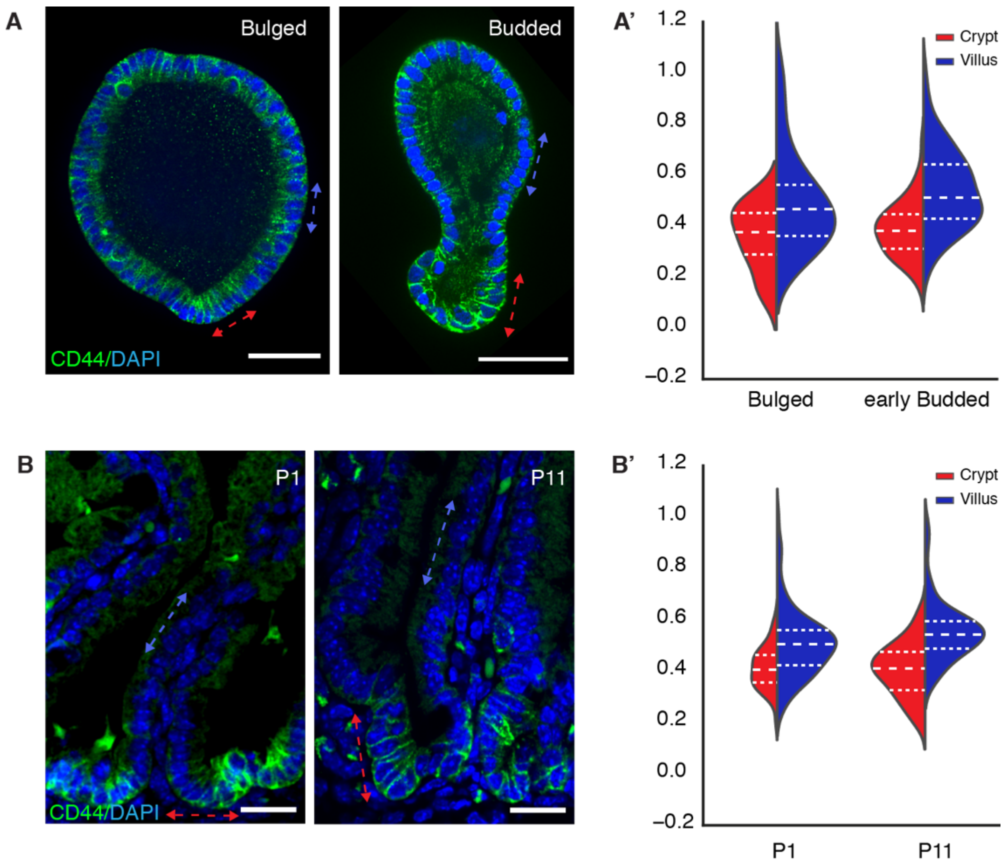


**Supplementary Figure 1. Tissue compaction along crypt-villus axis in the development of intestinal organoid and *in vivo* tissue. A**, Representative images of bulged and budded organoids. CD44 marks the regions with more stem and progenitor cell types (green), DAPI stains the cell nuclei (blue). Scale bars, 50 µm. **A'**. Plot of the normalized (to the size of nuclei diameters) distance between the nuclei of neighbour cells in crypt (red dashed double arrow head line) and villus (blue dashed double arrow head line) tissue as indicated in A. Two-tailed t-test for bulged crypt (n= 17 from 3 organoids) and bulged villus (n = 10 from 3 organoids), $p = 0.017$, for budded crypt (n= 53 from 8 organoids) and budded villus (n = 65 from 8 organoids), $P < 10^{-8}$. Violin plot lines denote quartile for each group. **B**. Representative images of immunostaining on the section of mouse intestine at the age of P1 and P11. Scale bars, 50 µm. **B'**. Plot for the normalized distance between the neighbour nuclei in crypt (red dashed double arrow head line) and villus (blue dashed double arrow head line) tissue as indicated in B. Two-tailed t-test for P1 crypt (n= 38 from 7 different crypt regions) and P1 villus (n = 87 from 7 villus regions), $p < 10^{-4}$, for P11 crypt (n= 54 from 6 different crypt regions) and P11 villus (n = 60 from 6 different villus regions), $P < 10^{-10}$. Violin plot lines denote quartile for each group.

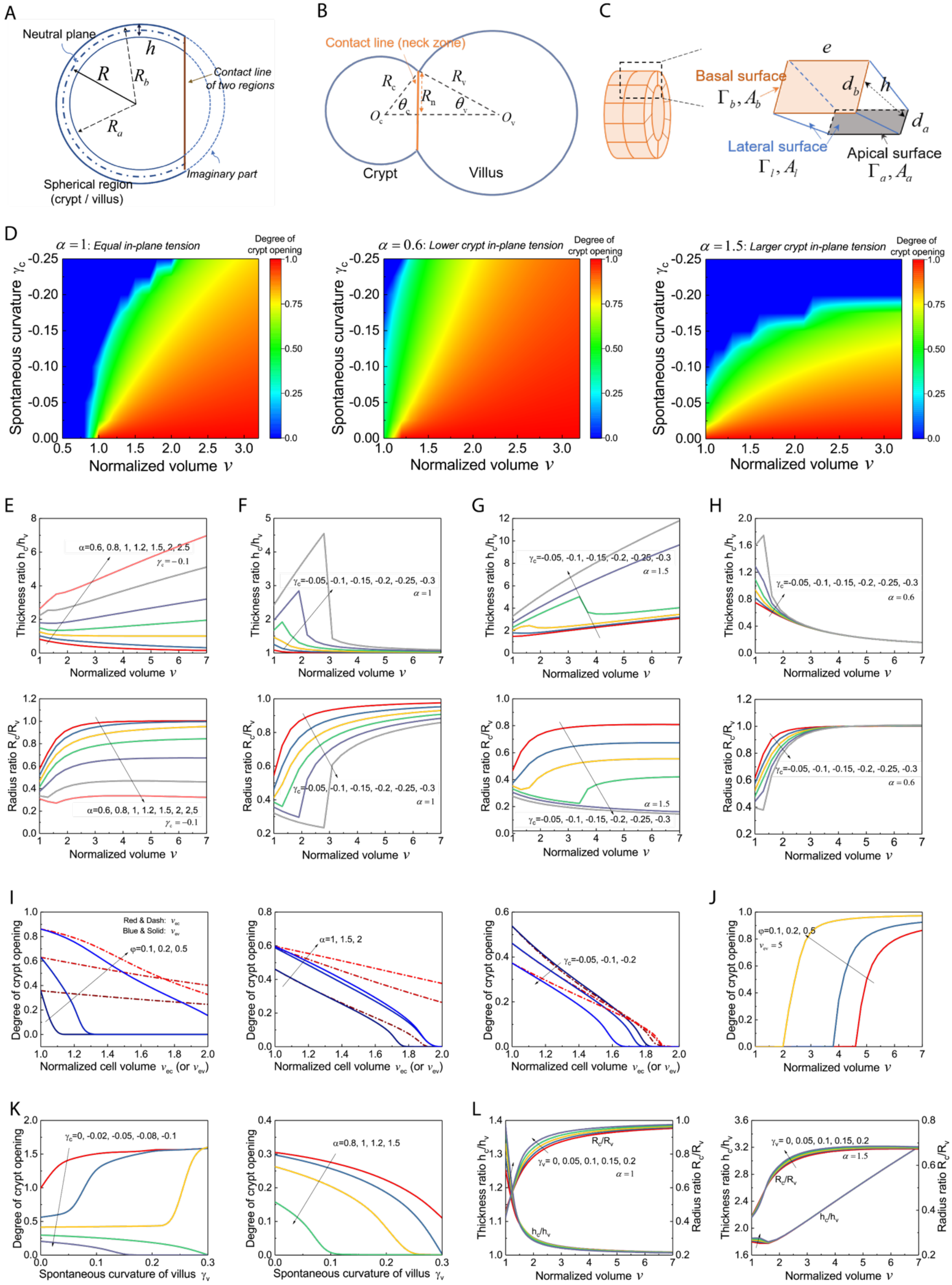
A
Neutral plane
h
R
$R_b$
$R_a$
Contact line of two regions
Imaginary part
Spherical region (crypt / villus)
B
Contact line (neck zone)
$R_c$
$R_v$
$R_n$
$\theta_c$
$\theta_v$
$O_c$
$O_v$
Crypt
Villus
C
e
Basal surface
$\Gamma_b, A_b$
$d_b$
h
$d_a$
Lateral surface
$\Gamma_l, A_l$
Apical surface
$\Gamma_a, A_a$
D
$\alpha = 1$: Equal in-plane tension
$\alpha = 0.6$: Lower crypt in-plane tension
$\alpha = 1.5$: Larger crypt in-plane tension
Degree of crypt opening
Spontaneous curvature $\gamma_c$
Normalized volume $v$
E
F
G
H
Thickness ratio $h_c/h_v$
Radius ratio $R_c/R_v$
Normalized volume $v$
α=0.6, 0.8, 1, 1.2, 1.5, 2, 2.5
$\gamma_c = -0.1$
$\gamma_c$=-0.05, -0.1, -0.15, -0.2, -0.25, -0.3
$\alpha = 1$
$\alpha = 1.5$
$\alpha = 0.6$
I
J
Degree of crypt opening
Red & Dash: $v_{ec}$
Blue & Solid: $v_{ev}$
φ=0.1, 0.2, 0.5
α=1, 1.5, 2
$\gamma_c$=-0.05, -0.1, -0.2
Normalized cell volume $v_{ec}$ (or $v_{ev}$)
$v_{ev} = 5$
Normalized volume $v$
K
L
$\gamma_c$=0, -0.02, -0.05, -0.08, -0.1
α=0.8, 1, 1.2, 1.5
Spontaneous curvature of villus $\gamma_v$
$\gamma_v$= 0, 0.05, 0.1, 0.15, 0.2
$R_c/R_v$
$h_c/h_v$
$\alpha = 1$
$\alpha = 1.5$
Thickness ratio $h_c/h_v$
Radius ratio $R_c/R_v$
Normalized volume $v$

**Supplementary Figure 2. Sensitivity analysis for how model parameters affect crypt morphology.** **A-C**. Schematic of the model and morphometric parameters used (see SI Text for details). **D**. Phase diagrams of crypt morphologies with varying volumes $v$ and spontaneous curvature of crypt $\gamma_c$ , for different values of in-plane contraction $\alpha$ (left to right: 1, 0.6, and 1.5). **E-H**. Evolution of thickness ratio $h_c/h_v$ and radius ratio $R_c/R_v$ during the inflation of an organoid (crypt size $\varphi = 0.2$, shape factor $\tilde{\kappa}_0 = 10$) : increasing $\alpha$ ($\gamma_c = 0.1$) (panel E) and $\gamma_c$ ($\alpha$ =1, 1.5, and 0.6) ( panels F-H). **I**. Influence of cell swelling on crypt morphology (degree of crypt opening) with varied crypt size $\varphi$ ($\alpha = 1.5, \gamma_c = -0.02$), $\alpha$ ($\gamma_c = -0.1, \varphi = 0.5$), and $\gamma_c$ ($\alpha = 1, \varphi = 0.5$). **J**. Morphological evolution during the inflation of an organoid ($\alpha = 1.5$, $\gamma_c = -0.02$) with swollen villus cells ($v_{ev} = 5$). **K**. Influence of spontaneous curvature of villus $\gamma_v$ on crypt morphology ($\varphi = 0.2$) with varied $\gamma_c$ ($\alpha = 1$) and $\alpha$ ($\gamma_c = -0.08$). **L**. Influence of $\gamma_v$ on the evolution of $h_c/h_v$ and $R_c/R_v$ during the inflation of an organoid with $\alpha$ is respectively 1 and 1.5 ($\gamma_c = -0.1, \varphi = 0.2$).

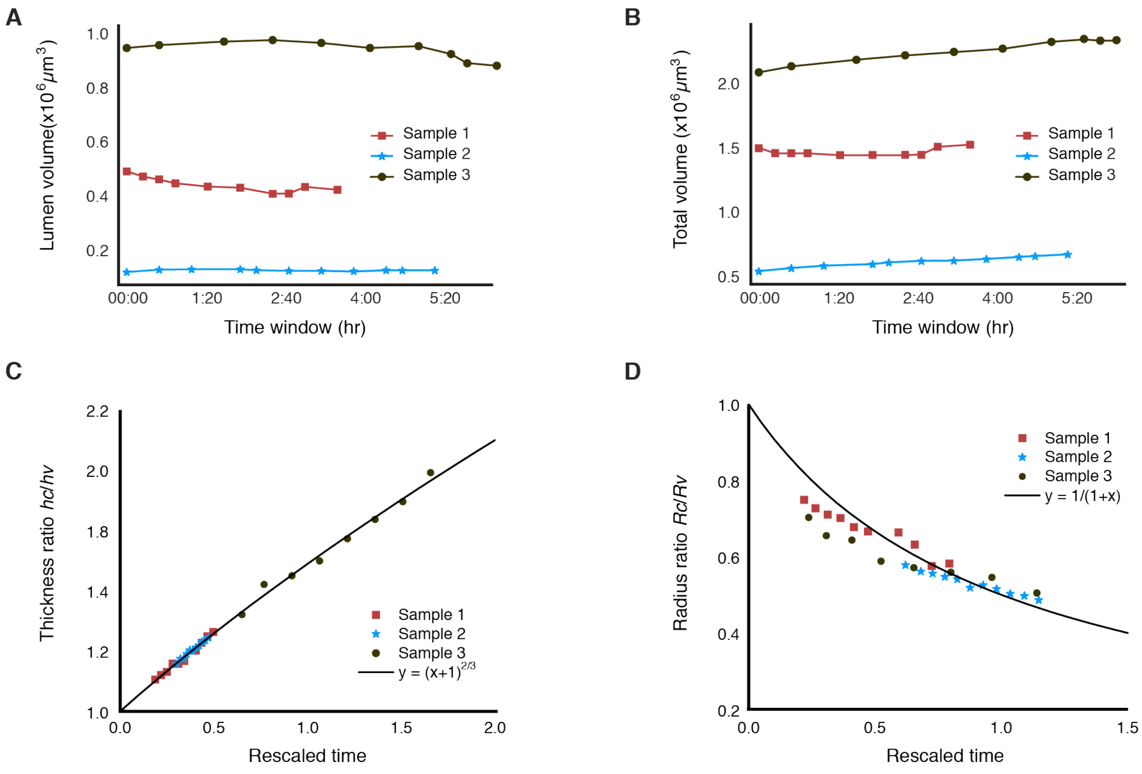


**Supplementary Figure 3. Evolution of the morphometric parameters during organoid bulging. A-B**. Plots of the dynamics of organoid lumen (A) and total (B) volume during crypt bulging of samples indicated in Fig. 2C. For each sample, 10 time windows were selected based on morphological change of organoid from Day3 cyst shape till bulging. **C-D**. All three samples of bulged organoids can be collapsed via $R_c/R_v$ vs time (C) or $h_c/h_v$ vs time (D).

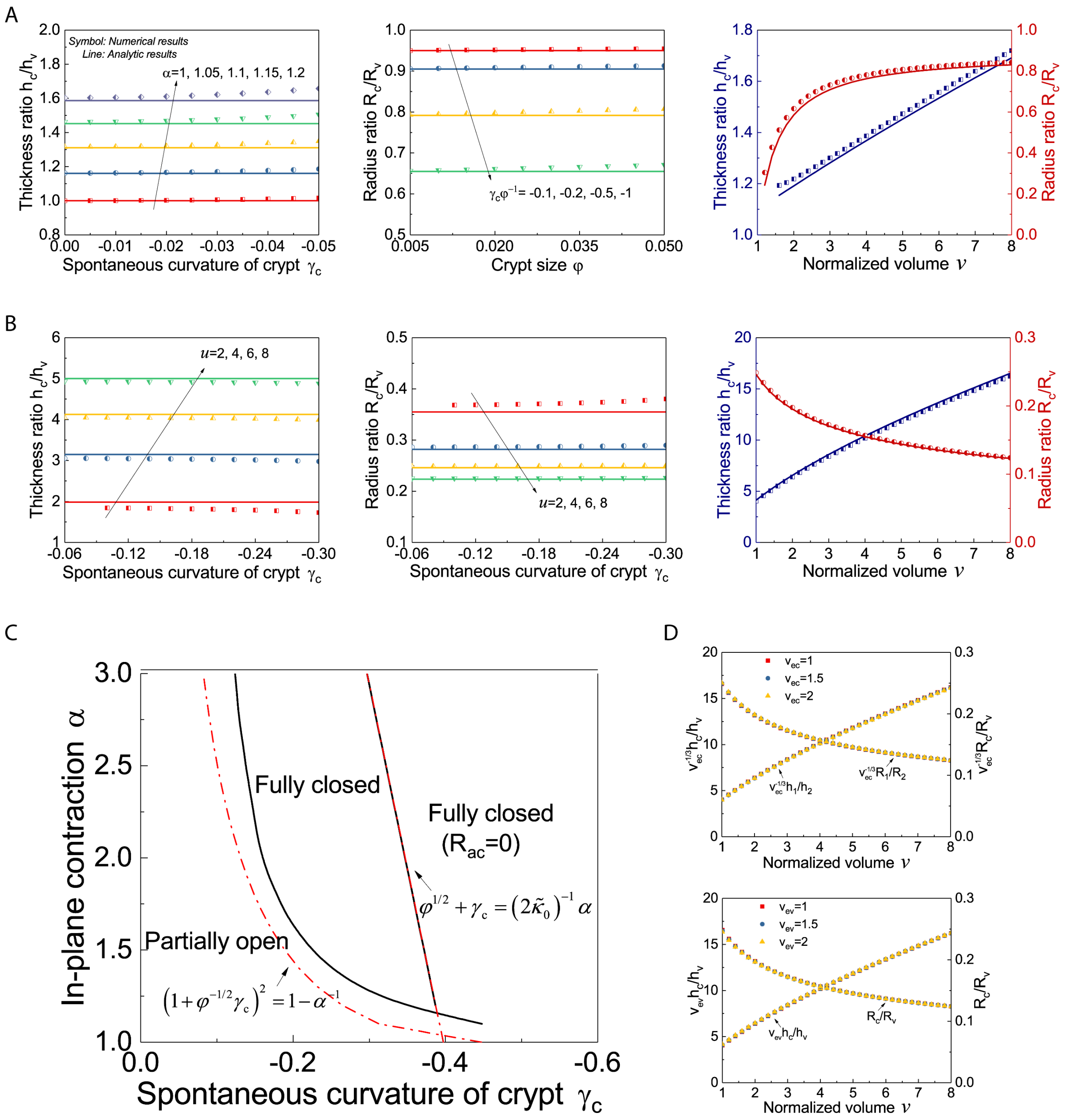


**Supplementary Figure 4. Comparison between numerical solutions of the full model and analytical scaling laws. A-B**. Comparison of numerical (symbol) and analytic (line) results to verify that, thickness ratio $h_c/h_v$ and radius ratio $R_c/R_v$ respectively depends on in-plane contraction $\alpha$ (with crypt size $\varphi = 0.05$) and coupled parameter $\varphi^{-1}\gamma_c$ (with $\alpha = 1.15$) for a bulged organoid (normalized volume $v = 5$) (panel A), both $h_c/h_v$ and $R_c/R_v$ depend on parameter $u$ for a budded organoid ($\varphi = 0.2$) (panel B), and analytic results also fit well with numerical results for the inflation of a bulged organoid ($\alpha = 1.15$, $\gamma_c = -0.025$, $\varphi = 0.05$) or a budded organoid ($u = 6$, $\varphi = 0.2$). **C**. Phase diagram of crypt morphologies of an organoid ($\varphi = 0.2$, $\tilde{\kappa}_0 = 10$) under infinite volume expansion ($v = 10^8$) (see SI Text for details), with varying spontaneous curvature $\gamma_c$ and in-plane contraction $\alpha$. **D**. Influence of normalized volume of a crypt cell $v_{ec}$ (top) and that of a villus cell $v_{ev}$ (bottom) on $h_c/h_v$ and $R_c/R_v$ of a budded organoid ($u = 6$, $\varphi = 0.2$).

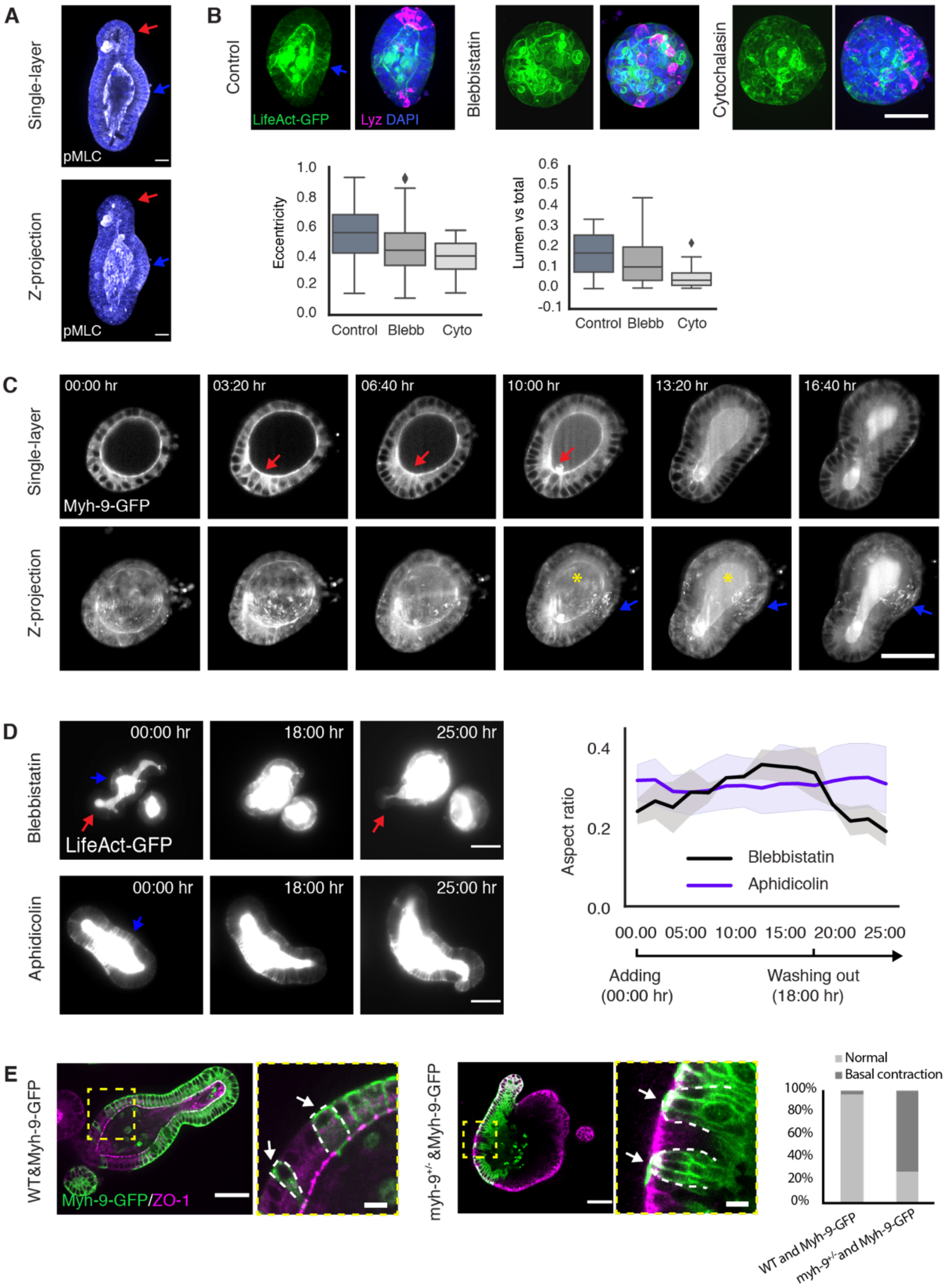


**Supplementary Figure 5**: **Actomyosin drives and maintains crypt budding. A.** Staining of phosphorylated Myosin light chain (pMLC, white) and DAPI staining for cell nuclei (blue) in budded organoid with maximum z-projection (upper panel) and single section (lower panel). Red arrows indicate crypt region, blue arrows indicate villus region. Scale bars, 20 µm. **B**. Actomyosin is required for crypt morphogenesis. Images, representative images of Day4 organoid expressing LifeAct-GFP (green) cultured in control condition treated with 10 µM DMSO or samples treated with 7.5 μM (-) Blebbistatin to inhibit Myosin activity or 1 μM Cytocalasin D to inhibit Actin polymerization. For each group of images, left, maximum z-projection of LifeAct-GFP; right, merged maximum z-projection images of LifeAct-GFP with Lyz staining for Paneth cells (magenta) and DAPI (blue). Blue arrow in control indicates the enrichment of LifeAct-GFP in villus region. Scale bars, 50 μm. Plots, Box plot for the quantification of organoid eccentricity and ratio of lumen volume *vs.* total volume (Lumen vs total). Two-tailed t-test for eccentricity to Control (n = 91 from three replications) with (-) Blebbistatin (n= 88 from three replications, $p < 10^{-5}$) and Cytocalasin D (n= 43 from three replications, $p < 10^{-5}$); and for lumen ratio with (-) Blebbistatin (p = 0.008) and Cytocalasin D ( $p < 10^{-10}$). Box plot elements show quartiles, and whiskers denote 1.5× the interquartile range. **C.** Representative light-sheet time-lapse images of organoid expression Myh-9-GFP (white) during crypt morphogenesis. Upper panel, images of middle sections. Lower panel, images of maximum z-projections. Red arrows indicate the enrichment of crypt apical Myh-9-GFP during bulging, blue arrows indicate the enrichment of villus basal Myh-9-GFP during budding. Myh-9-GFP signal at the basal domain in the center of villus region is not as bright as demonstrated in confocal images in Fig.2F (yellow asterisk), likely due to the limitation of light-sheet on z-axis resolution. Scale bar, 50 μm. **D**. Actomyosin is required for maintaining the crypt morphology. Images, light-sheet time-lapse recording of branched organoid treated with 7.5 μM Blebbistatin to inhibit myosin activity (upper panel, n = 2) or 0.6 μM Aphidicolin to block cell cycle (lower panel, n = 3) for 18 hours and washed out, cultured and recorded for another 7 hours. Organoids are visualized by LifeAct-GFP (white). Blue arrows indicate the enrichment of LifeAct-GFP in villus regions in budded organoids. Red arrows in blebbistatin treated organoid indicate the re-grow of crypt after Blebbistatin removal. Scale bars, 50 μm. Plot, corresponding plot for aspect ratio (width of the neck *v.s.* length of the organoid) of the organoids**.** Light-sheet recording of samples are with 10-mintues time interval, 150 time-points each. Data are collected from every 10 time windows (1:40 hrs). Solid lines represent the average values, shadow regions represent the standard deviations. **E.** Mosaic experiment indicates Myosin II is required for villus basal tension. Images, merged images for Myh-9-GFP (green) and ZO-1 staining (magenta) in organoid with wild-type and Myh-9-GFP cells (left group) or *myh-9*$^{+/-}$ and Myh-9-GFP cells (right group) in villus region. For each group of images, yellow dashed rectangles indicate the areas with two cell types, zoomed-in in right images. White arrows indicate representative Myh-9-GFP positive cells in the mosaic regions, with relatively relaxed basal domains (normal, as shown in wild-type and Myh-9-GFP mosaic) or with constricted basal domain (basal contraction, as shown in *myh-9*$^{+/-}$ and Myh-9-GFP mosaic). Scale bars, in left images of each groups, 50 μm, in right zoomed-in images, 10 μm. Plot, quantification on percentage of the basal constriction in Myh-9-GFP positive cells next to negative cells of WT (n = 70 from 16 organoids) and *myh-9*$^{+/-}$ (n = 79 from 21organoids), data are pulled from two independent experiments.

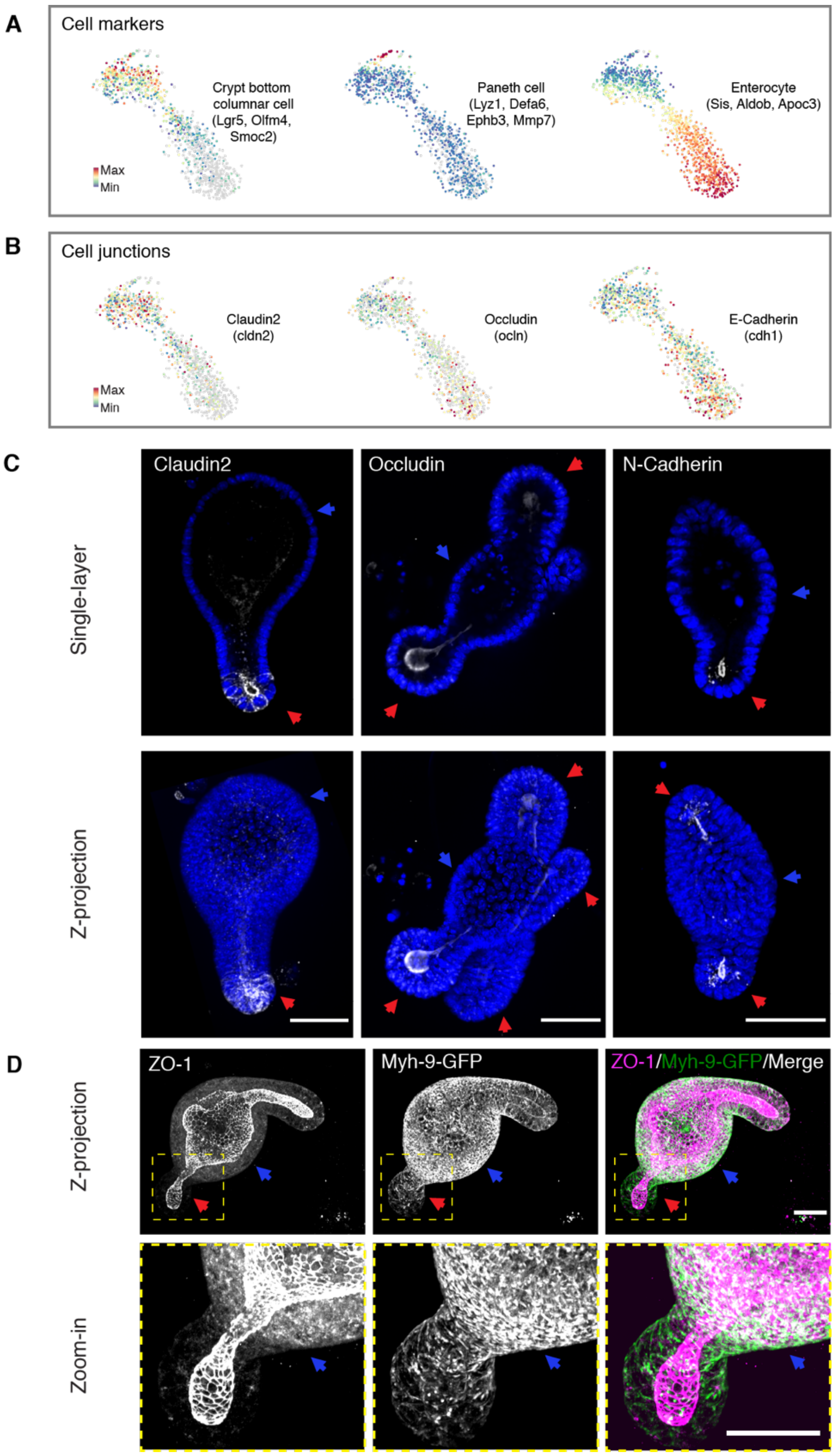


**Supplementary Figure 6: Region-specific expression of cell junctions. A-B**. Single-cell RNA (scRNA) analysis of budded organoid**.** TSNE-based visualization of single-cell degree of expression of marker genes for cell types (stem cells/Crypt columnar cells, Paneth cells and enterocytes) (A), and cell junctions (Claudin2, Occludin and E-Cadherin) (B). **C**. Middle single-layer section and maximum z-projection of the Claudin2, Occludin and N-Cadherin staining (white) with DAPI staining for cell nuclei (blue). **D**. Co-localization of ZO-1 and Myh-9-GFP in cell basolateral domains in villus region. From left to right: Z-projection (upper pannels) and zoom-in region (lower panels) of ZO-1 (white), Myh-9-GFP (white), and merged (merged signal in white) ZO-1 (magenta) and Myh-9-GFP (green). In C-D, red arrows indicate crypt regions, blue arrows indicate the villus regions. Scale bars, 50 μm.

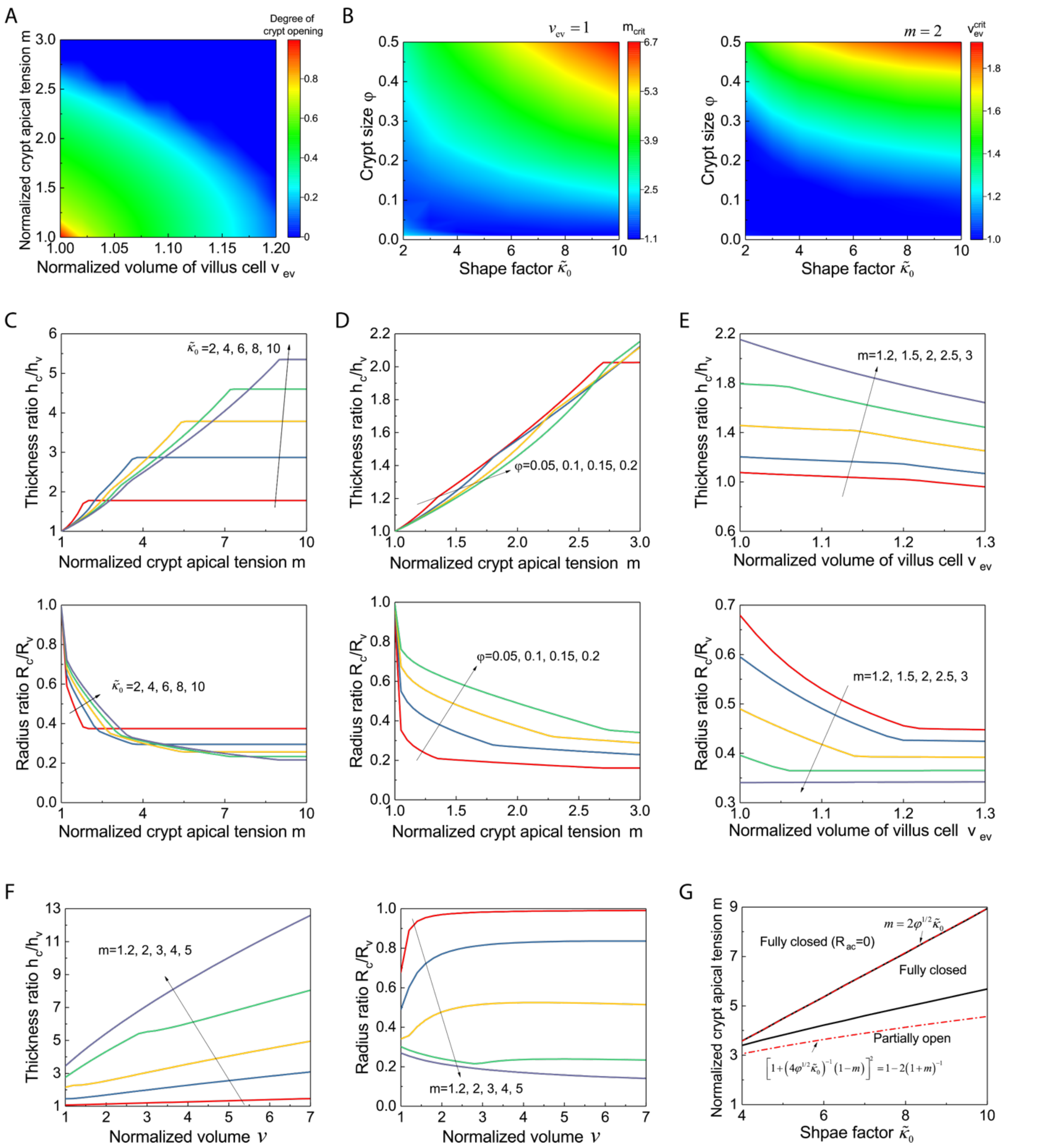


**Supplementary Figure 7. Sensitivity analysis of model results as a function of crypt apical tension and osmotic changes. A**. Phase diagram of crypt morphologies ($\tilde{\kappa}_0 = 6$, $\varphi = 0.2$) as a function of normalized crypt apical tension $m$ and villus cell volume $v_{\mathrm{ev}}$. **B**. Influence of crypt size $\varphi$ and shape factor $\tilde{\kappa}_0$ on critical values of $m$ (with $v_{\mathrm{ev}} = 1$) and $v_{\mathrm{ev}}$ (with $m = 2$) for organoid budding. **C-E**. Dependence of thickness ratio $h_{\mathrm{c}}/h_{\mathrm{v}}$ and radius ratio $R_{\mathrm{c}}/R_{\mathrm{v}}$ on normalized crypt apical tension $m$ ($v_{\mathrm{ev}} = 1$), with varied $\tilde{\kappa}_0$ ($\varphi = 0.2$) (panel C) and $\varphi$ ($\tilde{\kappa}_0 = 6$) (panel D), and on villus cell volume $v_{\mathrm{ev}}$ (panel E), with varied $m$ ($\tilde{\kappa}_0 = 6, \varphi = 0.2$). **F**. Evolution of $h_{\mathrm{c}}/h_{\mathrm{v}}$ and $R_{\mathrm{c}}/R_{\mathrm{v}}$

during organoid inflation with varied $m$ ($\tilde{\kappa}_0 = 6, \varphi = 0.2$, $v_{\text{ev}} = 1$). **G**. Phase diagram of crypt morphologies upon infinite volume expansion (normalized volume $v = 10^8$) (showing three possible phases: fully closed, partially opening, or fully closed with vanishing apical surface), with varying $\tilde{\kappa}_0$ and $m$ ($\varphi = 0.2$).

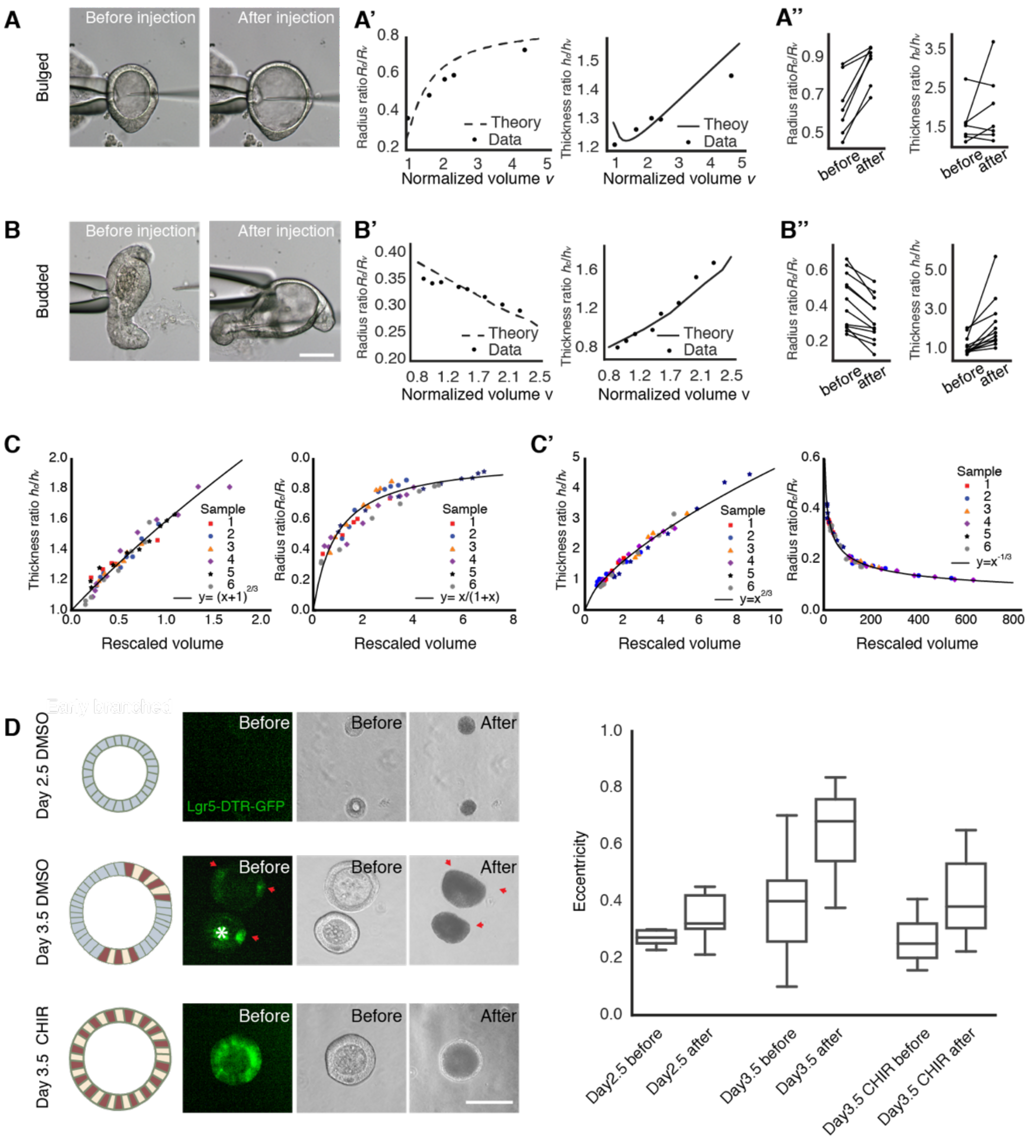

**Supplementary Figure 8: The impact of lumen volume on organoid morphology. A-B''**. Fitting for the lumen inflation experiments with the theoretical model. Images, microinjection of bulged (A) and budded(B) organoids for lumen inflation. Scale bars, 50 µm. Fittings (A' and B'), fitting experimental measurements (black dots, data) from the representative samples with its predictive model based on epithelial thickness (cell height) ratio (black lines, $h_c/h_v$), organoid radius ratio (black dashed lines, $R_c/R_v$) and lumen volume (v) change. Plots (A'' and B''), measurements of epithelial thickness ratio and radius ratio in bulged (A'', n = 7) and budded (B'', n = 12) organoids. Paired Student's *t* test for bulged $R_c/R_v$ ($p < 0.05$), bulged $h_c/h_v$ ($p = 0.37$), budded $R_c/R_v$ ($p < 10^{-2}$) and budded $h_c/h_v$ ($p < 10^{-2}$). **C-C'**. Evolution of morphometric parameters of bulged (C) or budded (C') organoids during volume inflation, induced by PGE treatment and microinjection (each with three samples), can be collapsed via $R_c/R_v$ vs volume and $h_c/h_v$ vs volume. **D**. Osmotic deflation facilitates the organoid shape transformation in organoids with different spontaneous curvatures. Cartoon images demonstrate Day2.5 organoid without tissue differentiation (upper panel), Day3.5 bulged organoid that has crypt and villus regions (middle panel), and Day3.5 organoid treated with 3 µM CHIR without obvious tissue difference (lower panel). Colours in the cartoon, red for Paneth cell, yellow for stem cell and blue for cell in villus tissue. Organoid images from left to right, fluorescent images of organoid expressing Lgr-5-DTR-GFP, bright filed image of organoid before, and after osmotic shock with 250 mM NaCl. Red arrows indicate the crypt region in Day3.5 organoids, white asterisk indicates the autofluorescence of dead cells in the lumen. Scale bar, 50 µm. Plot, box plot quantification on the eccentricity of organoids before and after osmotic shock. Paired Student's *t* test for organoids at Day2.5 ($n = 8$, $p = 0.03$), Day3.5 ($n = 22$, $p < 10^{-6}$) and Day3.5 CHIR ($n = 13$, $p = 0.002$). Box plot elements show quartiles, and whiskers denote 1.5× the interquartile range.

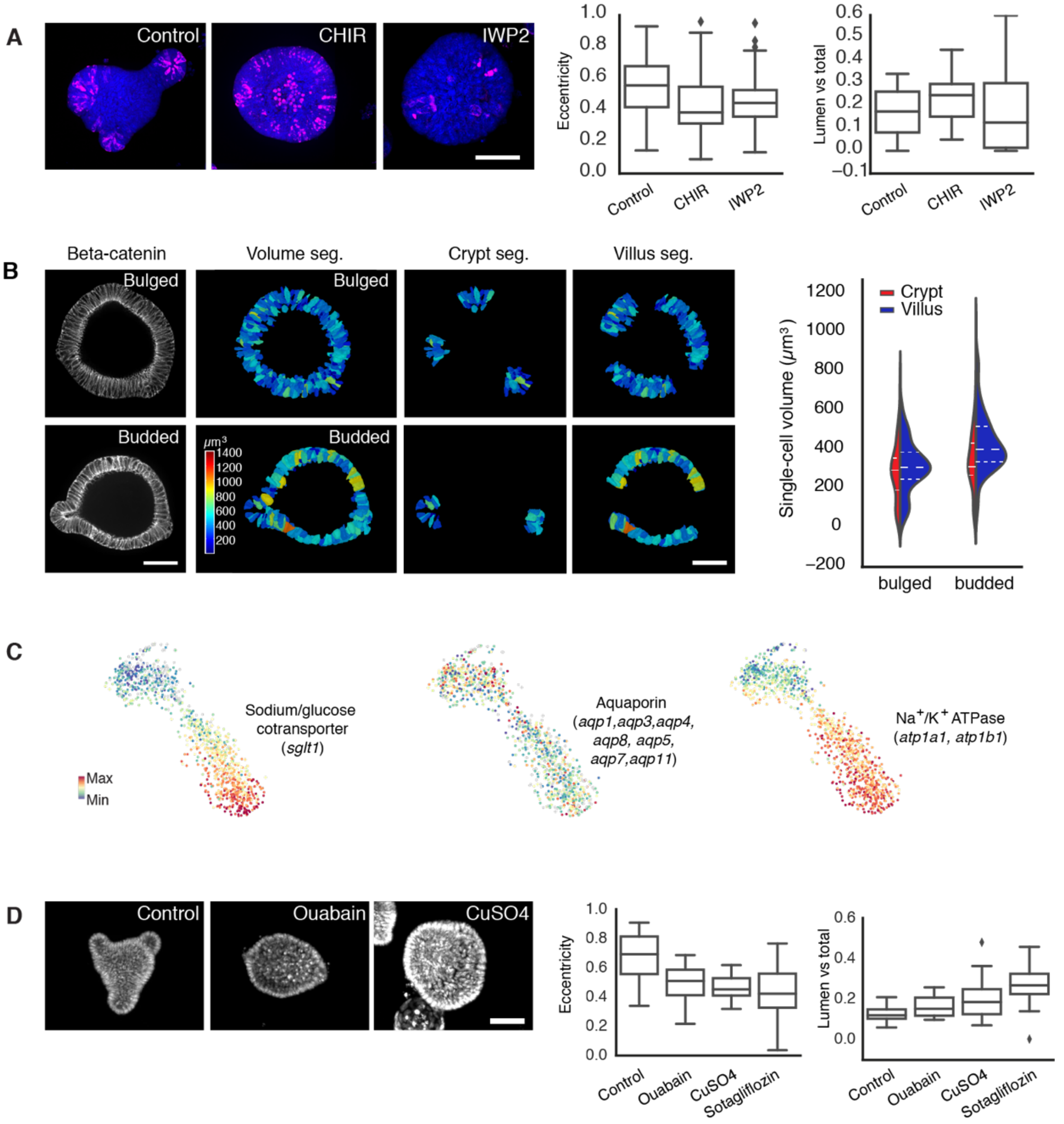


**Supplementary Figure 9: Regulation of lumen volume by enterocytes and the membrane transporters. A.** Enterocyte is responsible for the lumen volume reduction. Images, representative images of 5 µM DMSO-treated control, 3 µM CHIR- and 2 µM IWP2- treated organoids with Lyz staining for Paneth cells (magenta) and DAPI staining for cell nuclei (Blue). Scale bar, 50 µm. Plots, box plots for quantification of eccentricity and lumen ratio of organoids in control (n = 91 from 3 replications), CHIR (n = 140 from 3 replications), and IWP2 (n = 82 from 3 replications) treated samples. Two-tailed t-test for eccentricity of CHIR (to control, $p < 10^{-3}$), IWP2 (to control, $p < 10^{-3}$) and lumen ratio of CHIR (to control, $p < 10^{-3}$) and IWP2 (to control, $p = 0.75$). Box plot elements show quartiles, and whiskers denote 1.5× the interquartile range. **B**. Increased single-cell volume in villus tissue during budding. Images, Segmentation (Seg.) of single-cell volume based on the immunostaining of β-Catenin (left panels, white) in bulged and early-budded organoids. Plot, violin plot for the quantification of single-cell volume. Two-tailed t-test for the crypt (n= 109 from 3 organoid) and villus (n = 351 from 3 organoids) in bulged organoid ($p = 0.0996$), crypt (n= 41 from 2 organoids) and villus (n = 161 from 2 organoids) in budded organoids ($p < 10^{-2}$), and bulged villus and budded villus ($p < 10^{-17}$). Violin plot lines denote quartile for each group. **C**. Single-cell RNA analysis indicates the expression of membrane transporters. From left to right, TSNE-based visualizations indicate *sglt1* is enriched in enterocytes (Fig. S6A) in villus region, mRNAs of aquaporins are distributed broadly in cells in crypt and villus regions, *atp1a1* and *atp1b1* are enriched in enterocytes in villus region. **D**. Inhibition of the Aquaporins and $Na^+/K^+$ ATPase prevent the reduction of lumen volume. Images, Day4 organoid of 5 μM DMSO-treated control, 500 μM Ouabain treated sample with $Na^+/K^+$ ATPase inhibition and 12.5μM $CuSO4$ treated sample with Aquaporin inhibition. Scale bar, 20 μm. Plots: Box plots for the quantification on the eccentricity and lumen ratio (lumen volume versus total volume) of control (n = 80 from three replication), Ouabain treated sample (n= 57 from three replication), CuSO4 treated sample (n= 54 from three replication) and 5 μM Sotagliflozin treated sample (inhibition of SGLT-1, n= 46 from three replication). Two-tailed t-test for eccentricity of Ouabain (to control, $p < 10^{-3}$), CuSO4 (to control, $p < 10^{-4}$), Sotagliflozin (to control, $p < 10^{-5}$), and lumen ratio of Ouabain ( to control, $p = 0.012$) , CuSO4 ( to control, $p < 10^{-2}$) and Sotagliflozin (to control, $p < 10^{-8}$). Box plot elements show quartiles, and whiskers denote 1.5× the interquartile range.

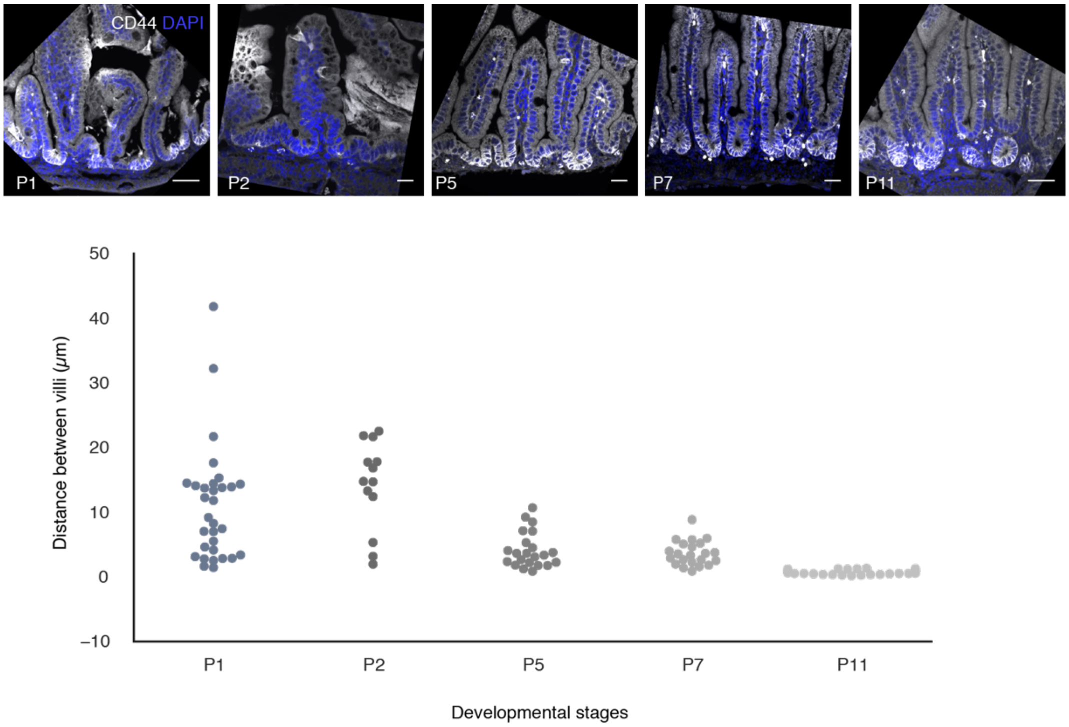


**Supplementary Figure 10: Reduced space between villi *in vivo*.** Upper panel: representative images of sections stained with CD44 for crypt region in white and DAPI for cell nuclei in blue in the mouse small intestine at P1, P2, P5, P7 and P11. Lower panel: Swarm plot for the quantification of distance between villi, which is measured by average 30 equally distributed distances within in each select region as the method applied to measure epithelial thickness indicated in Fig.1A. Two-tailed t-test for groups of P1 (n = 43 different regions) and P2 (n = 12 different regions) ($p = 0.01$), P2 and P5 (n = 22 different regions) ($p < 10^{-6}$), P5 and P7 (n = 21 different regions) ($p = 0.44$), P7 and P11(n = 29 different regions) ($p < 10^{-9}$). Scale bars, 50 µm.

**Table S1. Antibodies used in organoid immunostaining.**

| Primary Antibody | Company | Catalog NO. | Concentration |
|---|---|---|---|
| anti-ZO-1 | Thermo Fischer Scientific | 33910 | 1:200 |
| anti-SGLT-1 | Abcam | Ab14685 | 1:200 |
| anti-CD44v6 | BioRad | MCA1967 | 1:300 |
| anti-MLC-S1 | Abcam | Ab157747 | 1:200 |
| anti-Occludin | Thermo Fischer Scientific | 711500 | 1:200 |
| anti-N-Cadherin | BD transduction laboratories | 610920 | 1:200 |
| anti-Claudin2 | Thermo Fischer Scientific | 325600 | 1:200 |
| anti-Lysozyme | Abcam | Ab36362 | 1:500 |
| anti-Beta-Catenin | BD transduction laboratories | 610154 | 1:200 |
| **Secondary Antibody** | **Company** | **Catalog NO.** | **Concentration** |
| Alexa488 goat anti-rabbit IgG (H+L) | Thermo Fischer Scientific | B40943 | 1:2000 |
| Alexa568 goat anti-rat IgG (H+L) | Invitrogen | A11077 | 1:2000 |
| Alexa488 donkey anti-mouse IgG (H+L) | Thermo Fischer Scientific | A21202 | 1:2000 |

**Movie S1. Representative light-sheet time-lapse recording of crypt morphogenesis.**

A full stack of an organoid expressing LifeAct-GFP is acquired every 10 minutes for 16:40 hours from Day3 till budding. Left panel, single plane intersecting the middle of the organoid. Middle panel, maximum z-projection. Right panel, dynamic plots of organoid eccentricity, normalized lumen and total volume over time. Experiments were repeated at least five times. Scale bar, 20 µm.

**Movie S2. Representative light-sheet time-lapse recording of Myh-9-GFP expression in organoid during crypt morphogenesis.**

A full stack of an organoid expressing Myh-9-GFP is acquired every 10 minutes for 16:40 hours from Day3 till budding. Left panel, single plane intersecting the middle of the organoid. Right panel, maximum z-projection. Experiments were repeated at least three times. Scale bar, 50 µm.

**Movie S3. Representative light-sheet time-lapse recording of budded organoid treated with Blebbistatin.**

A full stack of budded organoid expressing LifeAct-GFP is acquired every 10 minutes from Day4, treated with 7.5 µM Blebbistatin for 18 hours and washed out, cultured and recorded for another 7 hours. Experiments were repeated at least two times. Scale bar, 50 µm.

**Movie S4. Representative light-sheet time-lapse recording of budded Organoid treated with Aphidicolin.**

A full stack of budded organoid expressing LifeAct-GFP is acquired every 10 minutes from Day4, The organoid was treated with 0.6 µM Aphidicolin for 18 hours, then cultured and recorded for another 7 hours after Aphidicolin washout. Experiments were repeated at least three times. Scale bar, 50 µm.

**Movie S5. Representative time-lapse recording of PGE induced inflation.**

Full stacks of bulged and budded organoids expressing LifeAct-GFP are acquired every 3 min immediately after treated with 0.5 µM PGE for less than 30 min. Left panel, maximum z-projection. Right panel, single plane intersecting the middle of the organoid. Experiments were repeated at least three times. Scale bar, 20 µm.

**Movie S6. Representative time-lapse recording of microinjection into organoid lumen for inflation.**

Bright-field time-lapse recoding of bulged and budded organoids under microinjection. Experiments were repeated at least three times. Scale bar, 50 µm.

**Movie S7. Representative time-lapse recording of Osmotic deflation.**

Lgr-5-DTR-GFP expression and Bright-field time-lapse recoding of Day2.5 organoids (left panel), Day3.5 organoids treated with 5 µM DMSO (middle panel) and Day3.5 treated with 3 µM CHIR (left panel), under osmotic shock by 250 mM NaCl. Experiments were repeated at least three times.

Scale bar, 50 µm.

**Movie S8. Representative light-sheet time-lapse recording of enterocyst from Day3.**

A full stack of enterocyst expressing LifeAct-GFP is acquired every 10 minutes from Day3 for 16:40 hours. Left panel, single plane intersecting the middle of the organoid. Middle panel, maximum z-projection. Right panel, dynamic plots of organoid eccentricity, normalized lumen and total volume over time. Experiments were repeated at least five times. Scale bar, 20 µm.

**Movie S9. Representative light-sheet time-lapse recording of organoid treated with CHIR from Day3.**

A full stack of an organoid expressing LifeAct-GFP is acquired every 10 minutes from Day3 for 16:40 hours while being treated with 3 µM CHIR from Day 3. Left panel, single plane intersecting the middle of the organoid. Middle panel, maximum z-projection. Right panel, dynamic plots of organoid eccentricity, normalized lumen and total volume over time. Experiments were repeated at least two times. Scale bar, 20 µm.
